# A Sequential Binding Mechanism for 5’ Splice Site Recognition and Modulation for the Human U1 snRNP

**DOI:** 10.1101/2024.04.18.590139

**Authors:** David S. White, Bryan M. Dunyak, Frédéric H. Vaillancourt, Aaron A. Hoskins

**Author notes:** Corresponding Author: Aaron A Hoskins.

## Abstract

Splice site recognition is essential for defining the transcriptome. Drugs like risdiplam and branaplam change how U1 snRNP recognizes particular 5’ splice sites (5’SS) and promote U1 snRNP binding and splicing at these locations. Despite the therapeutic potential of 5’SS modulators, the complexity of their interactions and snRNP substrates have precluded defining a mechanism for 5’SS modulation. We have determined a sequential binding mechanism for modulation of -1A bulged 5’SS by branaplam using a combination of ensemble kinetic measurements and colocalization single molecule spectroscopy (CoSMoS). Our mechanism establishes that U1-C protein binds reversibly to U1 snRNP, and branaplam binds to the U1 snRNP/U1-C complex only after it has engaged a -1A bulged 5’SS. Obligate orders of binding and unbinding explain how reversible branaplam interactions cause formation of long-lived U1 snRNP/5’SS complexes. Branaplam is a ribonucleoprotein, not RNA duplex alone, targeting drug whose action depends on fundamental properties of 5’SS recognition.

## INTRODUCTION

Splice site modulators are synthetic molecules, either oligonucleotides (oligos) or small molecules, that change the splicing outcome of a given pre-mRNA by either enhancing or preventing the use of a given splice site (the boundaries between exons and introns in a pre-mRNA)^1,2^. To date, the most successful of these molecules have been nusinersen (an oligo) and risdiplam (a small molecule) that are both approved treatments for spinal muscular atrophy (SMA)^3,4^. SMA is caused by mutations in the *SMN1* gene that prevent production of functional protein^5^. A nearly identical gene, *SMN2*, is also present that can be used to produce the same protein but is normally inactive due to exclusion of exon 7 from the mRNA transcript by alternative splicing^6–8^. Both nusinersen and risdiplam promote exon 7 inclusion to allow for translation of the functional protein in SMA patients^4^. However, the mechanisms by which these two drugs function are quite different. Nusinersen targets an intronic splicing silencer element downstream of exon 7 that naturally represses exon 7 splicing^9^. On the other hand, risdiplam and related molecules such as branaplam are believed to restore exon 7 splicing by increasing the interaction between the U1 small nuclear ribonucleoprotein (snRNP) splicing factor and exon 7’s weak 5’ splice site (SS)^10–13^.

Remarkably, risdiplam and branaplam do not modulate splicing at all 5’SS but instead exhibit a high degree of selectivity^12–15^. The origins of this specificity or differences in which sites are impacted between risdiplam and branaplam are not well-understood^14^. The *SMN2* exon 7 5’SS is atypical relative to more common sites in that it contains a bulged nucleotide at the -1 position (-1 represents the last nucleotide of the exon, +1 represents the first nucleotide of the intron). The bulged nucleotide (an adenosine) is predicted to be unpaired and flipped out of the RNA duplex formed between the 5’SS and the small nuclear RNA (snRNA) component of the U1 snRNP^10^. Biochemical and *in vivo* data have suggested that risdiplam and branaplam act specifically at these types of 5’SS and function by a “bulge repair” mechanism to convert the bulged duplex into something that more closely resembles a standard interaction between a 5’SS and U1 snRNA^10,14,16^. Since perturbations in U1 snRNP/5’SS binding can lead to changes in alternative splicing^17,18^, it is thought that these drugs enhance U1 snRNP affinity for the *SMN2* exon 7 5’SS, which in turn promotes spliceosome assembly, splicing modulation, and exon inclusion. Thus, these drugs appear to modulate the SS recognition process to ultimately alter the nucleotide sequences of spliced mRNA products.

The 5’SS is recognized by multiple factors during spliceosome assembly including the U1 and U6 snRNAs. U1 and U6 base pair sequentially to the 5’SS with U1 snRNP associating during the earliest stages of spliceosome assembly before being displaced by the U6 snRNA just prior to spliceosome activation, when the active site for splicing is formed^19^. The U1 snRNP is composed of the U1 snRNA and 10 protein factors. The snRNA region that base pairs with the 5’SS is found at the very 5’ end and can form up to 11 contiguous base pairs, flanking both sides of the exon/intron boundary^17^. In humans, most 5’SS have fewer potential base pairing interactions with U1 snRNA and are highly sequence diverse—more than 9,000 different 5’SS sequences are functional in humans, some of which contain atypical base pairing registers or bulged nucleotides as in SMN2^17^. While the GU at the +1 and +2 positions are nearly invariant due to the constraints placed on this location by splicing chemistry, every possible nucleotide can be accommodated at every other position^17,20^. In both yeast and humans, the zinc finger domain of the U1-C protein binds the backbone of the 5’SS/snRNA duplex near the “GU” dinucleotide and has been postulated to fine-tune U1 snRNP interactions with RNA^21^. While yeast U1-C is an obligate snRNP component^22^, human U1-C is smaller and readily dissociates from purified human U1 snRNPs at room temperature^23^.

Quantitative models for 5’SS recognition by human U1 snRNP could reveal how the splicing factor discriminates between different RNA sequences as well as how this process is modulated by U1-C or drugs such as branaplam. A thermodynamic model has recently been proposed for explaining the *in vivo* effects of splice site modulating drugs^14^; however, there are no detailed kinetic data yet available for splice site recognition by human factors. Obtaining a kinetic mechanism of 5’SS recognition in a biochemically-defined system is essential for understanding and predictive modeling of the splicing decisions that give rise to the cellular transcriptome as well as understanding the mechanism of action of splicing modulator drugs^14^. A kinetic description of this process is particularly important since splicing, like many events in gene expression, does not occur under equilibrium conditions and is likely limited to a “window of opportunity”^24^. The window of opportunity is defined, in part, by how quickly U1 can associate with a 5’SS and how long it may remain bound in order to promote spliceosome assembly before the RNA is degraded, exported, or a competing 5’SS is utilized. Splice site modulation by drugs such as branaplam is likely also restricted to the same, or related, window of opportunity.

To elucidate 5’SS recognition and modulation in humans, we reconstituted a model human U1 snRNP and assayed its interactions with RNA oligos in the presence and absence of branaplam. A combination of surface plasmon resonance (SPR), microscale thermophoresis (MST) and colocalization single molecule spectroscopy (CoSMoS) assays reveals how 5’SS containing a bulged adenosine at the -1 position (-1A) are recognized and modulated by drugs working collaboratively with protein splicing factors. Branaplam reversibly binds to the U1 snRNP/5’SS complex, and drug modulation of this complex is strictly dependent on reversible binding of U1-C. U1-C in turn can only bind to the snRNP if the 5’SS has not yet been engaged. Thus, our sequential binding mechanism predicts that 5’SS modulation by branaplam depends on an ordered series of events: U1-C binds to U1 snRNP, this complex then binds to the 5’SS, and finally branaplam binds to the U1 snRNP/U1-C/5’SS ternary complex. This mechanism reveals how a reversibly binding splicing modulator can elicit formation of long-lived U1 snRNP/5’SS interactions as well as fundamental features of human 5’SS recognition.

## RESULTS

### A -1A bulge in the 5’SS induces dynamic RNA binding to U1 snRNP

To measure the effects of branaplam on U1 snRNP/5’SS interactions, we reconstituted a minimal U1 snRNP based upon prior work from the Nagai laboratory (**Fig. 1A**, **S1**)^21^. Our U1 snRNP complex consisted of seven Sm proteins, the N-terminal region of U1-70K, the zinc finger region of U1-C, and a truncated U1 snRNA. It has previously been shown that the U1-A protein and snRNA stem loops I and II (which contains the binding site for U1-A) are not essential for splicing^25^. As in prior work by Nagai and coworkers, several Sm proteins were also truncated to remove unstructured domains and the U1-70K fragment was fused to SmD1 (see Methods). The sequence of our commercially synthesized U1 snRNA was further altered in the shortened stem loop 1 region to prevent dimerization (as opposed to the construct used by Nagai and coworkers for crystallization) and modified with 2’-*O*-methyls at the +1 A and +2 U positions and pseudouridines at +5 and +6 as found in the native human U1 snRNA. The snRNA was designed without a 5’ cap, which is also not believed to be required for SS recognition and splicing^25^. The complex was assembled in solution from purified components (including U1-C, unless otherwise noted) following established methods and purified via size exclusion chromatography^21^.

**Figure 1:**
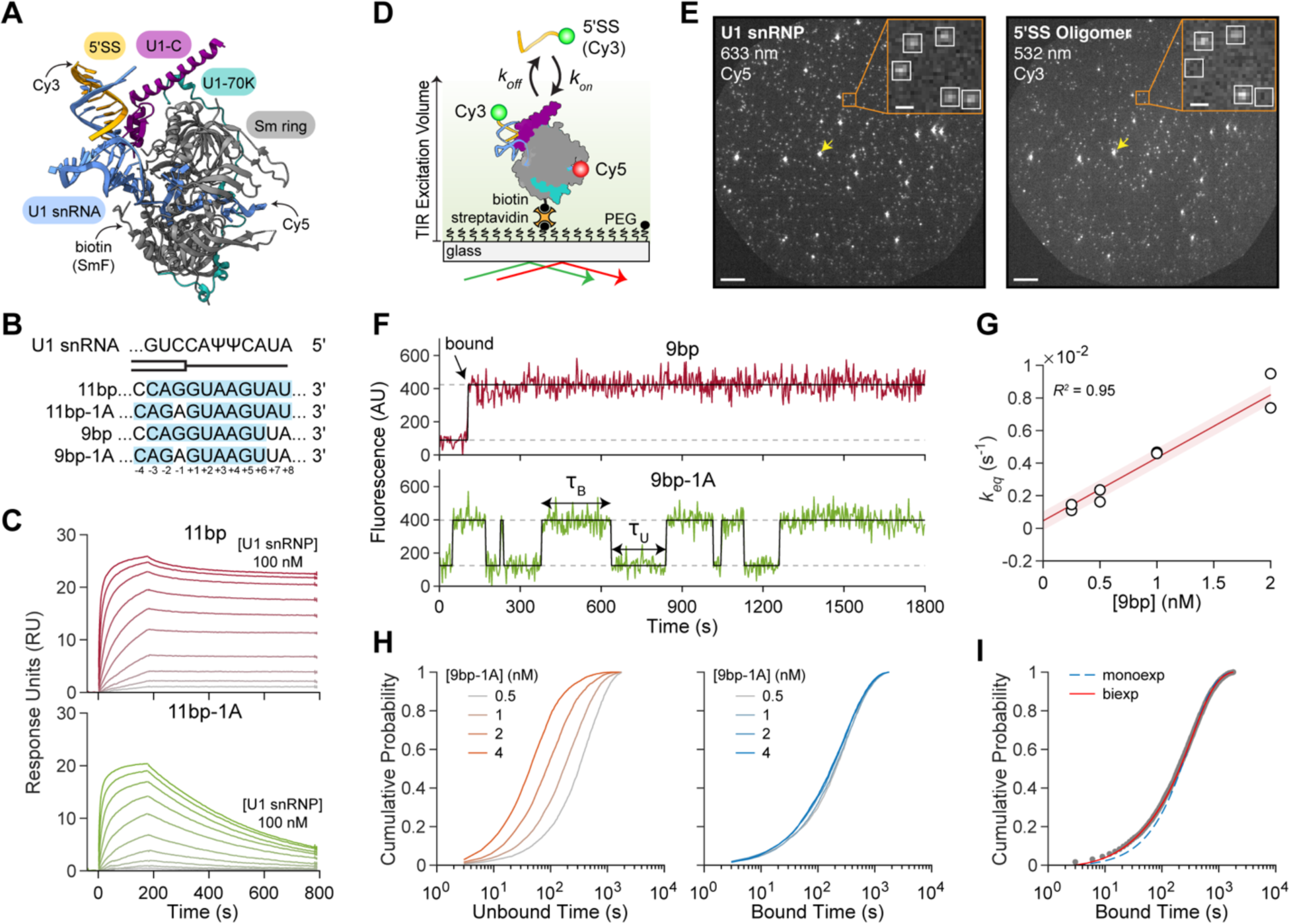
A -1A bulge induces dynamic RNA binding to U1 snRNP. (**A**) Crystal structure of minimal U1 snRNP (PDB: 4PJO). Arrows indicate relative placement of modifications for single molecule measurements. (**B**) RNA oligos containing 5’SS. Shaded region indicates predicted base pairing interactions with the U1 snRNA (shown at the top, above a schematic of the exon (box)/intron (line) junction). (**C**) SPR sensorgrams showing the association and dissociation of surface-tethered (top) 11bp and (bottom) 11bp-1A RNAs at various U1 snRNP concentrations (0.02 to 100 nM). (**D**) Cartoon schematic of the two-color CoSMoS assay for monitoring U1/RNA interactions. (**E**) Fields of view under 633 nm (left, immobilized U1 snRNP molecules) and 532 nm (right, interacting RNA oligos) excitation (scale bar = 20 µm). Inset highlights colocalization (scale bar 1 µm). Fluorescent beads were included as fiducial makers (yellow arrow). Images are rendered by averaging three consecutive images and applying uniform brightness and contrast values. (**F**) Fluorescence time series showing the binding of 9bp (top) and 9bp-1A (bottom) to surface tethered U1 snRNP (0.33 frames/s, Hz). Black lines indicate idealized fit. (**G**) Linear regression of *k_eq_* vales on 9bp concentration, given by *k_eq_* = *k_on_*([9bp]) + *k_off_*. Shaded region is the 95% confidence interval. Here, the *k_on_* value is fixed from maximum likelihood estimations of unbound dwell times (see Figure 1F) at *k_on_* = 3.9 x 10^6^ M^-1^s^-1^, resulting in a *k_off_* = 4.6 ± 2.9 x 10^-4^ s^-1^. (**H**) Cumulative probability distributions of (left) unbound dwell times (0.5 nM = 2670; 1nM, *N* = 5154; 2 nM, *N* = 3788; 4 nM, *N* = 4373) and (right) bound dwell times (0.5 nM = 2380; 1nM, *N* = 5216; 2 nM, *N* = 4093; 4 nM, *N* = 4895) across range of 9bp-1A RNA concentrations. (**I**) Cumulative probability distribution of bound dwell times of the 9bp-1A RNA at 1 nM (grey, *N* = 5216) overlaid with MLE of single (blue dashed) and biexponential (red) distributions.

We first determined the effect of a -1A substitution in the 5’SS on U1 snRNP/5’SS duplex formation using a fully complementary 5’SS oligo (11bp) and a -1A bulged variant (11bp-1A) (**Fig. 1B**) by surface plasmon resonance (SPR). In these assays, the 5’SS oligo is tethered to the surface via a 3’ biotin and U1 snRNP is added in solution at various concentrations (1.23-100 nM). As expected, the fully complementary 11bp sequence binds the U1 snRNP very tightly (**Fig. 1C top,** *K_D_* = 2.03 ± 0.08 x 10^-10^) while the introduction of the -1A reduces the affinity by an order of magnitude (*K_D_* = 2.70 ± 0.01 x 10^-9^ M). These values stem from global fitting to a single-site binding model consisting of a single *k_on_* and *k_off_*; however, this simple model does not sufficiently describe the sensorgrams of 11bp or 11bp-1A (**Fig. S2**). To circumvent the challenge of identifying unique kinetic parameters that best describe ensemble data in nonlinear models^26^, we used single-molecule co-localization spectroscopy (CoSMoS) to observe U1 snRNP/5’SS interactions.

For CoSMoS experiments, we generated a modified minimal U1 snRNP similar to that used for the SPR assays except that it also contained a biotin moiety on SmF for surface tethering and a 3’-Cy5 dye on the U1 snRNA (**Fig. 1A, S1B**). Due to potential interference from U1-C dissociating from the U1 snRNP^21,23^ under the dilute conditions required for single molecule immobilization, we reconstituted our U1 snRNP particle without U1-C. The U1-C zinc-finger domain was then separately purified and added directly into solution as required (typically at 100 nM unless otherwise noted; U1-C binding was also analyzed in depth as described subsequently).

The fluorescently-labeled and biotinylated U1 snRNP particle was immobilized with streptavidin onto a passivated and biotinylated glass slide for CoSMoS measurements (**Fig. 1D**). An RNA oligo containing a 3’-Cy3 dye was added to the solution for direct visualization of U1 snRNP/5’SS binding dynamics. Binding events were monitored using either sequential or alternating laser excitation of 532nm (5’SS, Cy3) and 633nm (U1 snRNP, Cy5). Co-localized spots were detected in each channel following channel alignment, and the fluorescence time series at each spot were idealized using unsupervised statistical methods to delineate bound and unbound events for each U1 snRNP molecule (Methods, **Fig. 1E**). For all 5’SS oligos we investigated, we flanked the 5’ and 3’ ends with an ‘ACA’ motif outside of the 11 nucleotides corresponding to the 5’SS. The secondary structure of each 5’SS/U1 snRNA duplex was predicted using the RNAstructure Web Server^27^ to ensure the minimum free energy structure matched our expectations (**Fig. S3**).

We first replicated the effect of a -1A bulge on the U1:5’SS interaction observed by SPR at the single molecule level. Considering the tight binding of the 11bp-1A RNA without branaplam, we opted to use a 9bp-1A oligo for single molecule experiments. We hypothesized the reduced number of base pairs would increase the *k_off_* and minimize the impact of photobleaching over extended observation times. Mismatches were incorporated at +7 and +8 of the 5’SS, as these positions are not well conserved in humans^17^. Similar to our SPR data, we see exceptionally tight binding of the 9bp compliment (**Fig. 1F, S4**). No binding was observed at 10 nM of a fully mismatched oligo at the same frame rate (observed fraction bound < 0.002 across 1297 U1 snRNP molecules compared to an observed fraction bound of 0.77 across 1271 molecules at 1nM of 9bp **Fig. S5**). This demonstrates that non-specific binding did not meaningfully impact our analysis under the experimental conditions.

Given that the dissociation of the 9bp 5’SS RNA was very slow, we implemented a non-equilibrium imaging scheme whereby the cumulative binding of the 9bp RNA was monitored over time immediately following its addition to solution^28^. This measurement was repeated across various concentrations of RNA from 0.25-2 nM (0.25 nM, *N* = 1204; 0.5 nM, *N* = 1283; 1 nM, *N* = 1271; 2 nM, *N* = 1156, where *N* is the number of U1 molecules). We then determined the apparent association rate (*k_apparent_)* at each concentration using maximum likelihood estimation (MLE) of our unbound dwell time distributions. These parameters stem from estimations of a modified exponential probability density function that accounts for the sampling rate and observation time^29^. Next, we performed linear regression of *k_apparent_* values obtained at different 9bp 5’SS RNA concentrations to determine a *k_on_* = 3.9 ± 0.2 x 10^6^ M^-1^ s^-1^. The equilibration rate (*k_eq_*) at each RNA concentration was determined by fitting the observed fraction bound over time curves to a single exponential function (**Fig. S6**). Finally, we performed linear regression of *k_eq_* values as a function of 9bp RNA concentration (**Fig. 1G**). The slope was constrained at the *k_on_* value from MLE, resulting in a *k_off_* = 6.4 ± 2.3 x 10^-4^ s^-1^. Overall, we estimate a *K_D_* ≈ 1.2 x 10^-10^ M for a 9bp 5’SS RNA, which closely matches our SPR data of an 11b 5’SS RNA (*K_D_* = 2.03 x 10^-10^). The similarity of these values provides high confidence that neither photobleaching nor surface immobilization is significantly impacting our analysis. Together the SPR and CoSMoS data indicate that U1 snRNP binds highly complementary RNAs very tightly with a *K_D_* of ∼100 pM, and this affinity is primarily attributable to formation of very stable bound complexes with lifetimes of ∼27 min rather than rapid association kinetics. These data also indicate that additional base pairing interactions between the +7 and+8 positions of a 5’SS with the AU dinucleotide present at the 5’ end of the snRNA do not necessarily confer significant changes in the dissociation constant, *K_D_*.

At the single molecule level, the introduction of the -1A bulge (9bp-1A) into the 5’SS results in dynamic RNA binding to U1 snRNP (**Fig. 1F**, **S7**). Given the faster kinetics from the weaker binding, we were able to perform equilibrium measurements whereby imaging commenced after equilibrium was reached. Compared to the 9bp RNA, we see only a slight decrease in *k_on_* to 2.9 ± 0.2 x 10^6^ M^-1^ s^-1^, indicating the energetic penalty of -1A bulge stems from duplex stability and not recruitment. On average, the 9bp-1A 5’SS RNA exhibits a *k_off_* of 3.4 ± 0.2 x 10^-3^ s^-1^ and *K_D_* = 1.2 x 10^-9^ M. These results are also close to our SPR data for the 11bp-1A 5’SS RNA and confirm an order of magnitude reduction in kinetic stability from the -1A substitution.

We next analyzed the single molecule distributions of bound and unbound dwell times (**Fig. 1H, S7**). Bound dwell times across all 9bp-1A 5’SS RNA concentrations were poorly described by a single exponential distribution and instead required at least two exponential components (**Fig. 1I, S8**). MLE resulted in in two bound time constants (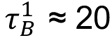 s and 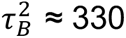 s), each with their own relative weight (A^1^ *≈* 0.12 and A^2^ *≈* 0.88). As expected for a bimolecular dissociation process, bound dwell time parameters are constant across -1A concentrations (**Fig. S8B**). We additionally performed MLE for unbound dwell times across -9bp-1A 5’SS RNA concentrations. Although subtle, we do see evidence for two time constants that becomes more apparent with increasing RNA concentration (**Fig. S8**). We conclude that introduction of a bulged-1A nucleotide reduces U1 snRNP affinity for a 5’SS by an order of magnitude; although, the interaction is still quite strong with a *K_D_* of 1-2 nM. The reduction in affinity is driven by a reduced stability of the U1 snRNP/5’SS complex whose average lifetime decreases from ∼27 to <5 min. This corresponds with a thermodynamic penalty due to the -1A bulge of ∼2 kcal/mol when binding to the U1 snRNP, similar to values reported for the impact of bulged nucleotides on the stabilities of RNA-only duplexes^30^.

### Branaplam modulates the binding of -1A bulged 5’SS RNA to U1 snRNP

We then investigated the modulation of the U1 snRNP/5’SS duplex by branaplam (**Fig. 2A, B**). At the ensemble level using SPR, we maintained a constant concentration of U1 snRNP in solution (10 nM) and varied the concentration of branaplam, revealing a branaplam-dependent slowing of U1 snRNP:11bp-1A 5’SS RNA dissociation (**Fig. 2C**). The dissociation phase of each sensorgram was fit to a single exponential decay function to determine *k_off_*. In the case of 11bp-1A RNA, the fitted *k_off_* values yielded an EC_50_ = 1.3 ± 0.1 µM for branaplam, and we see a 4-fold reduction in *k_off_* of the RNA due to inclusion of branaplam (2.35 ± 0.14 x 10^-3^ s^-1^ and 6.10 ± 0.21 x10^-4^ s^-1^ at 0 and 5 µM branaplam, respectively, **Fig. 2D**). Strikingly, under these high concentrations of branaplam, the *k_off_* of U1 snRNP:11bp-1A 5’SS RNA approaches the dissociation rate of the 11bp 5’SS RNA (*k_off_* = 2.11 ± 0.03 x 10^-4^ s^-1^). As expected, branaplam had no effect on the binding kinetics of the 11bp 5’SS RNA itself since this RNA lacks the -1A bulge (**Fig. 2D**).

**Figure 2:**
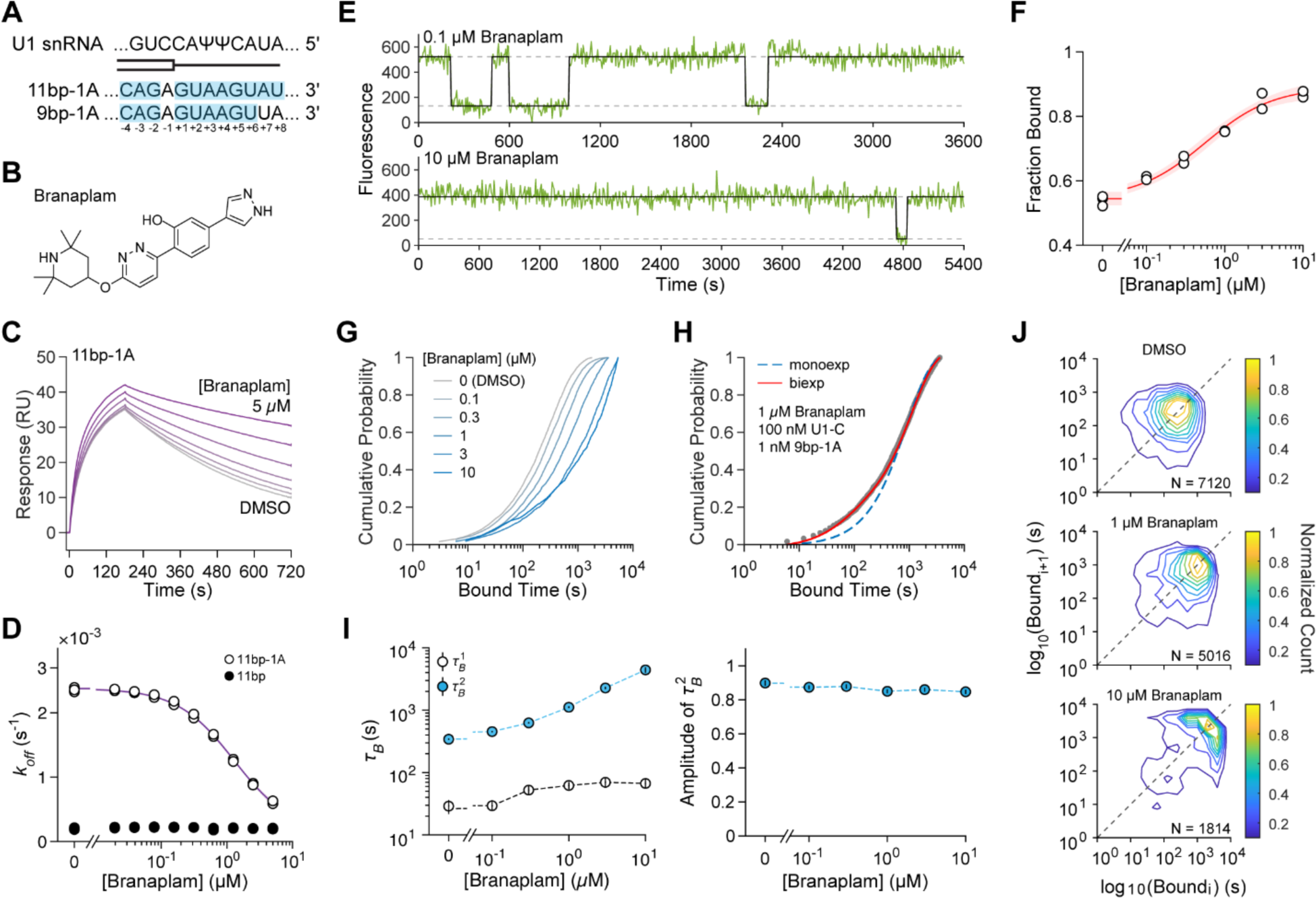
Branaplam stabilizes -1A bulged 5’SS RNA binding to U1 snRNP. (**A**) RNA oligos containing -1A bulged 5’SS. Shaded region indicates predicted base pairing interactions with the U1 snRNA (top). (**B**) Chemical structure of branaplam. (**C**) SPR showing the association and dissociation of 11bp-1A RNA across various concentrations of branaplam (0-5 µM) and at 10 nM U1 snRNP. (**D**) Dose response curve showing the fitted dissociation rates (*k_off_*) of response units vs branaplam concentration for 11bp-1A (white circles) and 11bp (black circles) RNAs. Data for 11bp-1A is overlaid with the fitted equation to determine EC_50_ value (1.3 ± 0.1 µM). (**E**) Fluorescence time series of 1 nM 9bp-1A RNA binding to immobilized U1 in the presence of 100 nM U1-C plus (top) DMSO (0.33 Hz) or (bottom) 10 µM branaplam (0.11 Hz) overlaid with idealizations (black lines) (**F**) Dose response curves of 9bp-1A RNA binding to U1 snRNP vs branaplam concentration. The red line and shading indicate the fit and 95% CI to the fitted equation to estimate an EC_50_ value (0.45 ± 0.17 µM). (**G**) Cumulative probability distribution of bound dwell times across branaplam concentrations (DMSO, *N* = 5216; 0.1 µM, *N* = 6234; 0.3 µM, *N* = 4811; 1 µM, *N* = 3640; 3 µM, *N* = 2211; 10 µM, *N* = 1503). (**H**) Cumulative probability distribution of bound dwell times at 1 nM 9bp-1A and 100 nM U1-C (grey circles, N = 3640) in presence of 1 µM branaplam overlaid with MLE of mono (blue dashed) and biexponential distributions (solid red). (**I**) Maximum likelihood estimates of (*left*) time constants (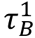 and 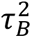) and (*right*) amplitude of 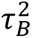 of a biexponential distribution for bound dwell times at 1 nM 9bp-1A RNA and 100 nM U1-C across branaplam concentrations (value ± standard error). (**J**) Contour plots showing the correlation between successive bound event durations (*i* and *i* + 1) within individual molecules. Dashed line is the identity line.

We then used CoSMoS assays to glean more detailed insights into how branaplam is altering 5’SS binding. CoSMoS experiments were conducted across branaplam concentrations between 0.1 to 10 µM, with 1 nM 9bp-1A 5’SS RNA and 100 nM U1-C in solution (**Fig. 2E, S9**). Consistent with our SPR data, we see the lifetimes of single binding events increase with increasing concentrations of branaplam (**Fig. 2G**). At our highest concentration of branaplam (10 µM), we observe some individual binding events persisting for over an hour. To capture these data with a minimal contribution from photobleaching, we decreased our sampling frame rate from 0.33 Hz in the absence of branaplam to 0.11 Hz at 10 µM branaplam. These experiments yielded an EC_50_ = 0.60 ± 0.12 µM for branaplam (**Fig. 2F**), similar to the EC_50_ determined by the *k_off_* rates from SPR for the 11bp-1A 5’SS RNA (1.3 ± 0.1 µM). We did not observe any significant effect of branaplam on the association rate of the 9bp-1A 5’SS RNA (**Fig. S10)**, consistent with branaplam specifically perturbing 5’SS RNA dissociation, but not association, rates.

We performed MLE of single and biexponential distributions of our bound dwell times across branaplam concentrations. As in the absence of branaplam, we find that two time constants are required to describe the distributions in all cases (**Fig. 2H**). We find that branaplam does not affect the relative amplitude of our two populations, but predominantly increases the slower bound time constant, 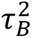, in a concentration dependent manner. The 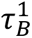 value and the relative amplitudes of the two time bound time constants are similar to the no branaplam conditions (**Fig. 2I**). Overall, an order of magnitude increase in 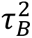 for the 9bp-1A 5’SS RNA is observed between the absence (342 ± 8 s) and presence of 10 µM branaplam (4426 ± 458 s). Together, the SPR and CoSMoS data indicate that branaplam does not facilitate -1A 5’SS RNA association and primarily functions to stabilize the U1 snRNP/-1A 5’SS RNA complex. However, only a subset of U1 snRNP/-1A 5’SS interactions are branaplam-sensitive.

We next tested whether or not this subset of branaplam-sensitive U1 snRNP/-1A 5’SS interactions arises from two different types of U1 snRNP/-1A 5’SS complexes on the surface (*e.g.,* +/- a particular, non-fluorescent subunit) or from reversible interconversion of a kinetically homogenous U1 snRNP/-1A 5’SS complex between two states (*e.g.,* a conformational change). We correlated the lifetimes of successive binding events at individual molecules. In the case of two different types of U1 snRNP/-1A 5’SS complexes, such analysis should yield at least two clusters of individual molecules showing distinct kinetic behaviors, whereas only a single cluster should emerge if a homogenous group of U1 snRNP/-1A 5’SS complexes were interconverting between two states^31^. This analysis indicates the presence of two distinct clusters, dependent on the branaplam concentration (**Fig. 2J**). At 10 µM branaplam, these clusters display unique average bound and unbound times for the -1A 5’SS RNA, and the molecules that more weakly and dynamically engage with the -1A 5’SS RNA account for 5% of the data (**Fig. S11**). This result suggests the presence of two types of U1 snRNP molecules on the surface, each with distinct kinetic properties. We were able to robustly detect the smaller population due to the analysis of many thousands of single molecule binding events and by avoiding ensemble averaging which would have obscured their presence.

### U1-C is required for 5’SS modulation by branaplam

We hypothesized that the source of the two distinct kinetic populations could be attributed to the presence or absence of U1-C given the reported lability of this factor and its contacts with the snRNA/5’SS duplex and site of the -1A bulge^21,23^. In this regard, the relative amplitudes from MLE of our bound time distributions in the presence of branaplam would presumably correspond to the fractions of U1 snRNP molecules without (the amplitudes of the short-lived parameters) and with U1-C (the amplitudes of the long-lived parameters).

In agreement with this hypothesis, we observed that withholding U1-C dramatically changed the binding kinetics (**Fig. 3A, B; S12**). In the absence of branaplam and U1-C, we see only weak binding at 1 nM 9bp-1A 5’SS RNA with an average bound lifetime of *τ_B_* = 65.2 ± 1.3 (*N* = 2381). This value matches the 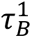 parameter we observed in the branaplam and no branaplam conditions. The addition of 10 µM branaplam in the absence of U1-C had no effect on duplex stability (τ*_off_* = 57.9 ± 0.9, *N* = 3806). These data are consistent with a model in which U1-C is required for branaplam binding and modulation of U1 snRNP/5’SS interactions.

**Figure 3.**
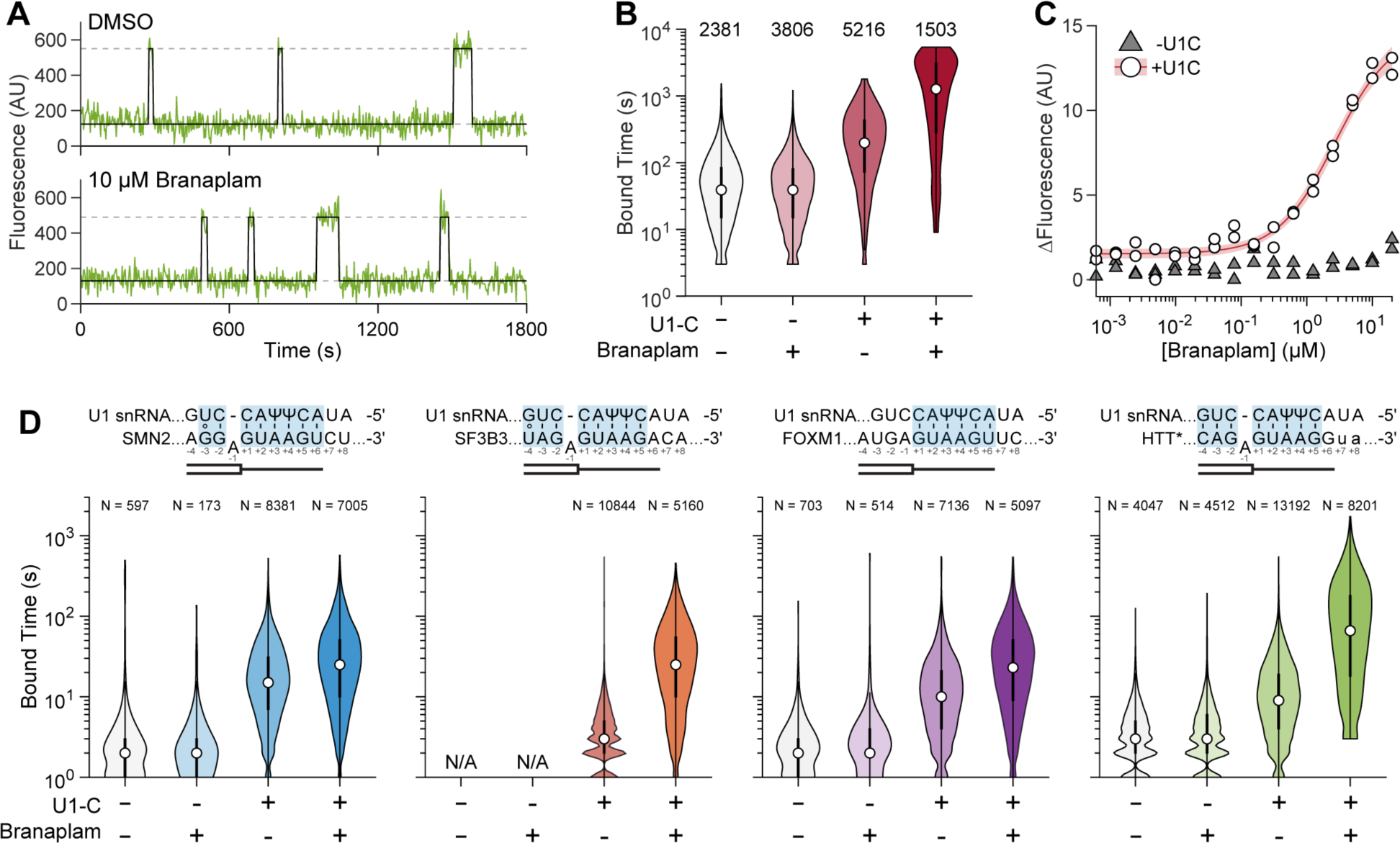
U1-C is required for branaplam modulation of -1A 5’SS RNA binding. (**A**) Fluorescence time series of 1 nM 9bp-1A binding in the absence of U1-C with (top) DSMO and (bottom) 10 µM branaplam overlaid with idealizations (black lines) (**B**) Violin plots of bound dwell time distributions at 1 nM 9bp-1A RNA with and without U1-C and/or branaplam. DMSO was included in the absence of branaplam. (**C**) Change in fluorescence due to branaplam binding to a duplex of 11bp-1A and U1snRNP in the absence (grey triangles) and presence (white circles) of U1-C by microscale thermophoresis (MST). Purple line indicates the fit and 95% CI to the fitted equation to estimate a *K_D_* value (2.69 ± 0.36 µM) in the presence of U1-C. (**D**) Bound dwell time distributions across different permutations of U1-C and branaplam concentrations in solution across indicated RNA oligo sequences (SMN2, FOXM1, SF3B3, HTT*). Highlighted nucleotides in the 5’SS sequences indicate predicted base pairs to the U1 snRNA. The lower case letters in HTT* indicate a +7G:A and +8G:U substitutions included to enable RNA synthesis. For **B** and **D**, numbers above the violins indicate the number of bound lifetimes included in each distribution.

Since the CoSMoS assay cannot monitor branaplam binding directly, our single molecule data cannot by themselves reject an alternative model in which branaplam initially binds the U1 snRNP/-1A 5’SS duplex. In this alternate model, branaplam would then interact with the bulged nucleotide to relieve a steric clash and enable U1-C association, as has been proposed for risdiplam and branaplam-like molecules^10,16^. To distinguish between U1-C-first and branaplam-first models, we used MST to directly monitor the binding of branaplam to the U1 snRNP bound to a fluorescent 11bp-1A 5’SS RNA at the ensemble level (**Fig. 3C**). We pre-formed a U1 snRNP/11bp-1A 5’SS RNA complex using our U1 snRNP particle and observed the shift in fluorescence intensity upon binding branaplam. When performed in the absence of U1-C, we do not observe any appreciable binding to branaplam at up to 20 µM. Upon inclusion of U1-C in solution, we see a branaplam concentration-dependent response and can estimate a *K_D_* of 2.7 ± 0.4 x10^-6^ M for branaplam binding, which is consistent with EC_50_ values we measured using SPR and CoSMoS (**Fig. 2C, F**). This assay was repeated using the 11bp 5’SS RNA and, as expected, did not detect any binding to branaplam due to the absence of the -1A bulge (**Fig. S13A**). Finally, we replaced the U1 snRNP particle with a RNA mimicking the U1 snRNA 5’ end, annealed to the 11bp-1A 5’SS RNA, and assayed for branaplam binding by MST. Under these conditions, we also could not detect any binding of branaplam to the RNA:RNA duplex. (**Fig. S13B**). Combined, these data show that branaplam binds a -1A bulged U1 snRNA/5’SS duplex only in the presence of the U1 snRNP and U1-C and support the U1-C-first model.

We next determined if the U1-C requirement extended to other naturally occurring -1A bulged 5’SS, including those with clinical relevance. We examined four 5’SS associated with alternative splicing and known to be affected by branaplam or risdiplam (**Fig. 3D**)^14,15^. These include exon 7 of SMN2 which is involved in SMA, pseudoexon 50a of HTT which is involved in Huntington’s disease, pseudoexon 2a of SF3B3 which is used as a model 5’SS for targeted exonization by branaplam for gene therapy^32^, and exon 9 of FOXM1 which is an off-target of risdiplam and branaplam^14,33,34^. In the case of the HTT 5’SS, the endogenous sequence proved to be synthetically intractable due to a stretch of guanine bases. Therefore, we substituted guanines at the +7 and +8 positions with UA (HTT*).

CoSMoS was used to measure the extents to which U1-C and branaplam modulate each 5’SS interactions with U1 snRNP. For simplicity, each oligo was only examined at one concentration (SMN2: 10 nM, SF3B3: 10nM, FOXM1: 10 nM, HTT*: 3nM) across four conditions: +/- U1-C (0 or 100 nM) and +/- branaplam (DMSO or 10 µM) (**Fig. S14-17**). Consistent with the 9bp-1A 5’SS RNA, all four 5’SS display weaker binding to U1 and no effect upon addition of branaplam in the absence of U1-C. In the case of SF3B3, we were not even able to observe enough binding events above background to determine a bound lifetime in the absence of U1-C.

The addition of U1-C alone promoted binding of each 5’SS by at least a factor of 1.7-fold, and this was further enhanced by branaplam. The magnitude of branaplam-enhancement varies among the different 5’SS (SMN2: 3.7x, SF3B3: 7.7x, FOXM1: 5.5x, HTT* 3.6x), consistent with additional sequence dependencies beyond just the presence of a -1A bulged nucleotide^14,15^. Overall, our combined single molecule and ensemble data show that U1-C must be present for branaplam to bind to and modulate U1 snRNP/-1A 5’SS complexes and that the extent of binding enhancement is sequence-dependent.

### U1-C promotes and stabilizes 5’SS binding to U1 snRNP

Given the critical roles of U1-C for both duplex stability and branaplam enhancement, we next investigated the effect of U1-C on 5’SS recognition in general. For these CoSMoS measurements, we held the concentration of 9bp-1A 5’SS RNA constant at 1 nM and varied the solution concentration of the U1-C from 0 to 100 nM (**Fig. S18**). The average fraction bound increased as a function of U1-C concentration with an EC_50_ = 1.3 ± 0.2 nM (**Fig. 4A**).

**Figure 4.**
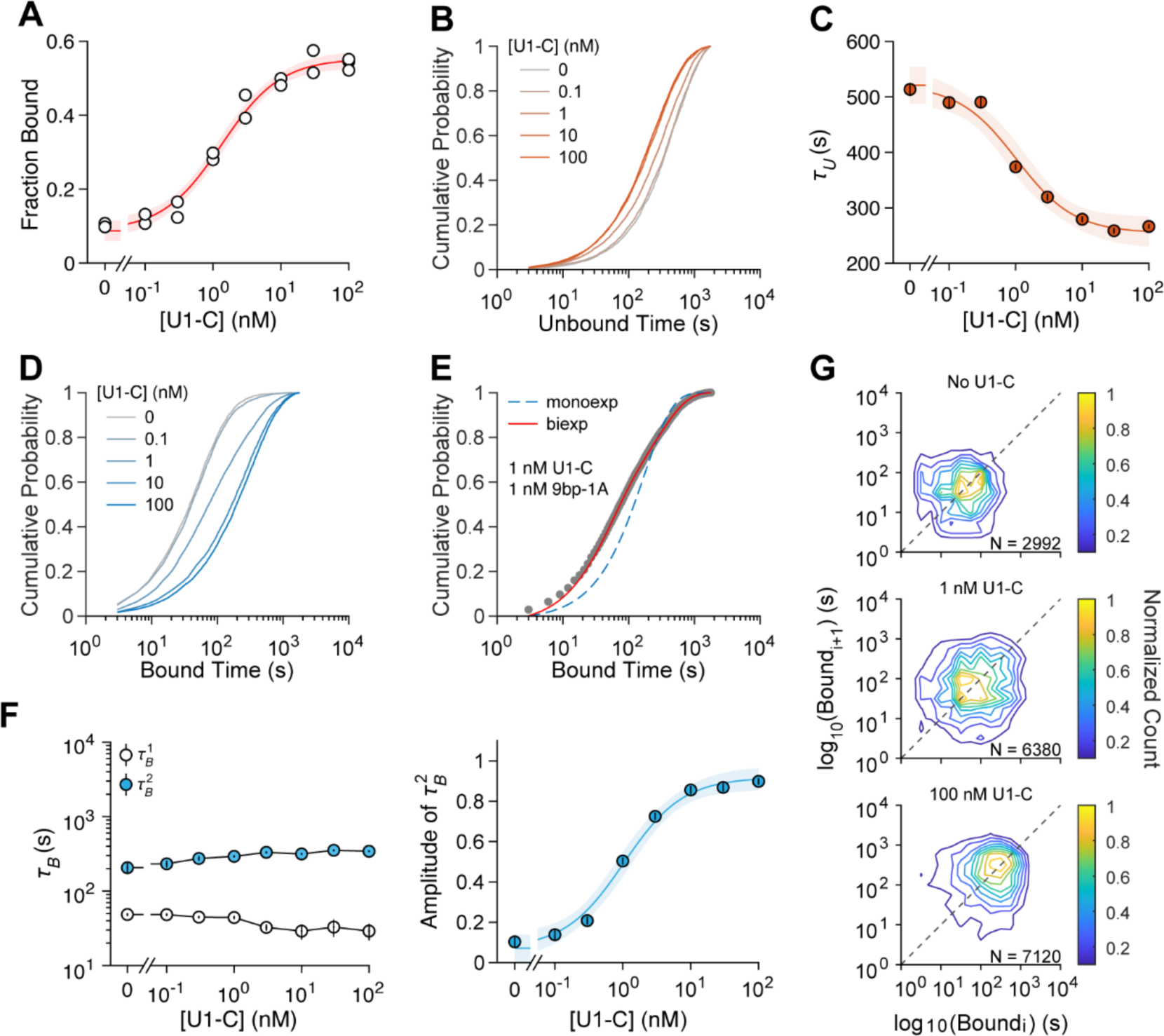
U1-C promotes and stabilizes the binding of a -1A RNA to U1 snRNP. (**A**) Dose response curves of 9bp-1A RNA binding to U1 snRNP vs. U1-C concentration. The red line and shading indicate the fit and 95% CI to the fitted equation to estimate an EC_50_ value (1.3 ± 0.20 nM). Cumulative probability distribution of unbound dwell times at 1 nM 9bp-1A RNA across U1-C concentrations (0 nM, *N* = 3106; 0.1 nM, *N* = 2920; 1 nM, *N* = 5543; 10 nM, *N* = 3169; 100 nM, *N* = 5154). (C) Unbound time constants determined from MLE of a monoexponential distribution for unbound dwell times overlaid with a fit to the fitted equation (EC_50_ = 1.8 ± 0.5 nM). (**D**) Cumulative probability distribution of bound dwell times at 1 nM 9bp-1A RNA across U1-C concentrations (0 nM, *N* = 2381; 0.1 nM, *N* = 2247; 1 nM, *N* = 4764; 10 nM, *N* = 3105; 100 nM, *N* = 5216). (**E**) Cumulative probability distribution of bound dwell times at 1 nM 9bp-1A RNA and 100 nM U1C (grey circles overlaid with MLE of mono (blue dashed) and biexponential distributions (solid red). (**F**) MLE of bound time constants (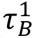 and 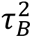, *left*) and amplitude of 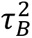 (*right*) of a biexponential distribution for bound dwell times at 1 nM 9bp-1A RNA across U1-C concentrations (value ± standard error). The amplitudes of 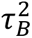 are overlaid with the equation fit (EC_50_ = 1.0 ± 0.2 nM). (**G**) Contour plots showing the correlation between successive bound event durations (*i* and *i* + 1) within individual molecules. Dashed line is the identity line.

We analyzed the single molecule dwell times of the unbound 9bp-1A 5’SS RNA events across these U1-C concentrations. We find that the RNA unbound lifetimes decrease with increasing U1-C concentration, demonstrating U1-C also helps promote the binding of a -1A 5’SS RNA (**Fig. 4B**). We estimated *k_apparent_* values using single exponential distributions and the resulting *k_apparent_* values yielded an EC_50_ = 1.8 ± 0.5 nM (**Fig. 4C**). Overall, we see the association rate at 1 nM 9bp-1A 5’SS RNA double between the absence and presence of saturating U1-C.

The larger effect of U1-C on the affinity for the 9bp-1A 5’SS RNA stems from an increase in the bound lifetime (**Fig. 4D**). MLE analysis of bound dwell times indicates that at least two time constants are required to describe the data (**Fig. 4E**). In contrast to the effect of branaplam on duplex lifetimes, we find that the value of the time constants does not significantly change across U1-C concentrations, but rather, their amplitudes change dramatically (**Fig. 4F**). On average, the 9bp-1A 5’SS RNA duplex exhibits 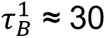 s and 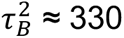 s^-1^. In the absence of U1-C, U1 snRNP can still form the longer-lived complexes with this RNA; however, these are rare relative to the shorter-lived interactions. Consistent with this, the amplitude of 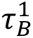 is dominant at low concentrations of U1-C, but 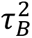 dominates at high concentrations. The change in the amplitude of 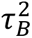 across U1-C concentrations yielded an EC_50_ = 1.0 ± 0.2 nM.

To test whether these two time constants reflect dynamic association/dissociation of U1-C with a kinetically homogenous population of U1 snRNP molecules, we correlated the dwell time durations of successive binding events within individual immobilized U1 snRNPs (**Fig. 4G**). At the extremes of either no U1-C or saturating U1-C, bound lifetimes of the 9bp-1A 5’SS RNA predominately align to a single cluster at the level of individual molecules corresponding to ether faster 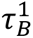 durations or slower 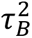. Near the EC_50_ of U1-C, we observe both short and long events corresponding to dynamic interconversion of the two time constants, which likely stems from the association and dissociation of U1-C. Higher concentrations of U1-C increase the probability of binding to U1 snRNP and the probability of observing a more stable duplex. Overall, these data show that U1-C dynamically binds U1 snRNP, and that its presence can help recruit and stabilize a -1A bulged 5’SS RNA.

### U1-C modulation of 5’SS binding is sequence dependent

We next aimed to determine if the U1-C dependent increase in *k_on_* and decrease in *k_off_* is generalizable to other 5’SS motifs using a small collection of designed 5’SS oligos (**Fig. 5A**). We first assayed if the association rate of a non-bulged 5’SS RNA (9bp 5’SS RNA) is enhanced by U1-C. We performed kinetic association experiments in the absence or presence of saturating U1-C at various 9bp (**Fig. S19**) or 9bp-1A 5’SS RNA (**Fig. S20**) concentrations. U1-C enhances the *k_on_* of 9bp by a factor of 2.1 and 9bp-1A by a factor of 1.9 (**Fig. 5B**), the latter of which confirms our result from varying U1-C at a constant RNA concentration (**Fig. 4C**). We could not conclude if U1-C decreases the *k_off_* of 9bp, as the estimated *k_off_* from non-equilibrium experiments in the absence of U1-C was not significantly different than the *k_off_* with saturating U1-C (*p* = 0.92, stepwise regression statistical test).

**Figure 5.**
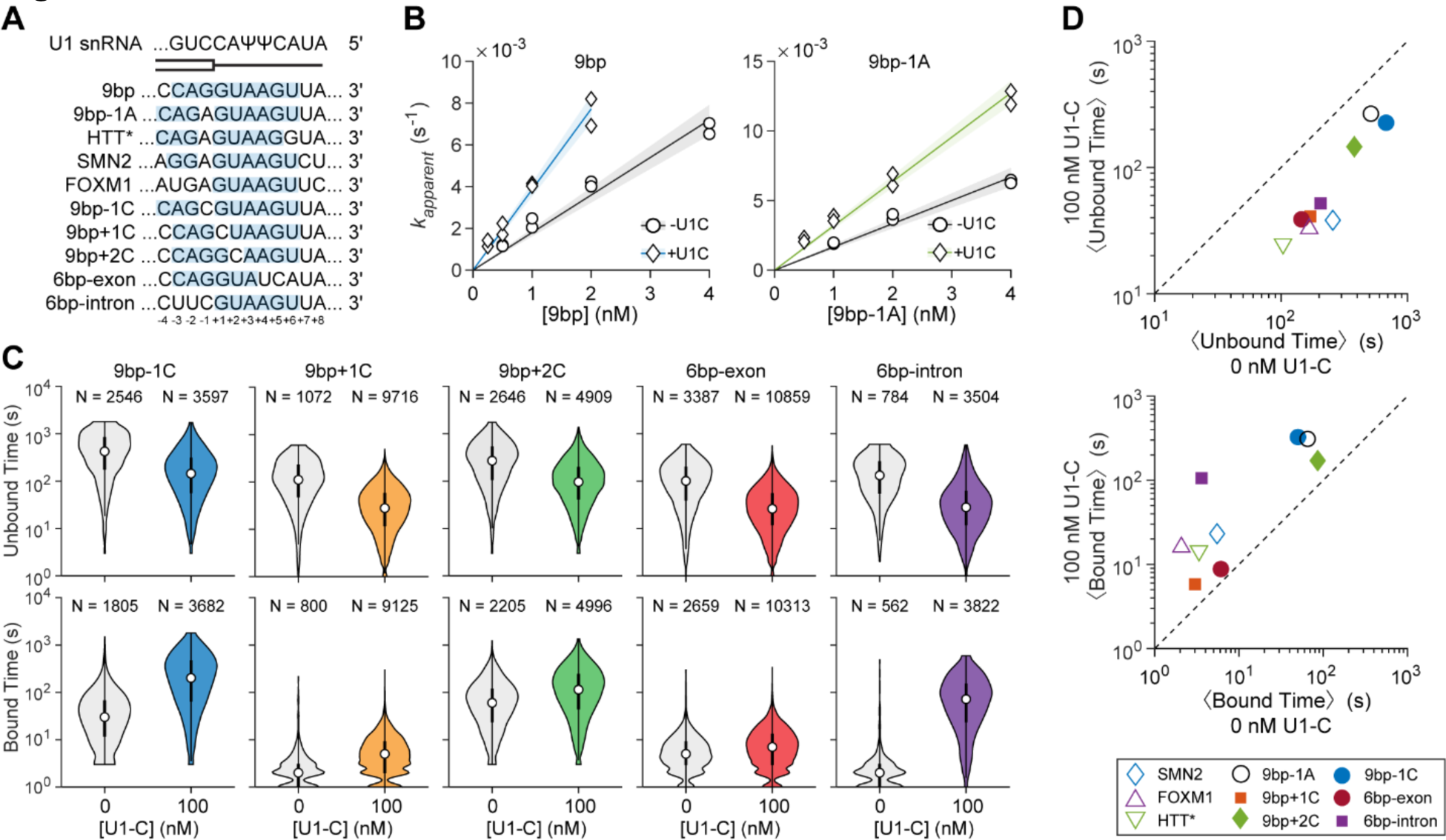
The Impact of U1-C on 5’SS Recognition is Sequence Dependent. (**A**) RNA oligos. Shaded region indicates predicted base pair interactions with the U1 snRNA (top). (**B**) Apparent association rates for 9bp (*left*) and 9bp-1A RNAs (*right*) at various concentrations in the absence and presence of saturating U1-C (100 nM). Data are overlaid with linear fits to determine *k_on_* (9bp with 100 nM U1-C: *k_on_* = 3.9 ± 0.4 x 10^6^ M^-1^ s^-1^, *R^2^* = 0.97; 9bp without U1-C: *k_on_* = 1.8 ± 0.5 x 10^6^ M^-1^ s^-1^, *R^2^* = 0.95; 9bp-1A with 100 nM U1-C: *k_on_* = 3.2 ± 0.6 x 10^6^ M^-1^ s^-1^, R^2^ = 0.98; 9bp-1A without U1-C: *k_on_* = 3.2 ± 0.6 x 10^6^ M^-1^ s^-1^, *R^2^* = 0.98). (**C**) Violon plots showing the distributions of unbound (*top*) and bound (*bottom*) dwell times for various RNAs at 0 and 100 nM U1-C. Data for 9bp-1A is presented at 1 nM. (**D**) Scatter plot showing the average unbound (*top*) and bound (*bottom*) dwell times of RNAs at 0 (x-axis) or 100 nM (y-axis) U1-C. Dashed line is the identify line.

In order to further investigate how U1-C enhances 5’SS recognition, we performed equilibrium binding measurements on various 5’SS RNAs to determine the effect of U1-C (**Fig. 5C, Fig. S21-25**). This collection of oligos included a -1C variant, which is known to form a bulged -1C but is not activated by branaplam^12^, a +1C mutant which can inhibit splicing^35,36^, and +2C, which converts the common +1/+2 GU 5’SS subtype to the less common +1/+2 GC motif (<1% human 5’SS)^17^. In addition, we designed two +1/+2 GU 5’SS-containing RNAs with 6 base pairs of complementarity to the U1 snRNA either on the exonic (-4 to +2 positions) or intronic (+1 to +6 positions) side of the exon/intron boundary. All experiments were conducted at a single concentration of the RNA (9bp-1C: 1 nM, 9bp+1C: 10 nM, 9bp+2C: 1 nM, 6bp-exon: 3 nM; 6bp-intron: 3nM; depending on their affinity) and either in the absence or presence of 100 nM U1-C.

For all of these RNAs, we see a decrease in the average unbound lifetimes, corresponding to a faster association rate, when U1-C is included relative to its absence (**Fig. 5D**, top). On average, U1-C increased the apparent association rate by 2.5-fold. This suggests that U1-C does not have a single, specific sequence requirement (*e.g.*, presence of a GU 5’SS) for facilitating RNA association to U1 snRNP.

In contrast, we see a larger variance in the effect on bound lifetimes by U1-C. The largest enhancement due to U1-C is a 29.3x increase in average bound lifetimes for the 6bp-intron 5’SS RNA (3.6s to 105.8s, **Fig. S25**) while the smallest is a 1.4x increase for 6bp-exon 5’SS RNA (6.1s to 8.8s, **Fig. S24**). The enhancement due to U1-C results in an ∼20-fold increased lifetime for the 6bp-intron U1 snRNP/5’SS RNA complex despite its predicted U1 snRNA/5’SS duplex being 2.2 kcal/mol less stable than that for the 6bp-exon 5’SS RNA (**Figure S3**). Therefore, U1-C stabilizes U1 snRNP interactions with RNAs dependent on the register of the 5’SS:U1 snRNA duplex and potential pairing potential with the U6 snRNA. In particular, this result reinforces observations made in yeast that Watson-Crick base-pairing potential alone is not predictive of 5’SS:U1 snRNP interaction lifetimes^37^.

The bound lifetimes of the 9bp-1C and 9bp-1A 5’SS RNAs are similar with (325.8s) and without U1-C (50.1s), despite no enhancement by branaplam in the case of the former (**Fig. S21**). Thus, the effect of branaplam is likely to be driven by the particular structural details of the -1A bulge rather than by inherently different kinetic properties of the U1 snRNP/5’SS RNA interaction. A +1C substitution that converts the canonical GU 5’SS to a CU nearly abolishes stable RNA binding (**Fig. S22**), and U1-C is ineffective at stabilizing the bound state. This suggests that U1 snRNP may enforce the requirement for a +1G at the 5’SS through kinetic selection against mismatches at this position. This selection results from both U1-C independent (poor binding in the absence of U1-C) and dependent (failure of U1-C to stabilize the bound state) components.

Finally, the 9bp+2C 5’SS RNA (which converts the +1/+2 GU 5’SS to the less frequent +1/+2 GC subtype) exhibits less stable binding than the +1/+2 GU 5’SS in the absence (86s) and presence (171s) of U1-C (**Fig. S23**). We estimate the 9bp 5’SS RNA with a “GU” motif binds nearly 10x more tightly compared to the GC variant. This correlates with results from high throughput *in vivo* splicing assays which showed strong preference for GU over GC^38^, reinforcing that splicing outcomes in cells are often dependent on U1-binding kinetics.

### A sequential binding model describes the mechanism of 5’SS modulation by branaplam

To fully describe -1A bulged 5’SS recognition and how it is influenced by U1-C and modulated by branaplam, we used hidden Markov modeling (HMM, see **Methods**) and simulations to fully define and evaluate potential kinetic mechanisms (**Fig. S26-S30, Table S3**). A sequential binding model provided the best description of our data (**Fig. 6**). This model predicts that 9bp-1A 5’SS RNA binding and unbinding can proceed through two pathways dependent on the presence of U1-C. The faster dissociation and slower association arise from U1 snRNP lacking U1-C, whereas the slower dissociation and faster association arise from U1-C bound to the U1 snRNP. Our model therefore predicts that the multiple components in unbound and bound dwell time distributions stem from the binding and unbinding of U1-C and are not inherent to the U1 snRNA:9bp-1A 5’SS interaction. Surprisingly, we find no mathematical evidence for the binding of U1-C once the 5’SS is already bound. We suspect that 5’SS:U1 snRNA duplex may sterically occlude the binding of U1-C, given that the U1-C zinc finger domain rests between the duplex and the SmD3 protein^21^.

**Figure 6.**
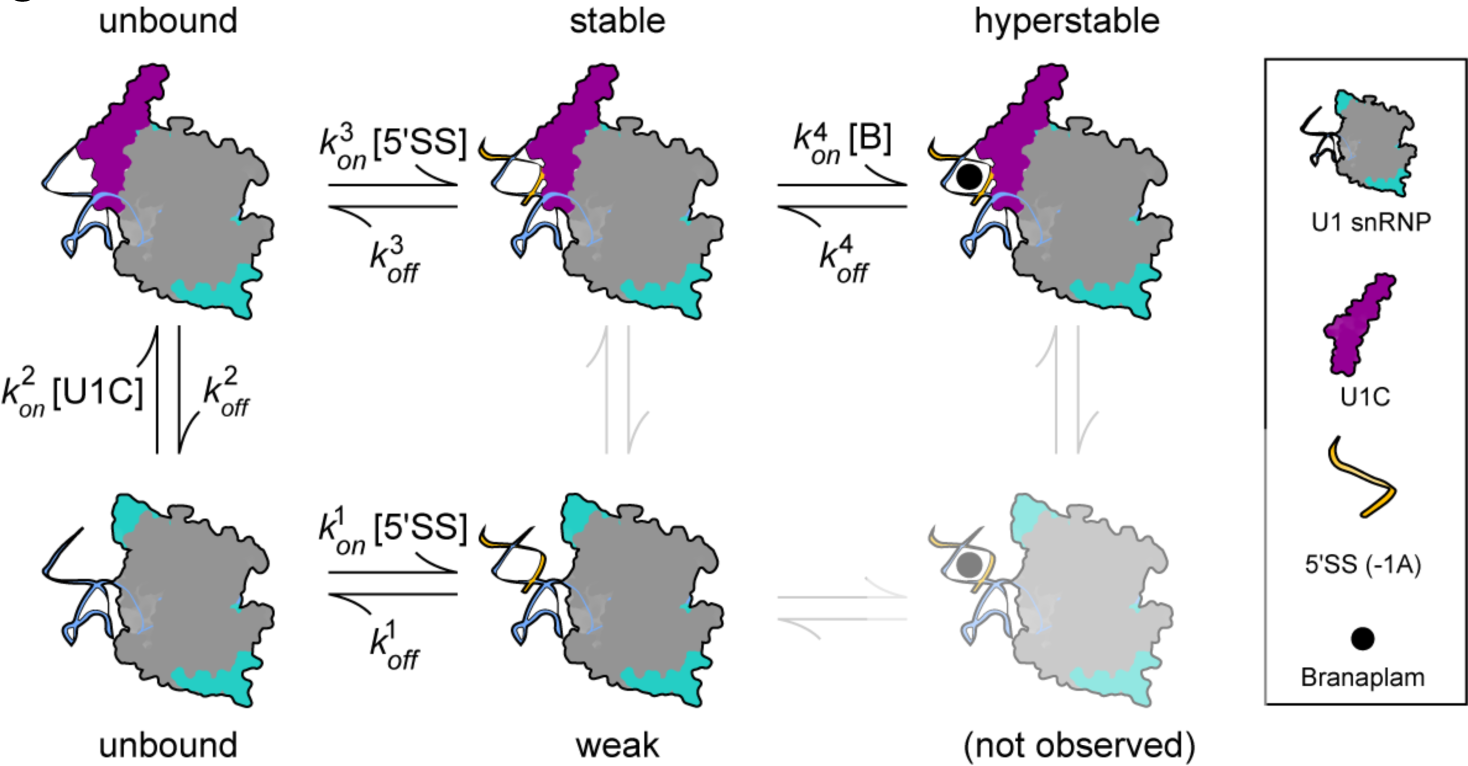
A sequential binding mechanism for -1A bulged 5’SS RNA recognition by U1 snRNP and modulation by branaplam. A kinetic model for -1A bulged 5’SS association and dissociation in the presence and absence of U1-C and branaplam. Optimized rate transitions for 9bp-1A are provided in **Table 1**. Equilibrium arrows in grey indicate transitions that are not supported by our experimental data or mathematical modeling.

**Table 1.** Globally optimized transition rates for 9bp-1A.

| Forward rate | Value ± SE (M<sup>-1</sup> s<sup>-1</sup>) | Reverse rate | Value ± SE (s<sup>-1</sup>) | K<sub>D</sub> ± SE (M) |
| --- | --- | --- | --- | --- |
| $k_{on}^1$ | 1.56 ± 0.01 × 10<sup>6</sup> | $k_{off}^1$ | 1.53 ± 0.01 × 10<sup>-2</sup> | 9.81 ± 0.09 × 10<sup>-9</sup> |
| $k_{on}^2$ | 2.43 ± 0.30 × 10<sup>6</sup> | $k_{off}^2$ | 7.69 ± 0.91 × 10<sup>-3</sup> | 3.16 ± 0.54 × 10<sup>-9</sup> |
| $k_{on}^3$ | 4.29 ± 0.06 × 10<sup>6</sup> | $k_{off}^3$ | 2.59 ± 0.03 × 10<sup>-3</sup> | 6.04 ± 0.11 × 10<sup>-10</sup> |
| $k_{on}^4$ | 1.27 ± 0.03 × 10<sup>4</sup> | $k_{off}^4$ | 8.40 ± 0.24 × 10<sup>-3</sup> | 6.61 ± 0.25 × 10<sup>-7</sup> |

Branaplam binding only occurs in the presence of U1-C and an existing -1A bulge U1 snRNA/5’SS duplex. Our model predicts a modest affinity for the branaplam interaction (*K_D_* = 6.59 ± x 10^-7^ M), which is similar to the *K_D_* determined via MST (**Fig. 3C**). Surprisingly, the dissociation rate of branaplam is three times faster than the dissociation rate of the 5’SS RNA when U1-C is bound. The increase in bound lifetimes with increasing branaplam concentrations is supported by this model. High concentrations of branaplam lead to its faster association relative to the dissociation rate of the 5’SS RNA, thereby driving the equilibrium toward branaplam rebinding. The dynamic binding of branaplam contributes to the observed high stability of the U1snRNP/-1A 5’SS RNA complex. We were able to confirm this prediction using SPR and by washing branaplam from solution after an initial binding period to the U1 snRNP/11bp-1A complex. Under these conditions, we do not see increased stability of the 11bp-1A RNA consistent with branaplam reversibly binding the complex and dissociating during the wash (**Fig. S31**). Our work therefore shows that transient, reversible drug binding can nonetheless contribute to formation of very long-lived U1 snRNP/5’SS interactions.

## DISCUSSION

Herein, we used a reconstituted U1 snRNP to study the detailed kinetics of its interactions with 5’SS RNA oligos and how these interactions change upon the inclusion of a small molecule splicing modulator. Both single molecule and bulk biophysical measurements show that U1 snRNP binds RNA in a sequence-specific manner, branaplam extends U1 snRNP/oligo lifetimes of -1A bulged 5’SS if U1-C is present, and that splicing modulation can involve a complex, multi-step process. The origin of this complexity is in part due to reversible binding of the U1-C component, which dynamically interacts with the snRNP. U1-C itself both promotes RNA binding by U1 snRNP and stabilizes the U1/RNA complex in a sequence-specific manner. We were able to use our feature-rich and large single molecule data sets to determine a sequential binding mechanism for U1 snRNP, U1-C, and branaplam interactions with a -1A 5’SS-containing oligo. In this mechanism, U1-C associates with the snRNP prior to 5’SS binding and decreases the *K_D_* for the 5’SS sixteen-fold. Branaplam then reversibly associates with this complex with a moderate *K_D_* (∼0.7 µM).

### Differences and Similarities Between Yeast and Human U1 snRNPs

Our work here with human U1 snRNP recapitulates several features of 5’SS recognition that we previously characterized in the much larger yeast complex^37,39^. Both human and yeast U1 snRNPs exhibit short and long-lived interactions with RNAs. In yeast, the data were consistent with a two-step binding mechanism in which RNAs could either be released from the snRNP or the U1 snRNP/RNA complex would transition to a more tightly bound state. This transition is possibly due to a conformational change in U1 snRNP that could involve the U1-C and Luc7 proteins^22,37,40,41^. In human U1 snRNP interactions with a -1A 5’SS, there is complexity due to dynamic binding of U1-C; however, we do not detect transitions or conformational changes of the U1 snRNP/U1-C/5’SS complexes between kinetically-distinct states. Thus, the origins of the complexity are different in each system, and this could reflect fundamental differences between the yeast and human splicing machinery. In yeast, there are fewer and smaller introns^42^, and U1 snRNP is ∼10-fold less abundant than in humans (∼0.3 µM in yeast vs 2.5 µM in human nuclei)^43^. Moreover, interacting domains between U1 snRNP and RNAP II found in humans are not conserved^44^ and putative yeast-specific Pol II interaction domains on U1 have no effect on co-transcriptional U1 snRNP recruitment or the early stages of spliceosome assembly^45^. Based on these observations, we hypothesize that rapid surveillance of the transcriptome for 5’SS enabled by reversible, two-step binding by yeast U1 snRNP may enable efficient splicing in yeast but be less essential in humans wherein U1 snRNP can be localized proximal to potential 5’SS as they are synthesized by RNAP II^44^. This hypothesis is also consistent with the lifetimes of yeast and human U1 interactions with highly complementary RNAs. The lifetimes of yeast U1/5’SS complexes are much shorter than their human counterparts (a longest-lived lifetime of <400 s vs. ∼1563 s for RNA oligos with 9 bp of complementarity for yeast and human U1 snRNPs, respectively)^37^. Long-lived U1 snRNP/5’SS interactions may be important for maintaining 5’SS definition and RNAP II elongation rates while long introns and the downstream exon are transcribed in humans^46^.

In both yeast and humans, base-pairing potential and predicted thermodynamic stability of the snRNA/5’SS duplex are not good predictors by themselves of the U1 snRNP/5’SS interaction lifetime. Neither yeast nor human U1 snRNP efficiently stabilizes duplexes containing mismatches at the +1 position even when extensive base pairing interactions are predicted at other positions. A +1G at the 5’SS is critical for formation of an atypical base-pairing interaction with the -1G of the 3’SS during exon ligation, which occurs in a spliceosome complex that lacks U1 snRNP^20^. Moreover, human U1 snRNP binds more stably to 5’SS with complementarity in the +1 to +6 (intronic) positions over the -4 to +2 (exonic) positions despite the greater predicted thermodynamic stability of the latter. This may reflect the importance of specifying the intronic portion of the 5’SS sequence in order to permit proper pairing with the U6 snRNA during the catalytic steps of splicing^20^. A similar, “ruler-like” function was previously noted for the yeast U1 snRNP^37^. Thus, kinetic selection of substrates chemically competent for splicing is a conserved feature of both human and yeast U1 snRNPs even though neither U1 snRNP is present during the transesterification steps.

### The Role of U1-C in 5’SS Recognition

For human U1 snRNP, selection against non-+1G 5’SS appears to involve both failure of the snRNP to associate stably with these RNAs in the absence of U1-C and failure of U1-C to stimulate formation of a longer-lived state. What causes the weaker binding of the RNA with a mismatch at +1 in the absence of U1-C is not clear but could be a combination of reduced overall stability of the remaining base-pairing interactions and the mismatch interfering with contacts between the snRNA/5’SS duplex and the nearby U1 snRNA helix H^21^. Failure of U1-C to stabilize binding of the RNA lacking a +1G is consistent with the protein’s location adjacent to the +1G base-pairing site with the snRNA^21^. Another prediction of our kinetic model is that U1-C can only associate with U1 snRNPs in the absence of 5’SS pairing. In other words, U1-C must be pre-associated with U1 snRNP prior to its engagement with RNA in order to modulate 5’SS recognition.

U1-C interacts dynamically with the snRNP in the absence of 5’SS pairing. We do not believe that this was due to our use of a model, reconstituted U1 snRNP since U1-C has previously been observed to be a salt labile component of endogenous U1 snRNPs^23^. In addition, interpretable density due to U1-C is absent in a cryo-EM structure of endogenous human U1 snRNP bound to RNA polymerase II despite its presence during the purification^44^. In our experiments, U1-C has an EC_50_ of ∼1 nM, and our model predicts a *K_D_* of ∼0.3 nM. While U1-C binds tightly, the calculated lifetime of the U1/U1-C complex is only ∼2 min in the absence of a paired RNA, less than the amount of time RNAP II would take to transcribe the average human gene (∼10 min for a median human gene size of 24 kbp^47^). Based on these kinetics, it should not be assumed that a U1-C-containing U1 snRNP that is recruited to the transcriptional machinery for co-transcriptional spliceosome assembly still contains U1-C at the moment a 5’SS is transcribed.

Combined, the above observations suggest that U1-C-lacking and U1-C-containing U1 snRNPs may both be biologically relevant for gene expression. In terms of splicing, it is intriguing to speculate that U1-C-lacking snRNPs could be fully functional in the splicing reaction for both 5’SS recognition and transfer to the U6 snRNA. If true, this could have implications for splicing regulation, the ATP requirement for 5’SS transfer to U6, and how 5’SS compete for U1 snRNPs when U1-C is limiting (*e.g.,* a hungry spliceosome-like mechanism^48^). Based on our data, we would predict that strong, highly complementary splice sites would be the most likely to stably recruit U1/ΔU1-C snRNP particles and that these sites could be preferentially used by the spliceosome when U1-C is limiting. In *Drosophila*, knockdown of U1-C appears to have this very effect and reduces usage of the nonconsensus *dAdar* exon 3a 5’SS in favor of the stronger exon 3 5’SS^49^. Alternatively, U1 snRNPs containing or lacking U1-C could aid in distinguishing complexes involved in spliceosome assembly from those functioning in other processes such as telescripting^50^.

As noted above, whether or not U1/5’SS interactions are influenced by the presence or absence of U1-C depends on the sequence of the 5’SS: a highly complementary RNA oligo binds tightly even in the absence of U1-C while U1-C can greatly enhance the bound lifetimes of RNAs containing mismatches. These properties—reversible binding to U1 and RNA sequence-specific effects—show that U1-C has features in common with many alternative splicing factors. In humans, U1-C may have more in common with these factors than with other constitutive components of the splicing machinery.

### Mechanism of 5’SS Modulation by Branaplam

While it may seem counterintuitive that a reversibly-binding splicing modulator leads to formation of long-lived interaction between U1 snRNP and -1A bulged 5’SS, our kinetic mechanism provides a rationale for this observation. The -1A 5’SS RNA can only dissociate from U1 snRNP when branaplam is not bound and rapid re-binding of branaplam limits the lifetime of this state.

Recently, a thermodynamic model for splicing modulator drug action has been proposed based on RNA-Seq data and measurements of mRNA production in cells^14^. In this work, the authors proposed two branaplam-binding modes for U1 snRNP: a risdiplam-like binding mode that occurs on -1A bulged 5’SS that also contain a -2G and a second state that leads to hyperactivation of some 5’SS that additionally contain a -3A. It is unlikely that these two states are due to presence/absence of U1-C since our data shows that branaplam can only bind U1 snRNP when U1-C is present.

While we did not study the sequence requirements for hyperactivation explicitly, we do note that the 5’SS RNA oligos that showed the largest changes in U1 snRNP bound state lifetimes were also those with the hyperactivation “AGA” motif (9bp -1A, SF3B3, and HTT*). These results suggest that the hyperactivation phenotype has a kinetic basis and might be due to larger changes in the lifetime of the U1 snRNP/5’SS interaction. In addition, while the thermodynamic model included two different branaplam-binding modes, the authors were not able to determine if the risdiplam-like binding mode is a necessary precursor for formation of the hyperactivated state. Our single molecule data supports only a single branaplam-bound state for the U1 snRNP/U1-C/5’SS complex. The risdiplam-like and hyperactivated binding modes of branaplam likely occur independently of one another, each involving particular molecular interactions with their corresponding 5’SS.

Finally, our results suggest that drug development efforts for new modulators that target other -1A bulged 5’SS RNA targets or different mismatched/weak 5’SS should consider the U1 ribonucleoprotein target rather than optimizing RNA duplex interactions alone. Combined with our observation of the reversibility of U1-C association, it is also likely that branaplam’s ability to act as a splicing modulator of a given transcript is a function of U1-C binding and re-binding rates. Differences in binding/re-binding rates for both branaplam and U1-C could contribute to variation in splicing modulation that has been observed among different transcripts^12,14^. Splicing modulation may, therefore, be limited in part by the inherent kinetic properties of the factors and processes involved.

## MATERIALS AND METHODS

### RNA oligonucleotides

RNA oligonucleotides (**Table S1**) for SPR, MST, and single-molecule experiments were purchased from Integrated DNA Technologies (IDT, CoSMoS), Dharmacon (SPR), or Metabion (MST). Stocks of fluorescent RNAs intended for single-molecule experiments were prepared by resuspending the lyophilized oligonucleotides in nuclease-free water (20-50 µM, Ambion). RNA concentrations were calculated from their absorbance values 260nm using a NanoDrop and the extinction coefficients from IDT via the Beer-Lambert law.

### Protein Purification

Individual components and reconstitution of the minimal U1 particle were produced and purified as previously described^21^. All plasmids were purchased from GenScript (Piscataway, NJ, USA) based on the pET-28a(+) backbone and codon optimized then transformed into *Escherichia coli* BL21 Star (DE3) cells (Cat# C601003, ThermoFisher Scientific, Waltham, MA, USA).

For the U1-70K_SmD1/D2 polycistronic construct, an N-terminal thioredoxin-6xHis-tag followed by a tobacco etch virus (TEV) protease cleavage site was appended to the U1-70K fragment comprised of residues 2-59 followed by a Gly-Ser triplet linker then residues 7-91 of SmD1. A second open reading frame containing SmD2 was comprised of residues 1-118. For the SmD3/B polycistronic construct, an N-terminal 6xHis-tag followed by a TEV cleavage site was appended to residues 1-126 of SmD3, which was followed by a second open reading frame for residues 1-95 of SmB. For the SmF/E/G polycistronic construct, an N-terminal His-SUMO-Avi tag was introduced prior to residues 1-75 of SmF, followed by additional open reading frames for SmE (residues 1-92) and SmG (1-76). For SPR and MST experiments, a construct with only a His-SUMO tag was used. For U1-C_61, a C-terminal 6xHis-tag was added after residues 1-61 of U1-C.

Cells were cultured at 37°C in 1L of 2xYT media supplemented with kanamycin (50 µg/mL) then induced with 0.5mM IPTG at 16°C overnight. Cell pellets were resuspended in lysis buffer (20mM HEPES, 1M KCl, 1M Urea, 5mM TCEP, pH 7.5) plus cOmplete ULTRA EDTA-free protease inhibitor cocktail (Roche, Basel, Switzerland), then lysed by sonication. Clarified lysates were diluted in IMAC Buffer A (20mM HEPES, 1M KCl, 1M Urea, 2mM TCEP, pH 7.5) then loaded onto a HisTrap HP 5mL Ni-NTA column (Cytiva, Marlborough, MA, USA) and eluted with a gradient of IMAC Buffer B (20mM HEPES, 1M KCl, 1M Urea, 2mM TCEP, 300mM Imidazole, pH 7.5). Eluted fractions were pooled in dialysis tubing (Cat# 68035, ThermoFisher Scientific) with TEV protease and dialyzed overnight against 20mM HEPES, 250mM KCl, 1M Urea, 2mM TCEP, pH 7.5. Following dialysis, the solution was adjusted to 1M KCl and loaded onto a HisTrap column equilibrated in IMAC Buffer A. The flow-through was collected and injected onto a Superdex HiLoad 75 26/60 column (Cytiva) equilibrated in 20mM HEPES, 250mM KCl, 2mM TCEP, pH 7.5 and the fractions were collected then concentrated by centrifugation (Cat# UFC9003, MilliPore Sigma, Burlington, MA, USA).

For the SmF/E/G trimer, cells were cultured like the other constructs with the addition of 1% (w/v) glucose to the culture media. Protein was similarly purified via the 6xHis-tag, cleaved by SUMO protease (Cat#12588018, ThermoFisher Scientific), and finally purified by IMAC and SEC as described. For single molecule studies, the SmF/E/G trimer was biotinylated on the SmF AviTag using the BirA biotin-protein ligase reaction kit (Avidity LLC, Aurora, CO, USA) and biotinylation was confirmed by MALDI-TOF MS.

### U1 snRNA Production

The U1snRNA used for reconstitution of the minimal U1 particle was purchased from AxoLabs (LGC Group, Kulmbach, Germany) and dissolved to a concentration of 500µM in RNAse-free ddH_2_O comprising the sequence: 5’-AmUmACψψACCU GGCAGUGACC ACCACACACU GCAUAAUUUG UGGUAGUGGG CGAAAGCCCG-3’, where Am and Um represent 2’-*O*-methyl nucleotides, and ψ is pseudouridine. To enable single molecule studies, a U1 snRNA of the same sequence was produced with an aminohexyl linker on the 3’ end that was subsequently labeled with Cy5 NHS ester as the fluorophore.

### Minimal U1 snRNP Reconstitution

Prior to complex reconstitution, U1snRNA was prepared by refolding at 80°C for 3 min and then cooling on ice for 10 min. In a pre-warmed solution of Reconstitution Buffer (20mM HEPES, 250mM KCl, 2mM TCEP, pH 7.5) containing 40 U/mL RNAsin (Cat#N2111, Promega, Madison, WI, USA), each Sm protein sub-complex was combined to a final concentration of 8 µM and incubated for 5 min at 37°C. To this, U1 snRNA was added to a final concentration of 4 µM and incubated for 45 min at 37°C. When required, U1-C_61 can be added to a final concentration of 8 µM, incubated for 15 min at 37°C, then the complex is cooled overnight at 4°C. The crude complex was then loaded onto a MonoQ 10/100 GL column (Cytiva) in Reconstitution Buffer and eluted with a gradient of Reconstitution Buffer containing 1M KCl. Eluted fractions were pooled and loaded onto a Superdex column (Cytiva) equilibrated in Reconstitution Buffer. After purification, fractions corresponding to minimal U1 were concentrated using a 30kDa MWCO centrifugal filter (Cat#UFC9030, MilliPore Sigma).

### Surface plasmon resonance (SPR)

For SPR assays, a Biacore 8K (Cytiva) was used with a streptavidin-coated Series S Sensor Chip SA (Catalog #BR100531). The instrument was equilibrated in 20mM HEPES, 200mM KCl, 5mM MgCl2, 0.5mM TCEP, 0.05% (v/v) Tween-20, and 2% (v/v) DMSO. For immobilization, RNA was synthesized with a 3’-biotin (-1A bulge: CAGAGUAAGUAU; SMN2: AGGAGUAAGUCU; Match: CAGGUAAGUAU; Reverse: UAUGAAUGGAC; Dharmacon) and injected at 1nM with 30s contact time at 10 µL/min to afford 3-5RU of capture. U1 snRNP binding studies were performed by injecting a titration of complex that was serially diluted from 100nM with a contact time of 180s and a dissociation time of 600s at a flowrate of 30 µL/min in duplicate. After each cycle, the chip surface was regenerated by an injection of 3M MgCl_2_, with a 30s contact time at 30µL/min, followed by a 50% (v/v) DMSO needle wash. All data was analyzed after reference subtraction, blank subtraction, and a 1.5-2.5% (v/v) DMSO solvent correction applied. Data were analyzed using a one-state Langmuir binding model in the Biacore Insight Software. To assess the effect of ligand on U1 snRNP binding kinetics, a “co-inject” format was used to allow for compound to be present during the association and dissociation phases of the experiment. A protein solution of U1 snRNP was prepared at 10nM with varying concentrations of Branaplam serially diluted from 5µM and injected across the immobilized RNA surface as previously described.

### MicroScale Thermophoresis (MST)

For branaplam binding assays, a serial dilution was performed in DMSO at 25x final concentration, followed by dilution with assay buffer (20mM HEPE, 200mM NaCl, 1mM MgCl2, 1mM TCEP, 0.05% Tween-20, pH 7.0) to 2x final concentration. Separately, a pre-formed complex of U1 snRNP ΔU1-C was prepared at 200nM in the presence of 20nM of a 5’SS oligonucleotide labeled with Cy5 (5’-Cy5-CAGAGUAAGUAU; Metabion) with or without the addition of 500nM U1-C supplementation. The branaplam titration was then mixed 1:1 with the pre-formed U1 snRNP complex and incubated at room temperature for 15 min before loading Monolith LabelFree Premium Capillaries (MO-Z025; NanoTemper Tech). Capillaries were analyzed by a Monolith X red-continuous instrument (NanoTemper Tech) at 25°C with 100% LED and laser power.

For U1-C binding assays, U1-C protein was serially diluted at 2x final concentration in assay buffer. Separately, a pre-formed complex of U1 snRNP ΔU1-C was prepared at 200nM in the presence of 20nM of a 5’SS oligonucleotide labeled with Cy5 with the addition of branaplam at 5µM or DMSO, to yield a final DMSO concentration of 2% (v/v). The U1-C titration was mixed 1:1 with the pre-formed U1 snRNP complex and assessed on a Monolith X as previously described.

### Equation fitting

Dissociation constants (*K_D_*) and EC_50_ values from CoSMoS (e.g., **Figure 2A-B**), SPR (e.g., **Figure 1D**) and MST (**Figure 2J**) were obtained by fitting binding data to Hill Equation 1

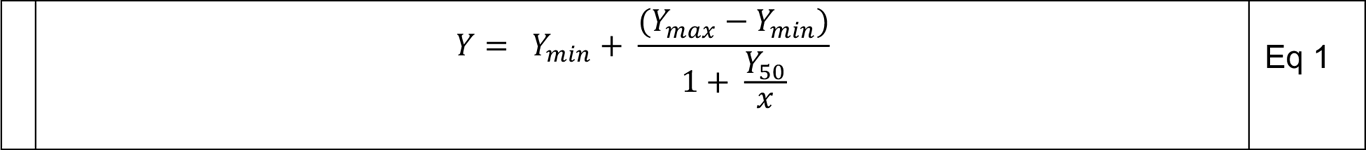

where *Y*_50_, is EC50 or *K_D_*, depending on the data, Y is the observed response (i.e., fluorescence) and *x* is the concentration. Curves were fit using nonlinear regression using the custom code in MATLAB with the function *nlinfit*. 95% confidence intervals were computed from the estimated Jacobian returned by nonlinear least squares fitting via *nlparci*.

### Single molecule chamber preparation

Single molecule imaging chambers were prepared using microscope slides (24mm x 60mm, #1.5, GoldSeal) and cover glasses (25mm x 25mm, #1.5, Corning) at least one day before each experiment. Substrates were first cleaned by successive sonication in 2% v/v Micro-90, 100% ethanol, and 1M KOH for 60 min each in slide-mailers (Fisher Scientific), with a MillliQ water wash after each step. Cleaned substrates were then dried with high purity nitrogen (Airgas) and aminosilanized with 1.5% (v/v) VECTABOND (Vector Laboratories) in acetone (Spectrophotometric Grade, Alfa Aesar) for 10 min. Substrates were washed with 100% ethanol, dried with nitrogen, and passivated by incubation of a 1:100 w/w mixture of mPEG-biotin-SVA (Laysan Bio) and mPEG-SVA (Laysan Bio) in 100mM NaHCO_3_ (pH 8) overnight. Prior to use, the substrates were rinsed with MilliQ water and dried with nitrogen. Imaging chambers were created by placing thin strips of double-sided tape and vacuum grease along the glass slide and adhering a cover glass on top. This typically resulted in three 25µL volume lanes per slide. Prior to use, assembled chambers were rinsed with at least 200 µL of wash buffer (WB: 20mM HEPES pH 7.5, 200mM KCl, 1mM MgCl_2_, 0.5% v/v Tween20, 0.1% w/v PEG8000).

### Single molecule data acquisition

Single molecule data were collected on a custom-built micro-mirror total internal fluorescence microscope system as previously described^37,51^. Each experiment utilized a 532 nm (CrystaLaser) and 633nm (Power Technology Inc.) laser for fluorophore excitation, and a 785nm laser (Power Technology Inc.) for continuous focal plane drift correction (TIRF-Lock, Mad City Labs). Laser powers were measured before each video (532nm, 900-1400µW; 633nm, 800-1400 µW; 785 nm, 250µW) at a point prior to the objective in the optical path. Excitation and emission passed through a 60x 1.49 NA oil immersion objective (Olympus). Emission was passed through a short-pass filter (FES0700, ThorLabs) and imaged onto two separate sCMOS detectors (Hamamatsu ORCA-Flash4.0 V3). All imaging was controlled with Micro-Manager 2.0^52^ and the TIRF-Lock was controlled with LabView. All images were collected 2x2 pixel binning with active hot pixel correction.

Immediately prior to data collection, streptavidin-labeled fluorescent beads (T10711, Invitrogen) were flowed into the lane at a low concentration (∼5x10^4^-fold dilution from stock in WB) to serve as fiducial markers for channel alignment and lateral drift correction. The lane was then washed with 50µL of 0.2 mg/mL streptavidin (SA10-10, Agilent) for two minutes, followed by 50 µL of WB to remove unbound beads and streptavidin. The U1 snRNP particle labeled with Cyb5 was then diluted to 10-20 pM in WB and incubated in a lane for one minute, followed by a 50µL wash with WB. The surface density of U1snRNP-ΔU1-C-Cy5 was checked by flowing in imaging buffer (IB: 20mM HEPES pH 7.5, 200mM KCl, 1mM MgCl_2_, 0.05% v/v Tween20, 0.1% w/v PEG8000, 5mM protocatechuic acid (PCA), 1U/mL protocatechuic-dioxygenase (PCD), 1mM Trolox, 2% (v/v) DMSO.

Two imaging schemes were used for data collection: alternating laser excitation or sequential laser excitation. In alternating laser excitation, a 50 µL solution containing variable concentrations of RNA, U1-C, and branaplam in IB was added and successive images were captured with a 1s exposure under 532 nm then 633 nm excitation, separated by a 400 ms switching time. In sequential laser excitation, approximately 30s were first recorded (633nm, 1 Hz) to identity areas of interest (AOI) followed by addition of 50 µL solution containing variable concentrations of RNA, U1-C, and Branaplam in IB was added and images were captured sequentially at 1 Hz (532nm, 1s exposure). Regardless of imaging scheme, a total of 600 frames were collected across varying frame rates (0.11 to 1 Hz) to minimize photobleaching. Finally in both schemes, the lane was washed with WB to remove oxygen scavengers and images were collected under 633nm excitation until all surface tethered molecules photobleached (typically 30-60 frames). Importantly, this step allowed us to ensure we only analyzed AOIs featuring a single U1 snRNP molecule.

Data were collected under both non-equilibrium and equilibrium conditions. In the case of non-equilibrium experiments (*e.g.*, 9bp RNA), imaging commenced immediately after addition of RNA to the lane, with the goal of characterizing initial association rates. In the case of equilibrium binding experiments (*e.g.*, 9bp-1A RNA), the RNA solution was incubated on the lane for up to 15 min time to reach equilibrium (the rate of which was determined in pilot experiments not reported herein) prior to the start of imaging. No lane was used for longer than two hours after U1snRNP deposition to minimize possible surface effects. A summary of each dataset is found in Table S2.

### Single molecule data analysis

Raw tiff stacks from Micro-Manager 2.0 were processed using custom code written in MATLAB (MathWorks, see Software Availability below). The 532nm channel (RNA) was mathematically mapped to the 633nm channel (surface tethered U1) in a two-step process using the fluorescent beads present in each spectral channel. First, a nonreflective similarity transformation was applied to align the detectors in physical space (rotation and translation), followed by an affine transformation using the beads as anchor points to correct for chromatic aberrations between the spectrally separated emitters (rotation, translation, and shear). Lateral drift was automatically corrected in each image by computing a nonreflective similarity transformation between temporally separated images. Areas of interest (AOIs) likely containing at least one single molecule were detected in a 633nm channel by averaging the first five frames and performing a generalized likelihood ratio test on the resulting image^53^. Detected AOIs were fit to a two-dimensional gaussian function within a 5x5 pixel space. AOIs were filtered by removing those with intensity values of greater than three scaled median absolute deviations from the median (*e.g.*, beads, multiple overlapping molecules) and those with a Euclidean distance less than 5 pixels away from a neighboring AOI. Accepted AOIs were then mapped to the 532nm channel using the mathematical transformations described above. The time dependent fluorescence of each AOI in each channel was computed by integrating over all frames in a 3x3 pixel space centered on each AOI’s sub-pixel location. All the steps of this process were incorporated into a graphical user interface (*smVideoProcessing*).

Fluorescence trajectories of each AOI in both 633nm (surface U1 snRNP, Cy5) and 532nm channels (solution RNA binding, Cy3) were first idealized using the divisive segmentation and clustering algorithm (DISC)^54^. Only trajectories showing single step photobleaching or photoblinking of Cy5 (633nm channel) were included for further analysis. For binding data, time series idealization enabled a binary interpretation of the time-dependent signal, whereby each frame was classified as a 0 for unbound or 1 for bound. All identified binding events were visually inspected to ensure on-target binding by examining a cropped image at each AOI for each frame in both channels (*i.e.*, image gallery). Events determined to be off-target binding (*e.g.*, a fluorescent RNA diffused within the AOI) were manually removed in the idealized time series. In the case of sparse, fast binding dynamics a variational Bayesian hidden Markov modeling algorithm (vbFRET) was used instead of DISC^55^, owning to their known differences in event detection accuracy^54^. Only molecules exhibiting at least one binding event were included for further analysis. All the steps of this process were incorporated into a graphical user interface (*smTraceViewer*).

### Single molecule dwell time analysis

Unbinned dwell time distributions were treated as a mono- or bi-exponential distributions and the underlying parameters of each distribution were estimated using maximum likelihood (MLE)^29^. We used a conditional probability density function (PDF) that accounts for the experimental limitations of frame rate (*t_min_*) and experiment duration (*t_max_*), as any observed dwell time *t* must therefore be *t_min_* ≤ *t* ≤ *t_max_*. The conditional PDFs of mono (PDF1) and biexponential (PDF2) distributions are given by (Eq 2 and 3)

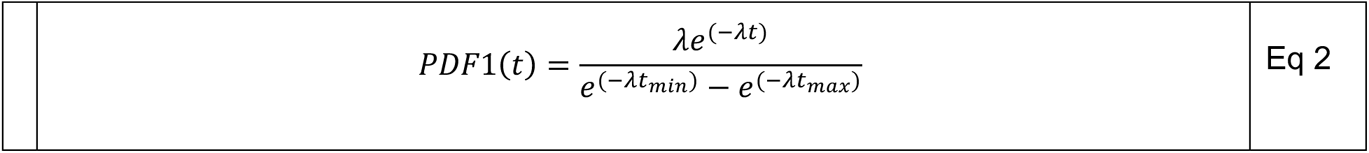

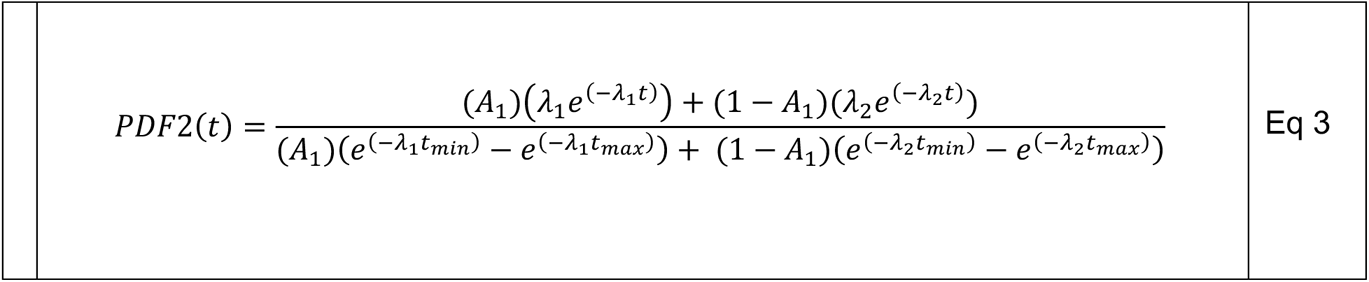

whereby each exponential distribution is described by a rate constant *λ_n_* and an amplitude *A_n_*where 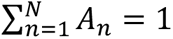. Rate constants obtained from MLE are often interpreted as their corresponding time constants (τ) rather than rates (*λ*) by the relationship *λ* = τ^-1^. A log likelihood-ratio (LLR) test was performed for each bound dwell time distribution to compare the goodness of fit of mono- and biexponential distributions (Eq. 4).

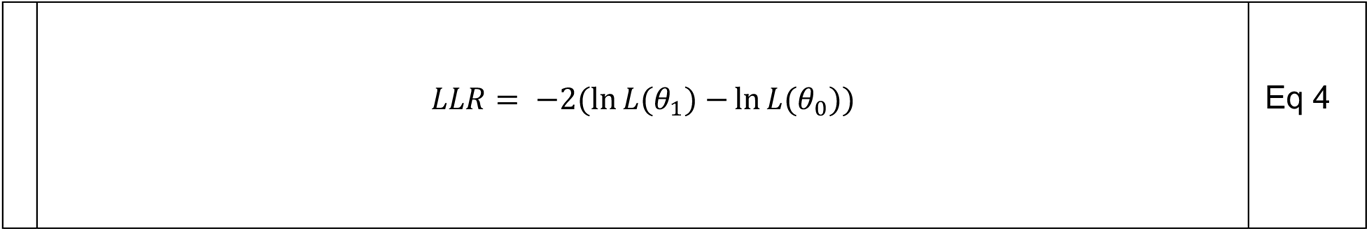

Where *L*(*ϑ*_1_) is the likelihood of the data given the parameters of the alternative hypothesis (biexponential distribution) and *L*(*ϑ*_0_) is the likelihood of the data given the parameters of the null hypothesis (exponential distribution). Values are reported as the MLE of the parameter ± standard error. Here, standard error is calculated from the 95% confidence interval obtained from the *mle* function in MATLAB using the Wald method.

Dwell time distributions are visualized as either their cumulative probabilities, violin plots, or histograms. Cumulative probability plots were constructed by computing a cumulative distribution function (CDF) estimate (Eq X) where the value of each bin (*v_i_*) is computed by Eq 5.

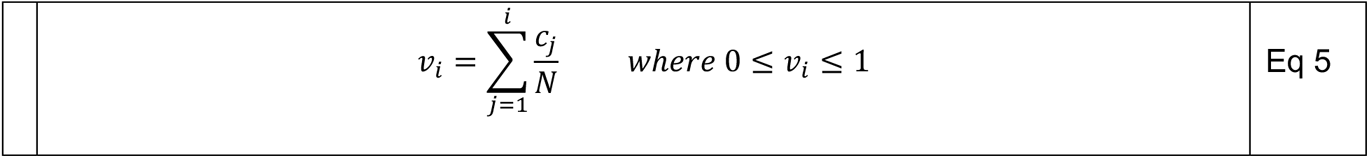

Overlays of MLE of mono- and biexponential distributions were computed by integrating PDF1 and PDF2 over a range [*t_min_, t_max_*] as provided by Eq 6 and 7.

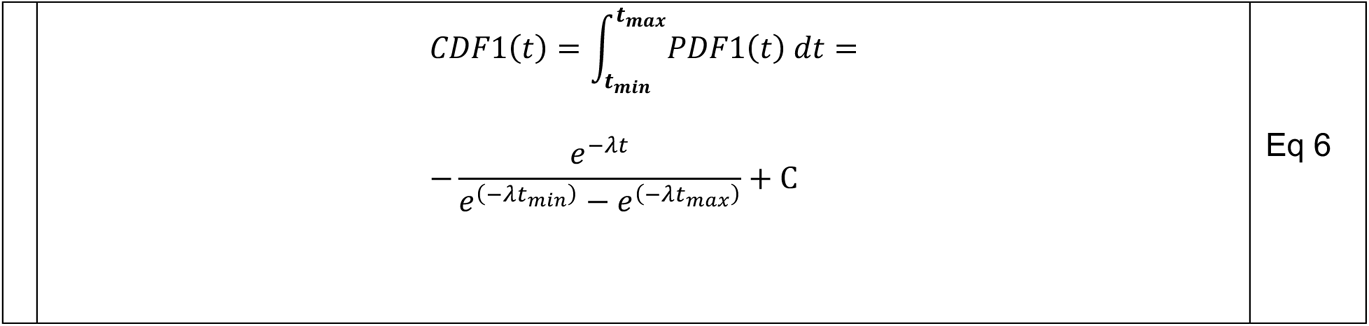

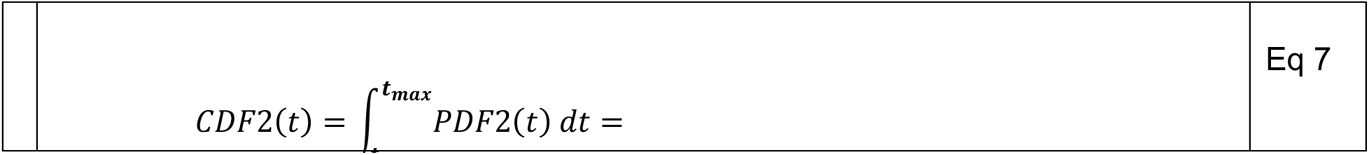

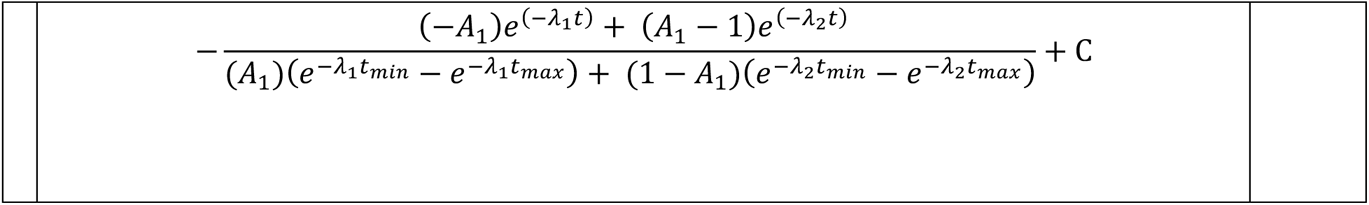

In both cases, *C* was computed as either *CDF*1(*t_min_*) or *CDF*2(*t_min_*) which returns values in the range [0,1]. In the case of histograms, binned dwell times are overlaid with MLE estimates using PDF values computed from Eq 2 and Eq 3 and scaled for the number of observations in the distribution. For data collected at equilibrium (*e.g.*, 9bp-1A), all unbound and bound events considered for MLE. For non-equilibrium data (*e.g.*, 9bp, **Figure 4b-c**), only the initial unbound event (aka “time to first binding”) was considered for MLE estimation to reduce the potential bias introduced by photobleaching of tight binders.

### Global HMM modeling of single molecule data

Kinetic modeling of single molecule data was performed with QuB^56^. Hidden Markov models were globally optimized to simultaneously describe the idealized behavior across all molecules within a particular dataset. Contrary to MLE analysis of single-molecule dwell time distributions, HMMs enable hypothesis testing of state connectivity and globally optimize transitions rates across all molecules for a model postulated *a priori*. Given the complexity and scale of our CoSMoS data collected with the 9bp-1A 5’SS RNA, we tested specific models on the following four subsets of data: (1) varying 9bp-1A concentration at 0 nM U1-C in DMSO, (2) varying 9bp-1A concentration at 100 nM U1-C in DMSO, (3) varying U1-C concentration at 1 nM 9bp-1A in DMSO, and (4) varying branaplam concentration at 1 nM 9bp-1A with 100 nM U1-C (**Fig. S26, Table S3**). In the latter case, 10 µM branaplam data was excluded from QuB modeling due to our inability to fully sample the bound dwell time distribution (**Figure S9F**). Each dataset was used to optimize specific rate parameters, the results of which are reported in **Table 1**. Transition rates of user-defined models were optimized using maximum idealized point (MIP) likelihood rate estimation^57^. In all cases, a “dead time” parameter in QuB was set equal to the sampling time. For each condition, multiple models of varying complexity were built and ranked by their Bayesian Information Criterion^58^ (Eq 8)

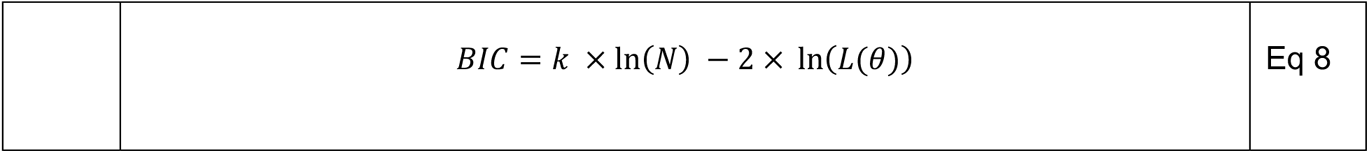

where *k* is the number of free parameters, *N* is the number of data points (frames), and *L*(*ϑ*) is the likelihood of the data given the model returned by MIP. Optimized rate constants are provided in **Supplementary Table 3**.

To confirm this model matched our experimental data, we performed single molecule simulations (**Figure S27-31**). Single molecule trajectories were simulated as noiseless transitions between discrete states following the global model of 9bp-1A binding under Markov assumptions (**Figure 6**, **Table 1**). Each simulation consisted of 10,000 molecules for a given concentration of 9bp-1A, U1-C, and branaplam. Five percent of these molecules were simulated without the inclusion of U1-C, to mimic of single molecule observations (**Figure S11**). Each molecule was simulated for 600 frames at the frame rate used in data collection (**Table S2**).

## Key Reagents

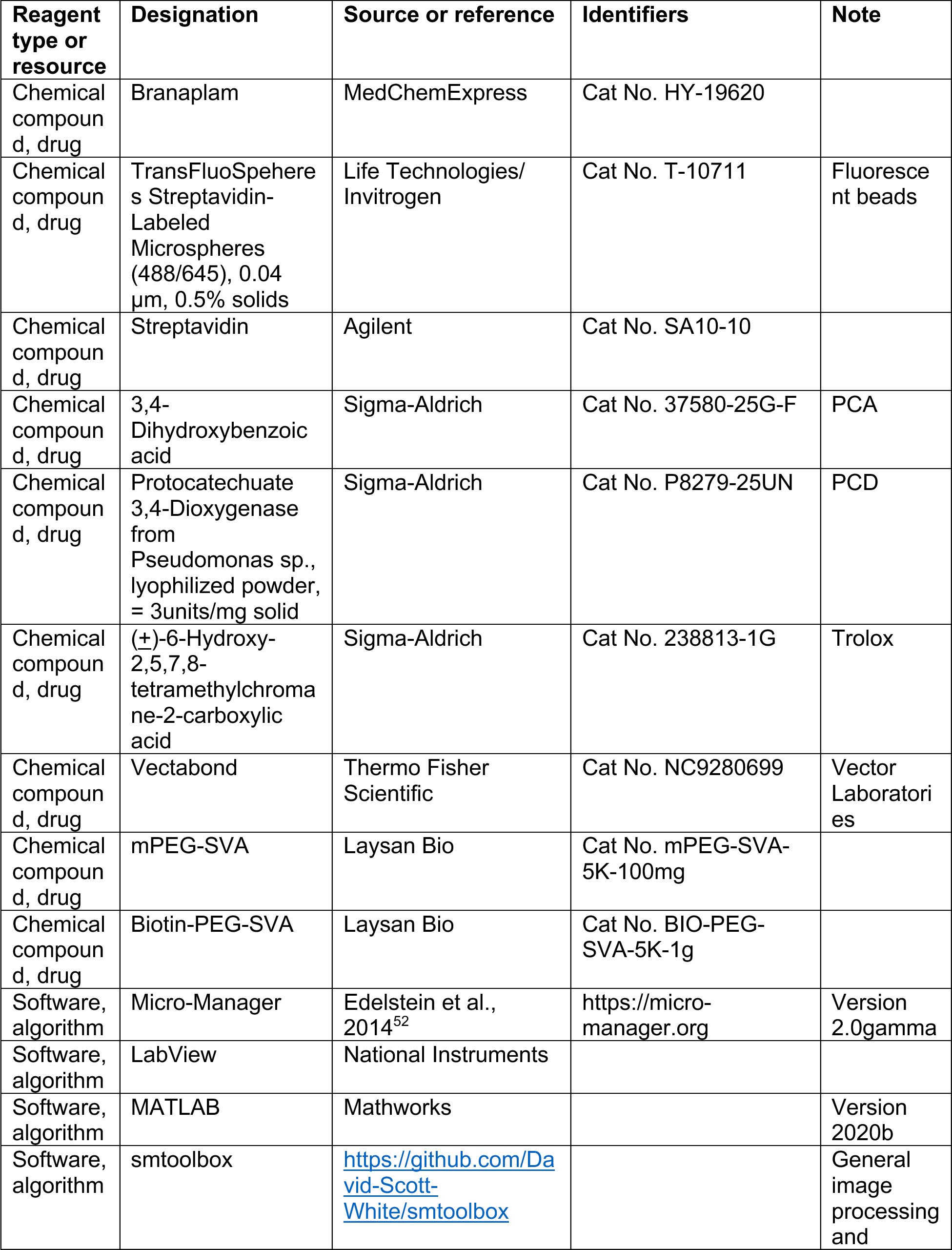
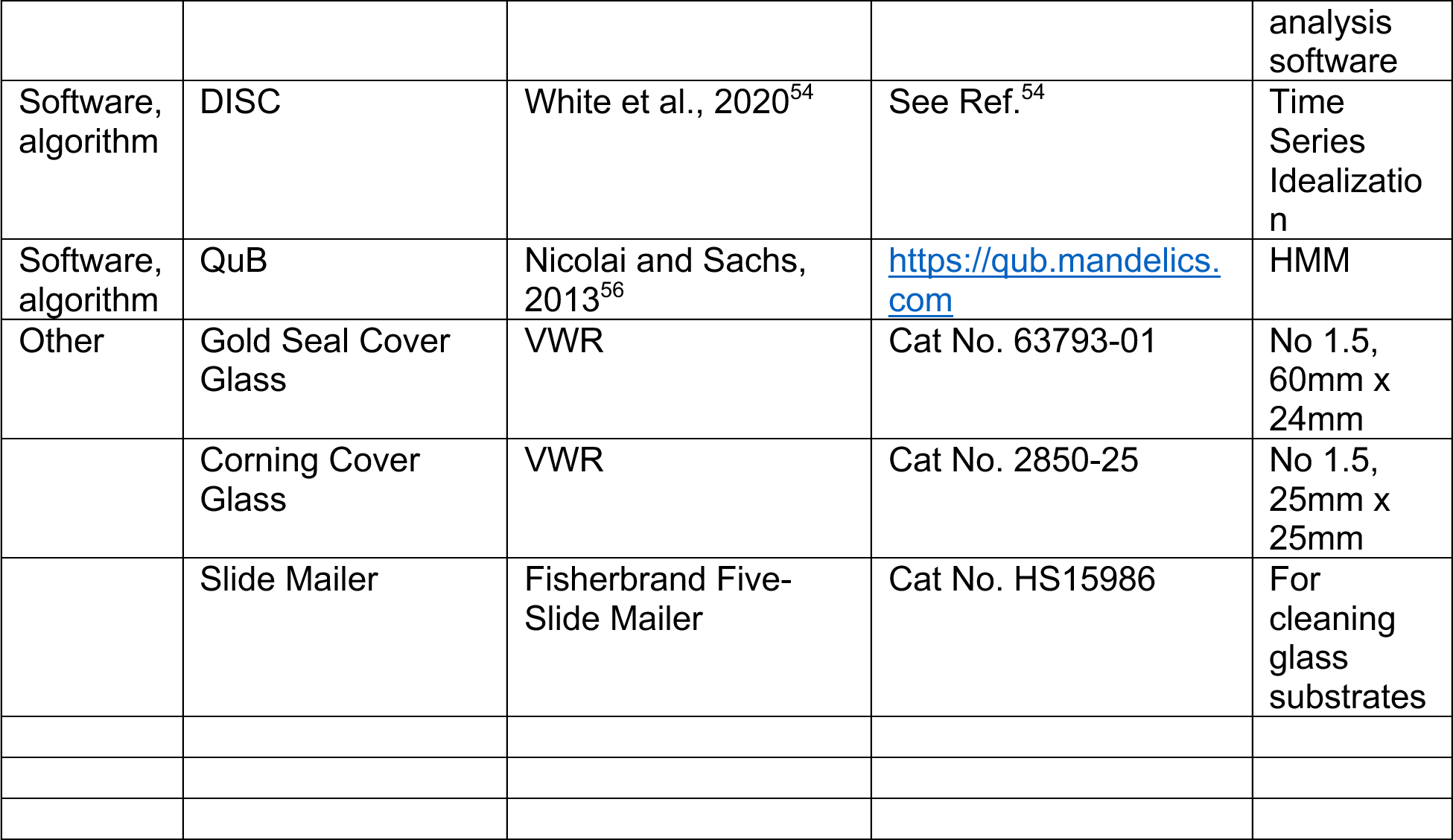

## Data Availability

Data will be made available through Zenodo upon publication.

## Code Availability

Our general-purpose single molecule analysis software smtoolbox is available at [github to smtoolbox]. Custom MATLAB scripts for analysis specific to these data is available at [github to repo for this analysis].

## ACKNOWLEDGEMENTS

This work was supported by funding from Remix Therapeutics, grants from the National Institutes of Health (R35 GM136261 to AAH) with additional support from a Research Forward grant award from the Wisconsin Alumni Research Foundation and a NIH postdoctoral fellowship award (F32 GM143780 to DSW). We thank Nathalie Drouin, Mélissa Morin, Myriam Létourneau, and Amira Yazidi at NMX Research Solutions for assistance with protein production, Maria McGresham at August Bioservices for assistance with SPR experiments, and Maximilian Plach at 2bind GmbH for assistance with MST experiments. We thank Jeff Gelles, David Bentley, Sam Butcher, and members of the Hoskins and Herschlag labs for helpful discussions.

