## Supplemental Figures and Tables for "A Sequential Binding Mechanism for 5’ Splice Site Recognition and Modulation for the Human U1 snRNP"

**Affiliations:**
[1] Department of Biochemistry, University of Wisconsin-Madison, Madison, WI
[2] Remix Therapeutics, Watertown, MA
[3] Department of Chemistry, University of Wisconsin-Madison, Madison, WI
[4] Present Address: Element Biosciences, San Diego, CA

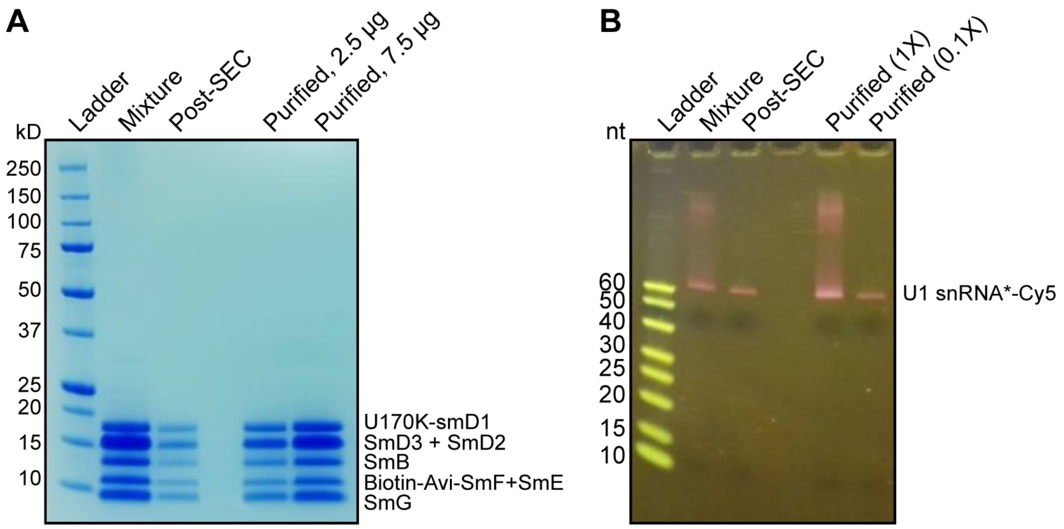


**Figure S1. In vitro reconstitution of minimal U1 snRNP.** (**A**) Protein components of *in vitro* reconstituted U1 snRNP after SEC purification visualized on a NuPAGE Bis-Tris (4-12%) SDS gel stained with Coomassie. (**B**) Visualization of the U1 snRNA component of the purified U1 snRNP. The U1 snRNA is labeled with Cy5 (pink color in gel) on a TBE-Urea (15%) gel stained with SpyroGold.

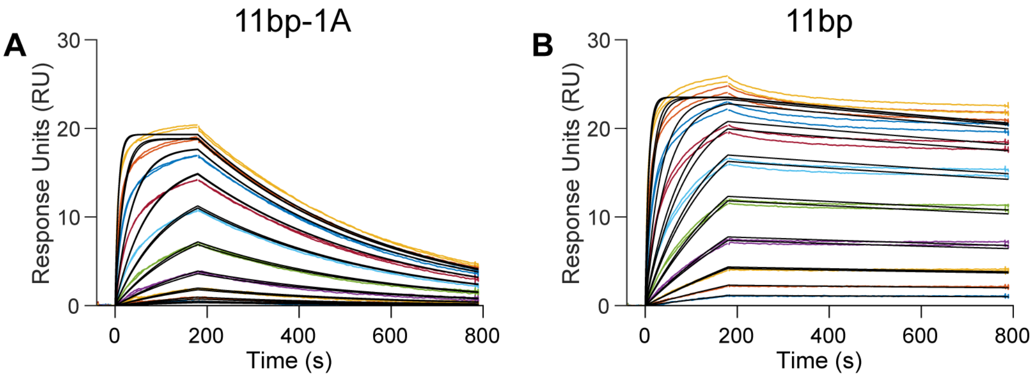


**FigureS2: Single state binding model fits to SPR data of 11bp and 11bp-1A.** SPR sensorgrams of (A) 11bp-1A and (**B**) 11bp at variable concentrations of U1 snRNP overlaid with fits from a one-state (black lines) Langmuir binding model.

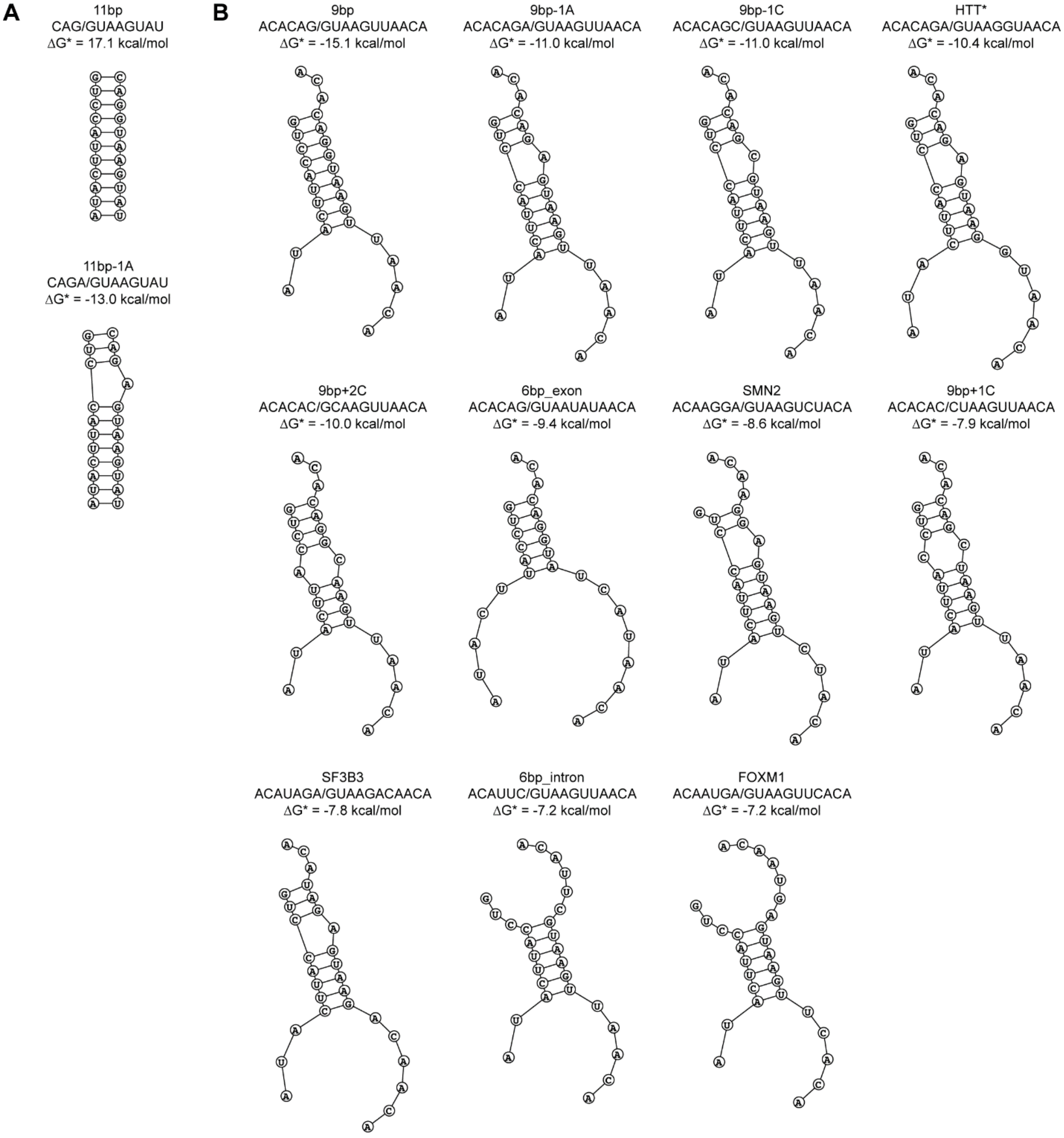


**Figure S3. Predicted bimolecular interaction between 5'SS oligos and U1 snRNA.** (**A**) Oligomers used in SPR and MST assays. (**B**) Oligomers used in CoSMoS. For both **A** and **B**, the bimolecular secondary structure and the ΔG* value correspond to the minimum free energy predictions from RNAstructure Web Server (Reuter & Mathews, 2010).

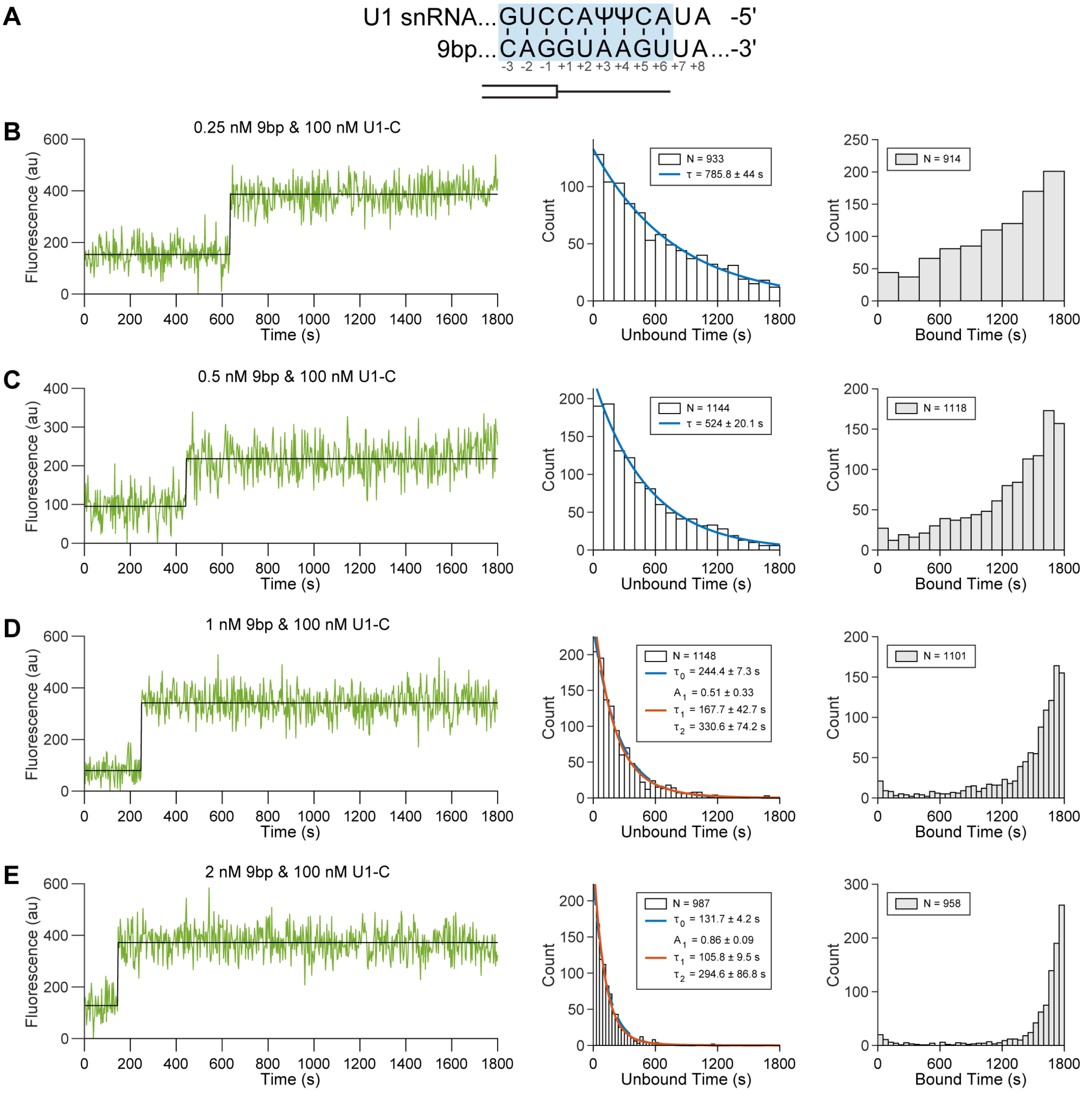


**Figure S4. CoSMoS of the 9bp RNA oligo in presence of 100 nM U1-C.** (**A**) Predicted duplex between U1 snRNA SSRS and the 9bp RNA oligo. (**B-E**) (Left) Representative time series with idealization (black line), (middle) unbound dwell time distributions, and (right) bound dwell time distributions across 9bp RNA oligo concentrations (0.25 nM, 0.5 nM, 1 nM, 2 nM) with 100 nM U1-C. Dwell times are overlaid with expectations from MLE (red and blue lines).

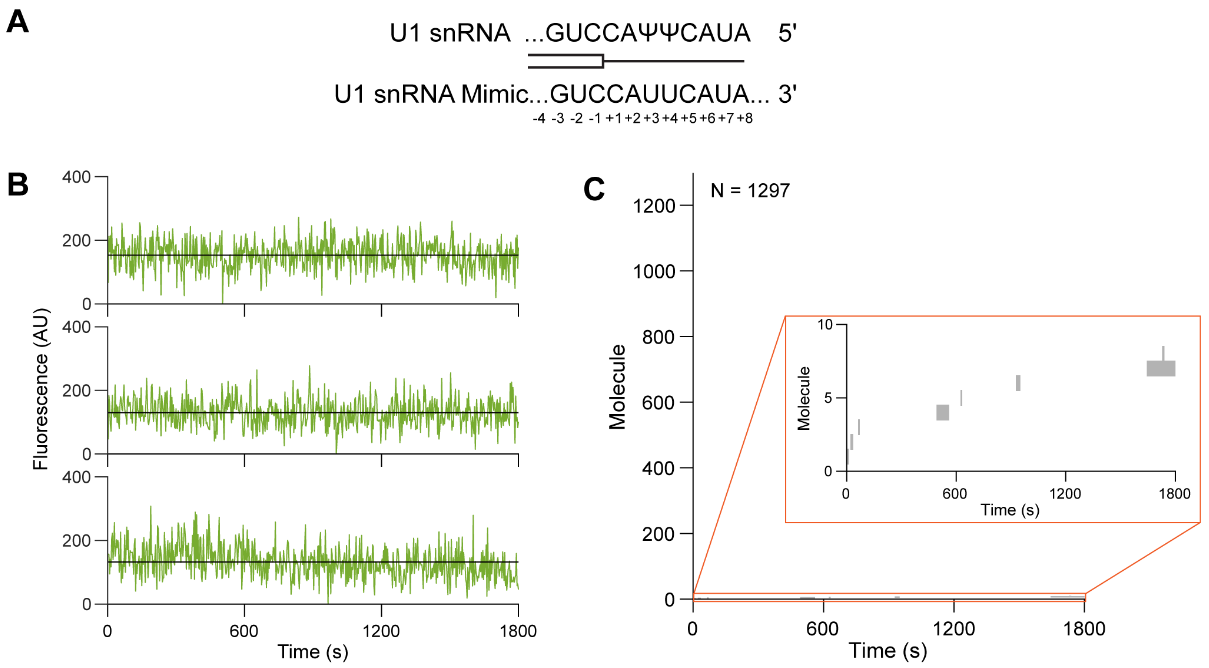


**Figure S5. A control oligo shows little stable association with the U1 snRNP.** (**A**) Alignment of the U1 snRNA SSRS and control oligo (no binding predicted). (**B**) Representative traces with idealization (black line) of 10 nM control oligo with 100 nM U1-C. (**C)** Rastergram summarizing the detected binding events across all molecules over time (binding events in grey), sorted by time of first binding. Data presented are a combination of two replicates (replicate 1: *N* = 736 immobilized U1 molecules; replicate 2: *N* = 561 immobilized U1 molecules) (Inset) Truncation of the y-axis to zoom in on the observed co-localized, control oligo binding events.

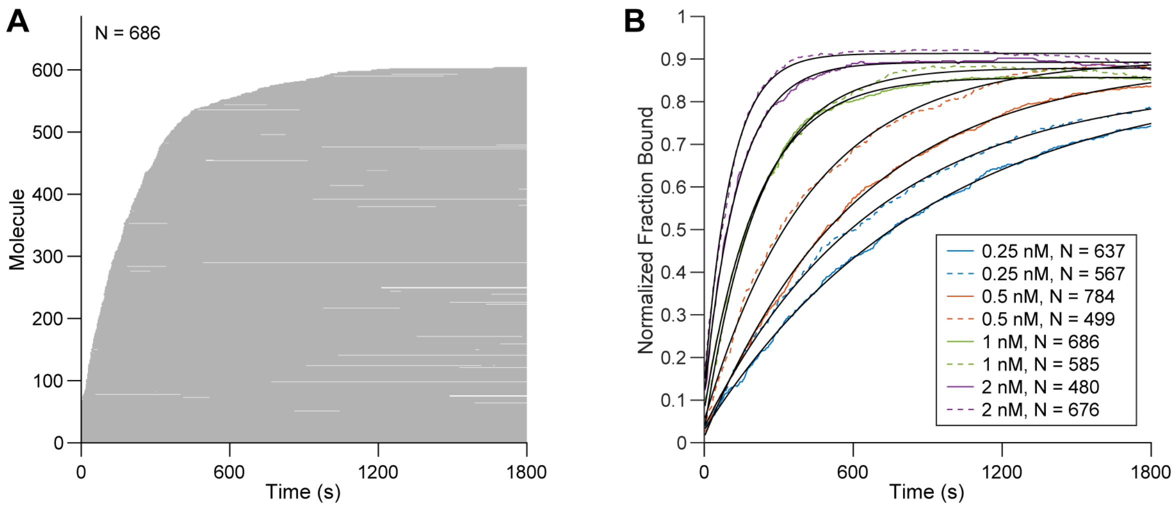


**Figure S6. Non-equilibrium kinetics of the 9bp RNA oligo.** (**A**) Rastergram summarizing the detected binding events across all molecules over time (binding events in grey) of 1 nM 9bp RNA oligo in the presence of 100 nM U1-C (**B**) Observed fraction bound (normalized to total number of immobilized U1 molecules, indicated by *N* in legend) overlaid with a single exponential function with a time constant of $k_{eq}$ (black). Replicates are shown as solid and dashed lines of the same color.

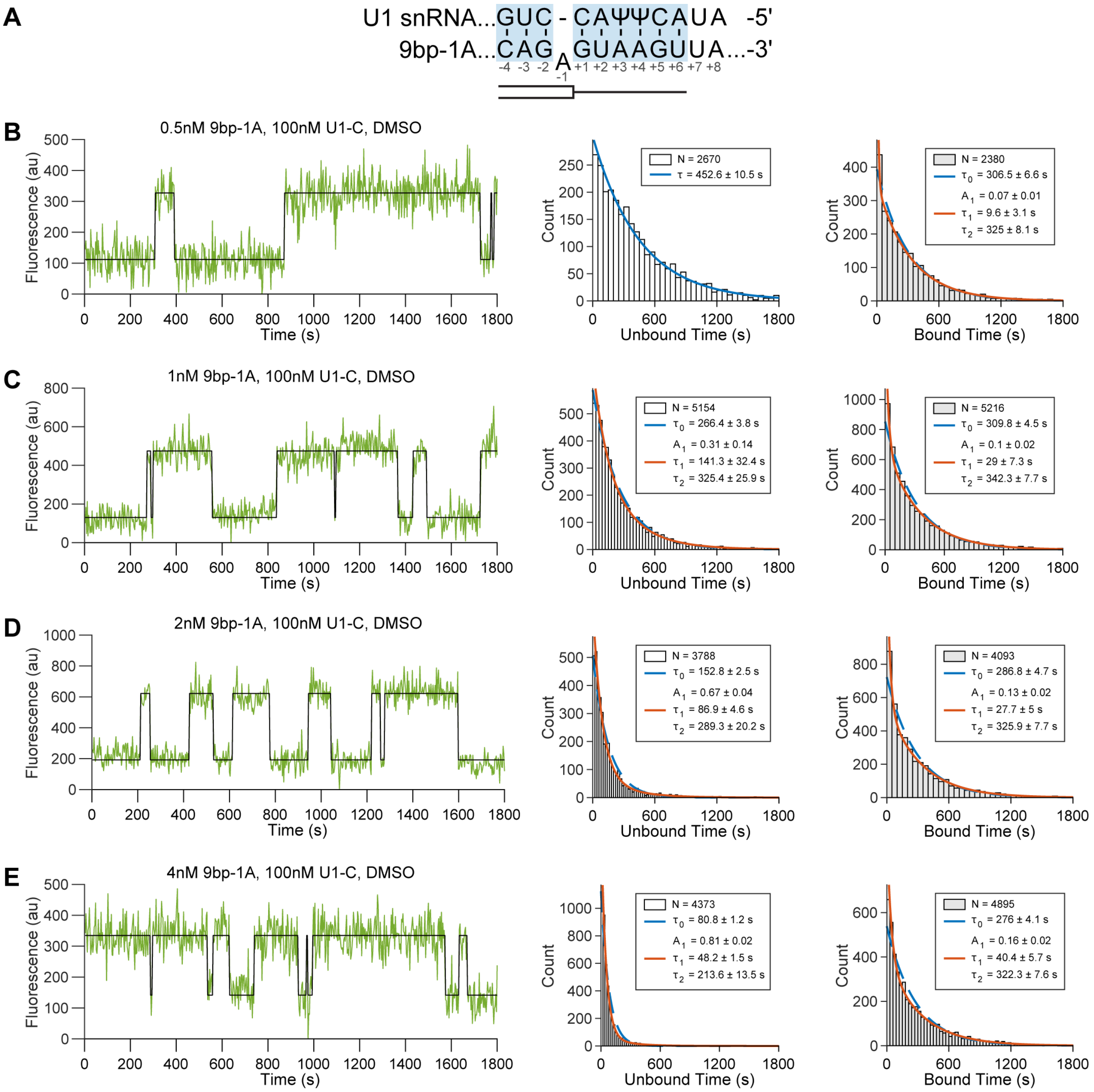


**Figure S7. CoSMoS of 9bp-1A RNA oligo in presence of 100 nM U1-C.** (**A**) Predicted duplex between U1 snRNA and 9bp-1A RNA oligo. (**B-E**) (Left) Representative time series with idealization (black line), (middle) unbound dwell time distributions, and (right) bound dwell time distributions across 9bp-1A concentrations (0.5 nM, 1 nM, 2 nM, 4 nM) with 100 nM U1-C. Dwell times are overlaid with expectations from MLE (blue and red lines).

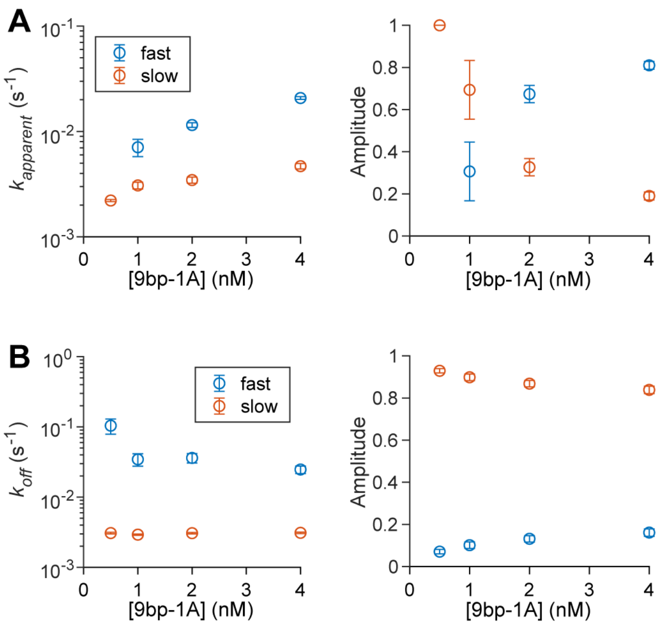


**Figure S8. Maximum likelihood estimates of 9bp-1A RNA oligo unbound and bound dwell time distributions.** (**A**) MLE of a biexponential distribution $k_{app}$ values and associated amplitudes from unbound dwell times. (**B**) Maximum likelihood estimations of a biexponential distribution $k_{off}$ values and associated amplitudes from bound dwell times. Errors are standard error.

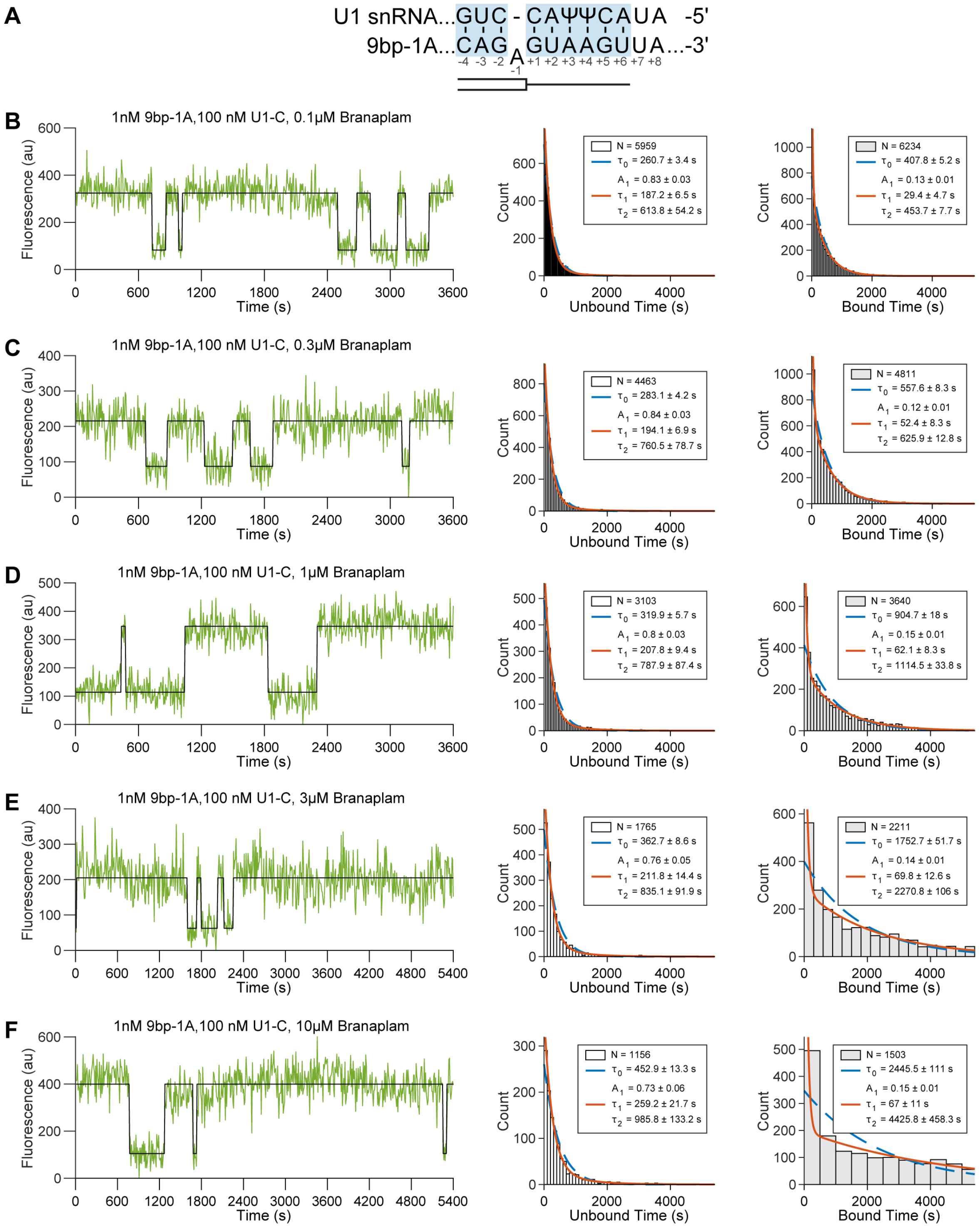


**Figure S9. CoSMoS of 9bp-1A RNA oligo in presence of 100 nM U1-C at various branaplam concentrations.** (**A**) Predicted duplex between U1 snRNA and 9bp-1A RNA oligo. (**B-F**) (Left) Representative time series with idealization (black line), (middle) unbound dwell time distributions, and (right) bound dwell time distributions across branaplam concentrations (0.1 μM, 0.3 μM, 1 μM, 3 μM, and 10 μM) at 1 nM 9bp-1A and 100 nM U1-C. Dwell times are overlaid with expectations from MLE (blue and red lines).

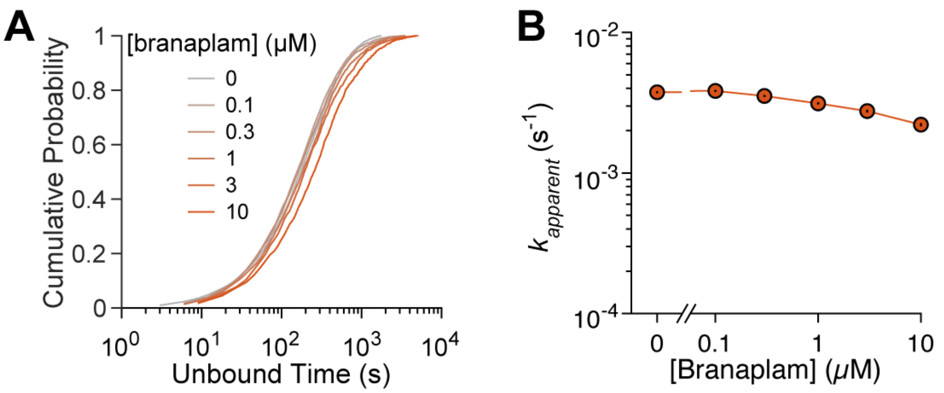


**Figure S10. Effect of branaplam on 9bp-1A RNA oligo association rate.** (**A**) Cumulative probability of unbound dwell times across branaplam concentrations (0 µM, *N* = 5154; 0.1 µM, *N* = 5959; 0.3 µM, *N* = 4463; 1 µM, *N* = 3103; 3 µM, *N* = 1765; 10 µM, *N* = 1156. (**B**) Average association rate ($k_a$) determined for MLE of single exponential distributions at each branaplam concentration.

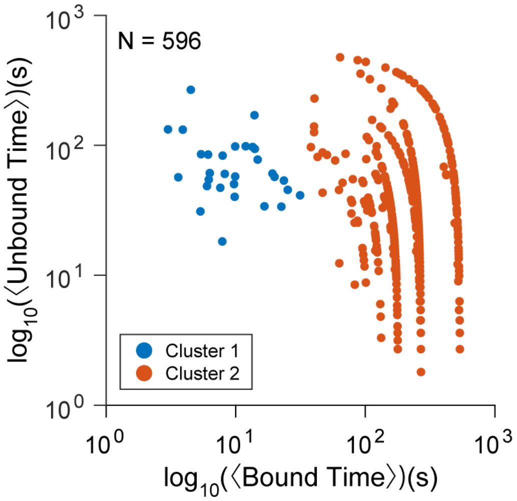


**Figure S11. Clustering of average dynamics at 10 μM branaplam.** Results of hierarchical clustering of the average bound and unbound times across all molecules collected under 1 nM 9bp-1A, 100 nM U1-C, and 10 μM branaplam. Cluster 1 exhibits faster average dynamics and accounts for 5% of the data.

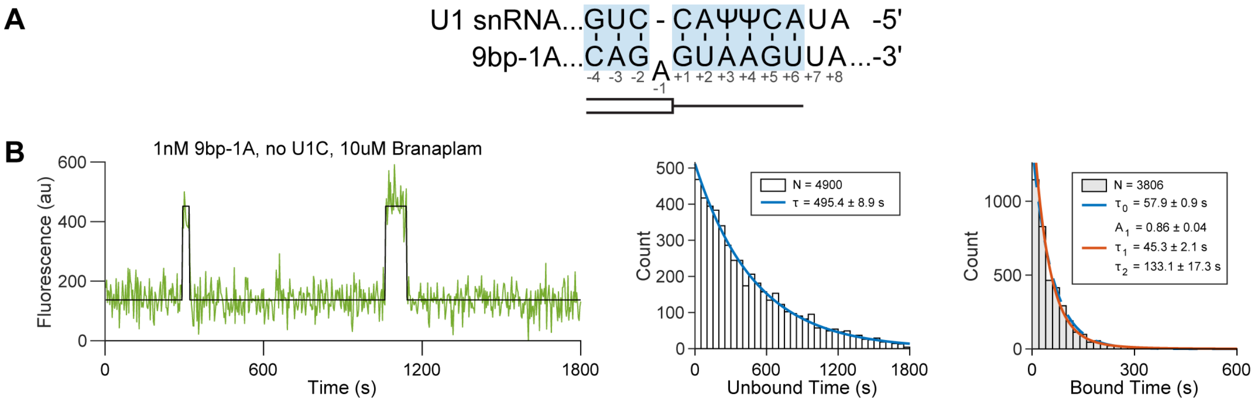


**Figure S12. CoSMoS of 9bp-1A RNA oligo in absence of U1-C with 10 μM branaplam.** (**A**) Predicted duplex between U1 snRNA SSRS and 9bp-1A RNA oligo. (**B)** (Left) Representative time series with idealization (black line), (middle) unbound dwell time distributions, and (right) bound dwell time distributions at 1 nM 9bp-1A, 10 μM branaplam, and no U1-C. Dwell times are overlaid with expectations from MLE (blue and red lines).

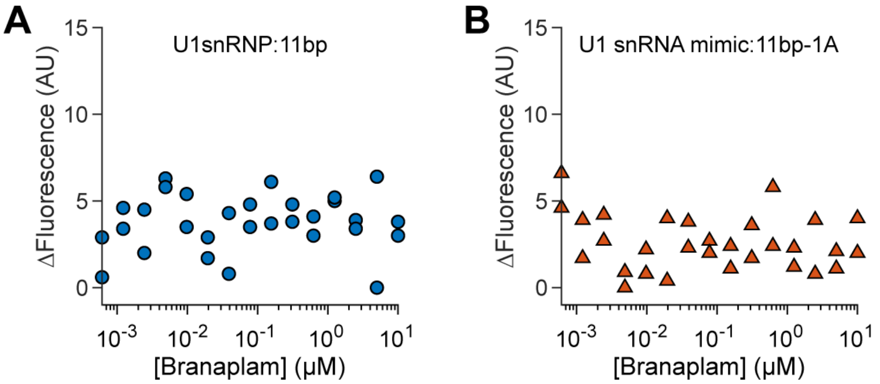


**Figure S13. MST measurements of branaplam binding.** (**A**). Change in fluorescence of a pre-formed U1 snRNP:11bp RNA oligo complex without U1-C in response to branaplam. (**B**) Change in fluorescence of a pre-formed U1 snRNA:11bp-1A RNA:RNA duplex response to branaplam.

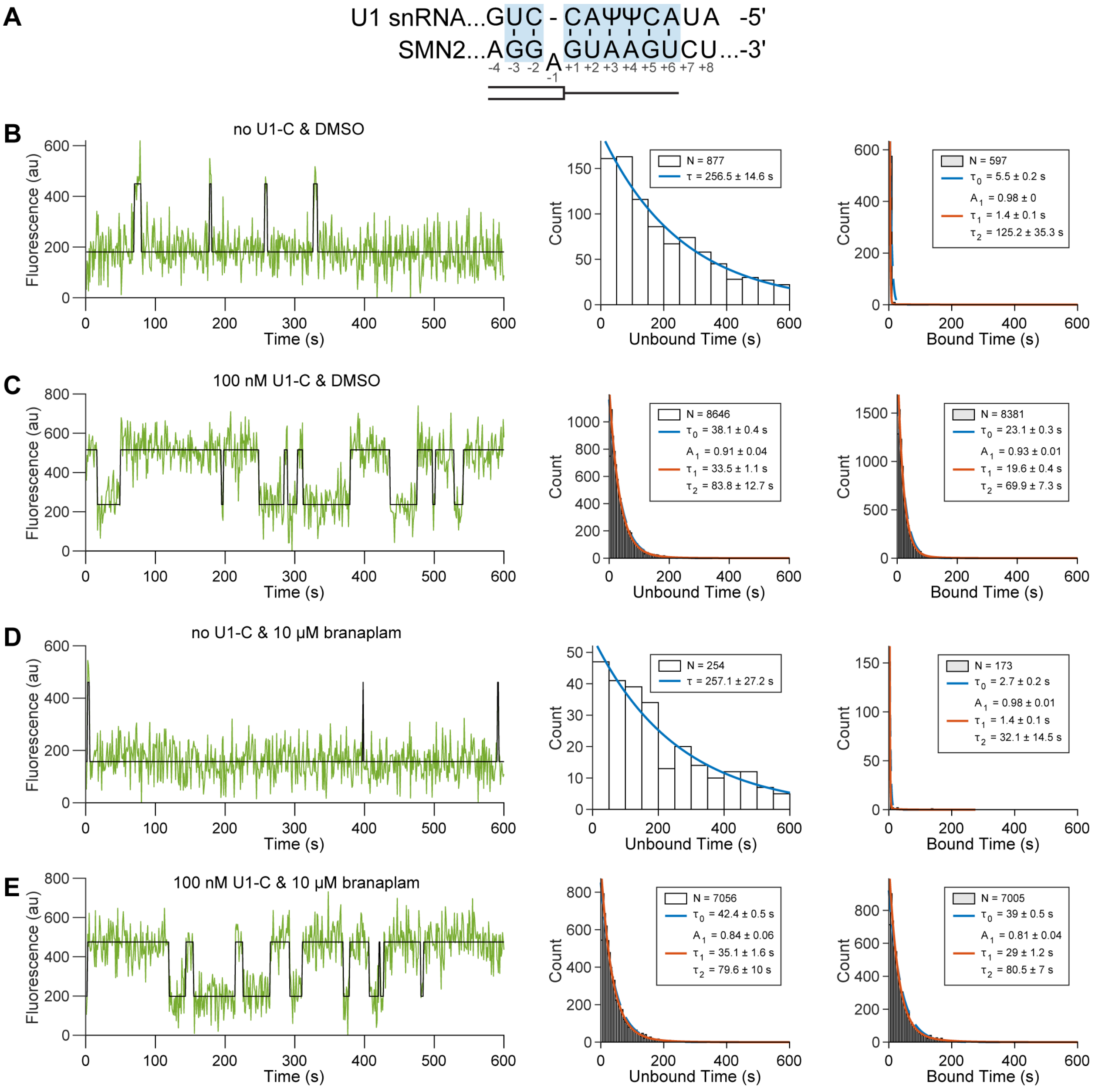


**Figure S14. Interaction of U1-C and branaplam on SMN2 RNA oligo binding.** (**A**) Predicted duplex between U1 snRNA and SMN2 RNA oligo. (**B-E**) (Left) Representative time series with idealization (black line), (middle) unbound dwell time distributions, and (right) bound dwell time distributions with (100 nM) or without U1-C and with (10 μM) or without (DMSO) branaplam and 10 nM SMN2. Dwell times are overlaid with expectations from MLE (blue and red lines).

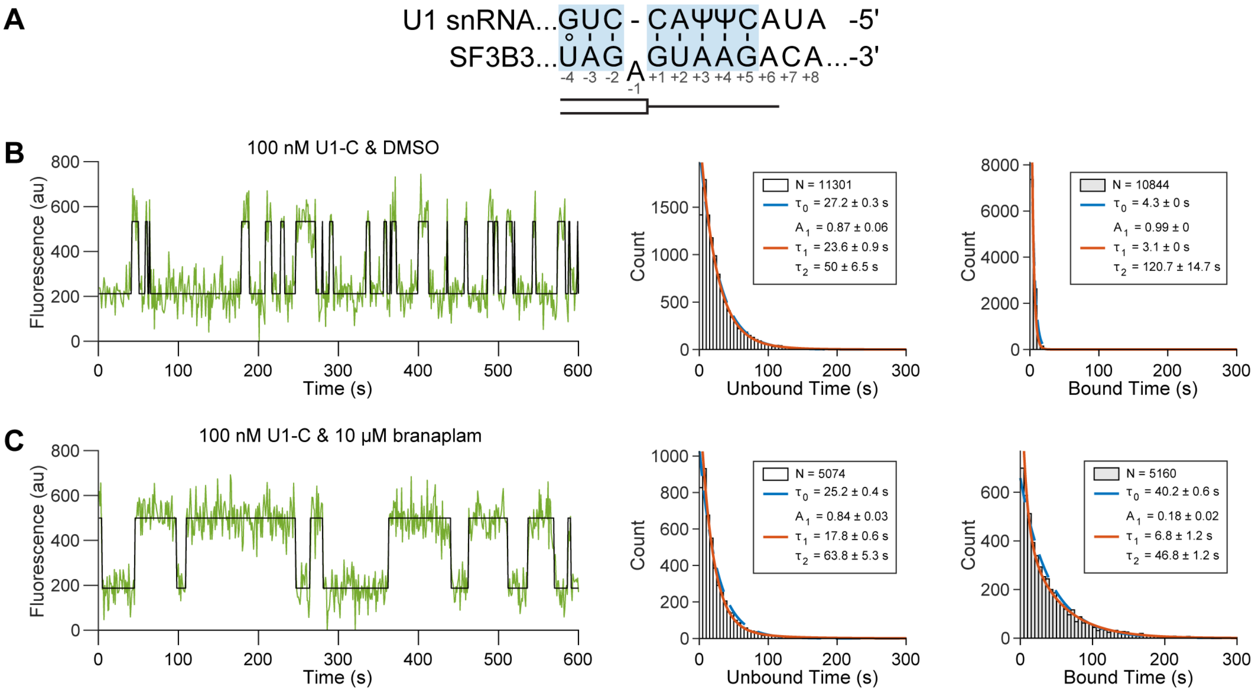


**Figure S15. Interaction of U1-C and branaplam on SF3B3 RNA oligo binding.** (**A**) Predicted duplex between U1 snRNA and SF3B3 RNA oligo. Circle indicates a wobble base pair. (**B-C**) (Left) Representative time series with idealization (black line), (middle) unbound dwell time distributions, and (right) bound dwell time distributions with (100 nM) or without U1-C and with (10 μM) or without (DMSO) branaplam and 10 nM SF3B3. Dwell times are overlaid with expectations from MLE (blue and red lines).

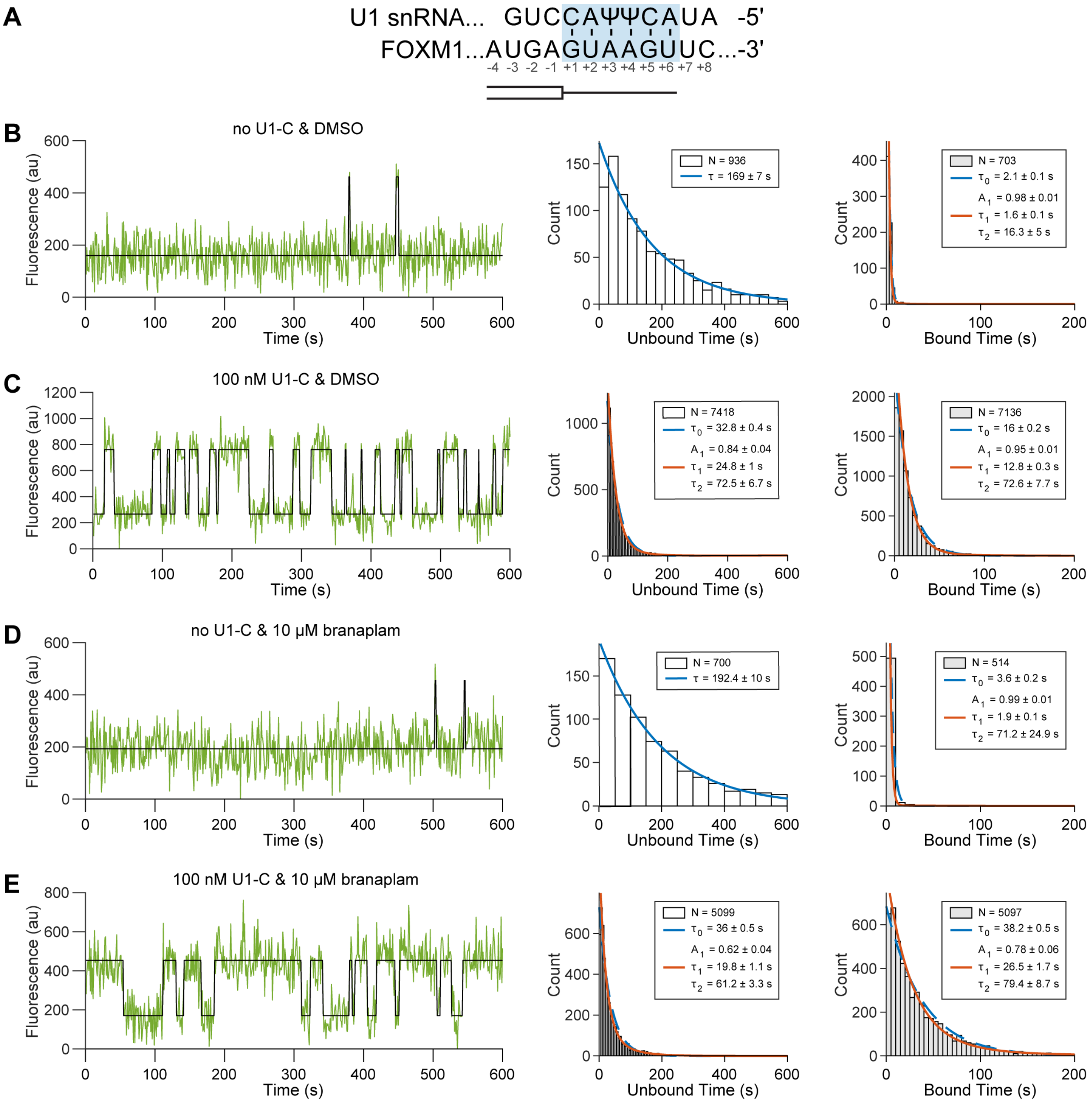


**Figure S16. Interaction of U1-C and branaplam on FOXM1 RNA oligo binding.** (**A**) Predicted duplex between U1 snRNA and FOXM1 RNA oligo. (**B-E**) (Left) Representative time series with idealization (black line), (middle) unbound dwell time distributions, and (right) bound dwell time distributions with (100 nM) or without U1-C and with (10 μM) or without (DMSO) branaplam and 10 nM FOXM1. Dwell times are overlaid with expectations from MLE (blue and red lines).

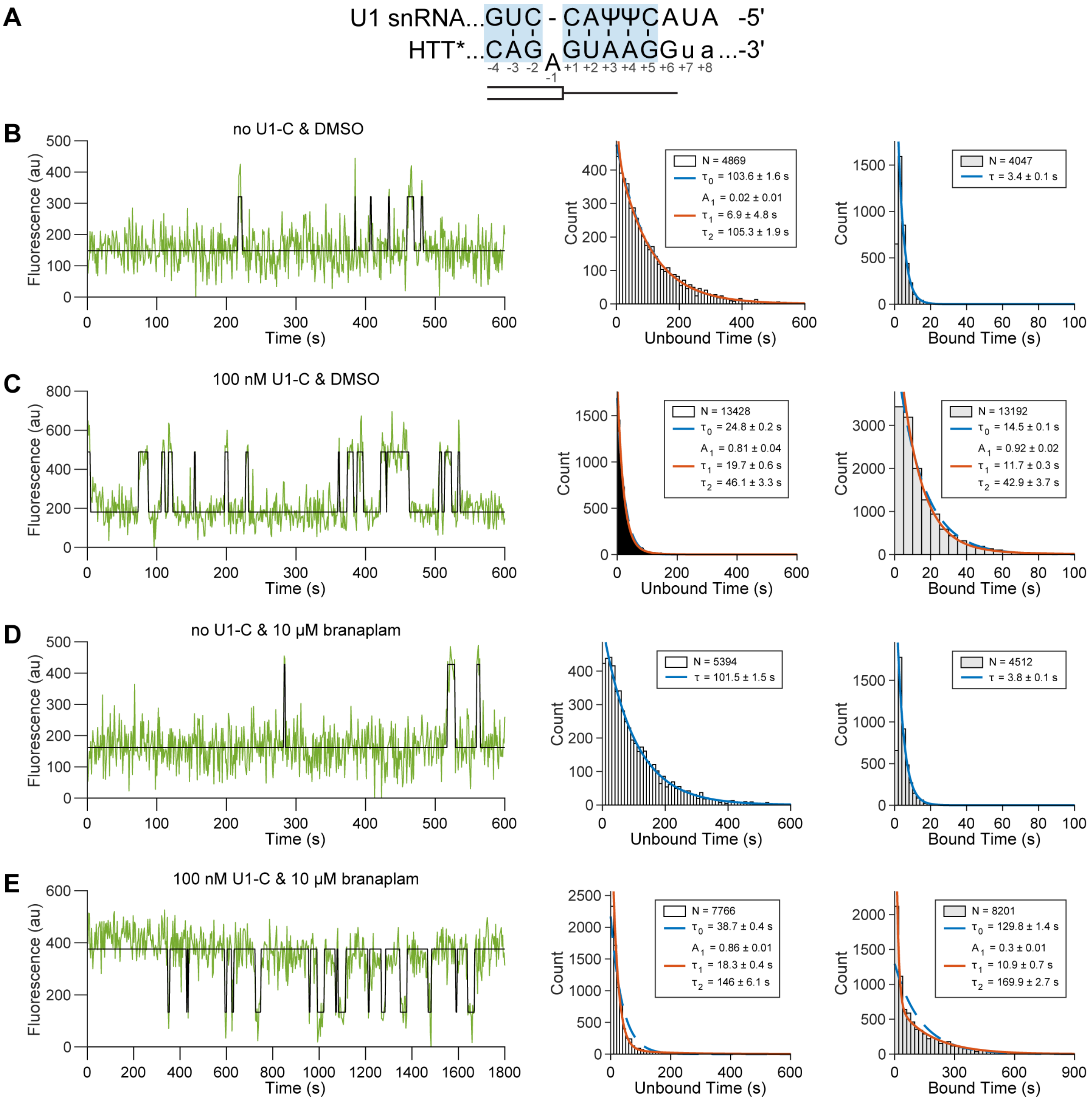


**Figure S17. Interaction of U1-C and branaplam on HTT* RNA oligo binding.** (**A**) Predicted duplex between U1 snRNA and HTT*. Lowercase letters indicate mutations from wild type needed commercial synthesis (WT: CAGAGUAAGGGG) (**B-E**) (Left) Representative time series with idealization (black line), (middle) unbound dwell time distributions, and (right) bound dwell time distributions with (100 nM) or without U1-C and with (10 μM) or without (DMSO) branaplam and 3 nM HTT*. Dwell times are overlaid with expectations from MLE (blue and red lines).

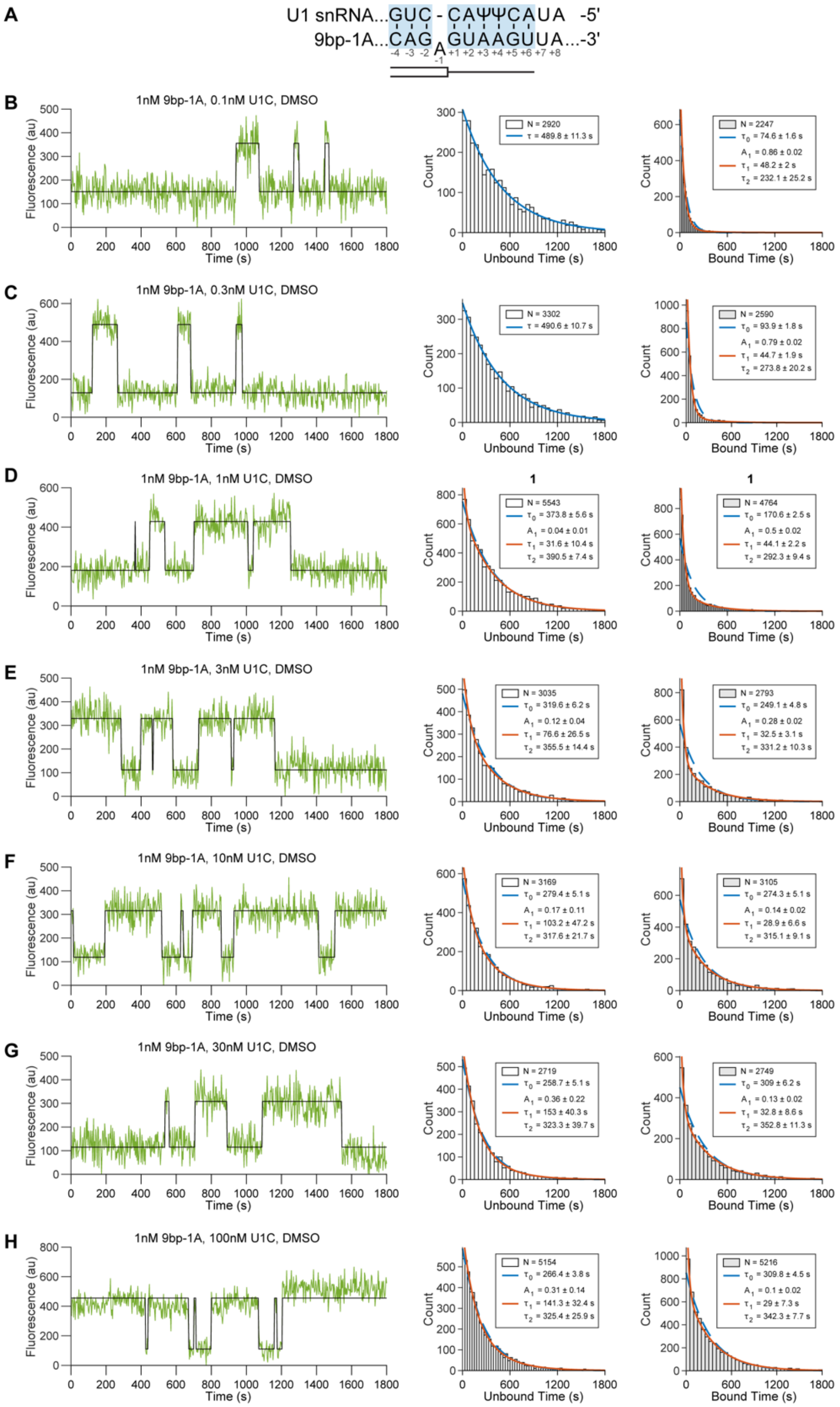


**Figure S18. CoSMoS of 1 nM 9bp-1A RNA oligo across U1-C concentrations.** (**A**) Predicted duplex between U1 snRNA SSRS and 9bp-1A RNA oligo. (**B-H**) (Left) Representative time series with idealization (black line), (middle) unbound dwell time distributions, and (right) bound dwell time distributions across U1-C concentrations (0.1 nM, 0.3 nM, 1 nM, 3 nM, 10 nM, 30 nM, 100 nM) at 1 nM 9bp-1A. Dwell times are overlaid with expectations from MLE (blue and red lines).

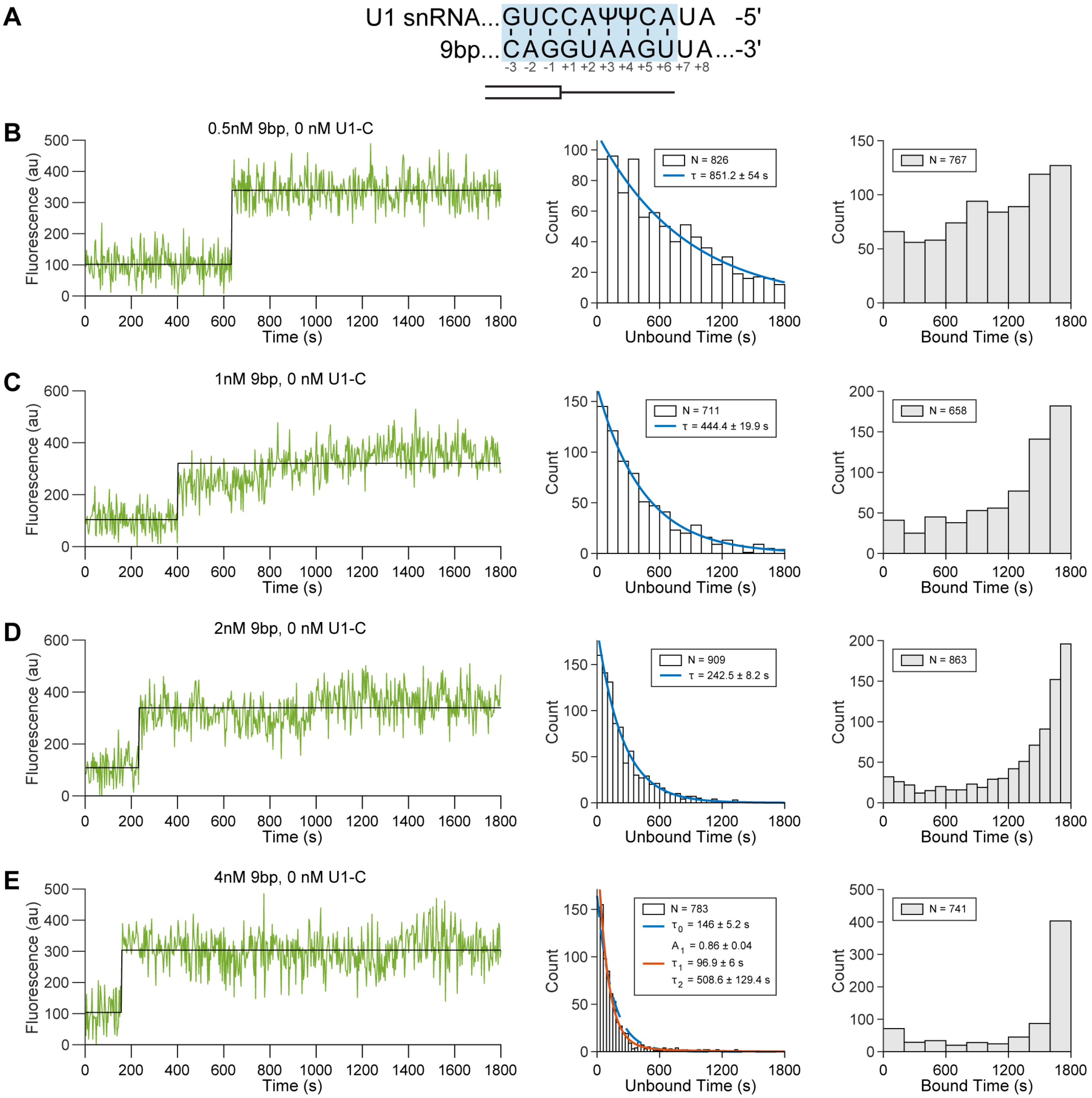


**Figure S19. CoSMoS of 9bp RNA binding in the absence of U1-C.** (**A**) Predicted duplex between U1 snRNA SSRS and 9bp RNA oligo. (**B-E**) (Left) Representative time series with idealization (black line), (middle) unbound dwell time distributions, and (right) bound dwell time distributions across 9bp concentrations (0.5 nM, 1 nM, 2 nM, 4 nM) without U1-C. Dwell times are overlaid with expectations from MLE (blue and red lines).

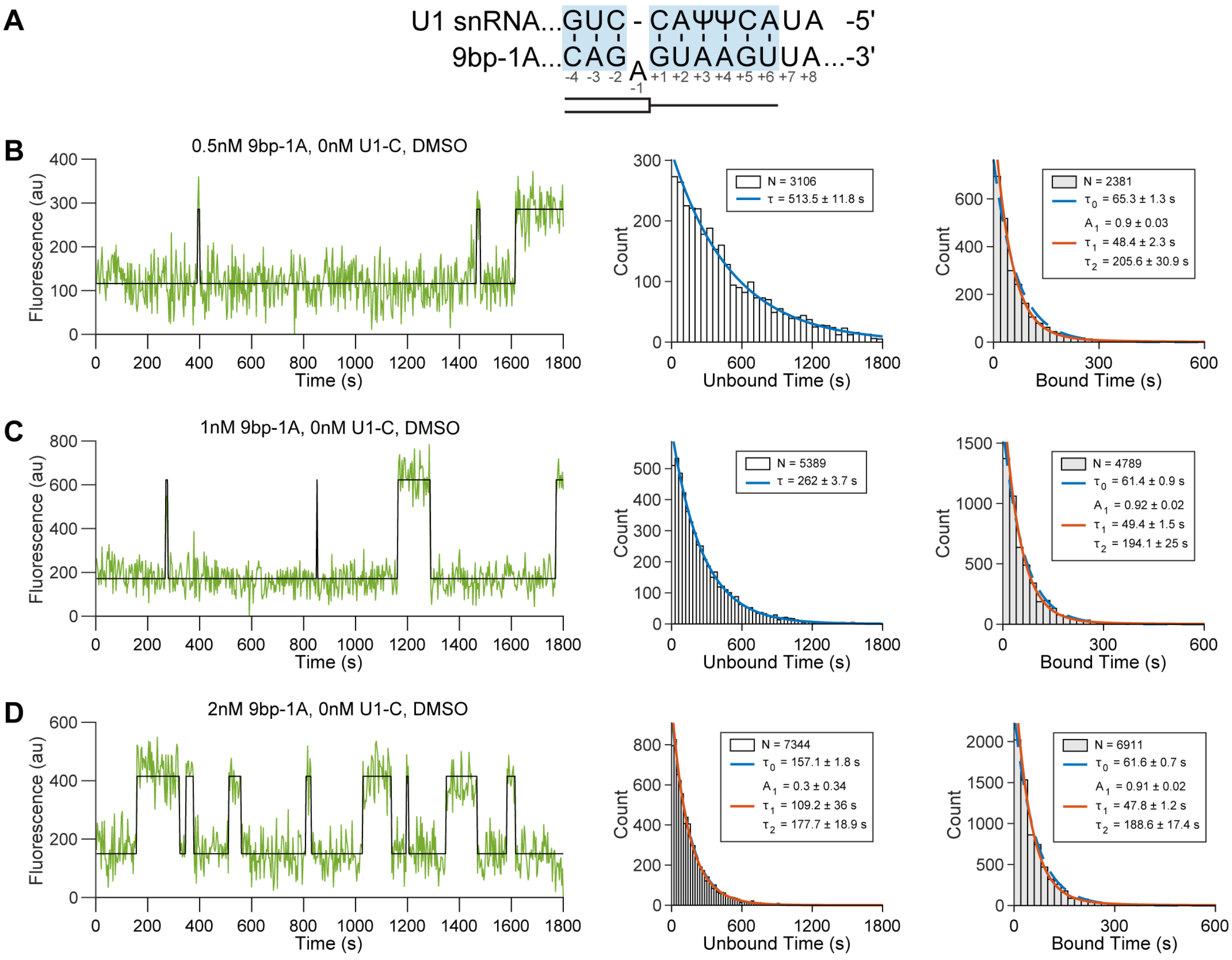


**Figure S20. CoSMoS of 9bp-1A RNA oligo binding in the absence of U1-C.** (**A**) Predicted duplex between U1 snRNA SSRS and 9bp-1A RNA oligo. (**B-D**) (Left) Representative time series with idealization (black line), (middle) unbound dwell time distributions, and (right) bound dwell time distributions across 9bp-1A concentrations (0.5 nM, 1 nM, 2 nM) without U1-C. Dwell times are overlaid with expectations from MLE (blue and red lines).

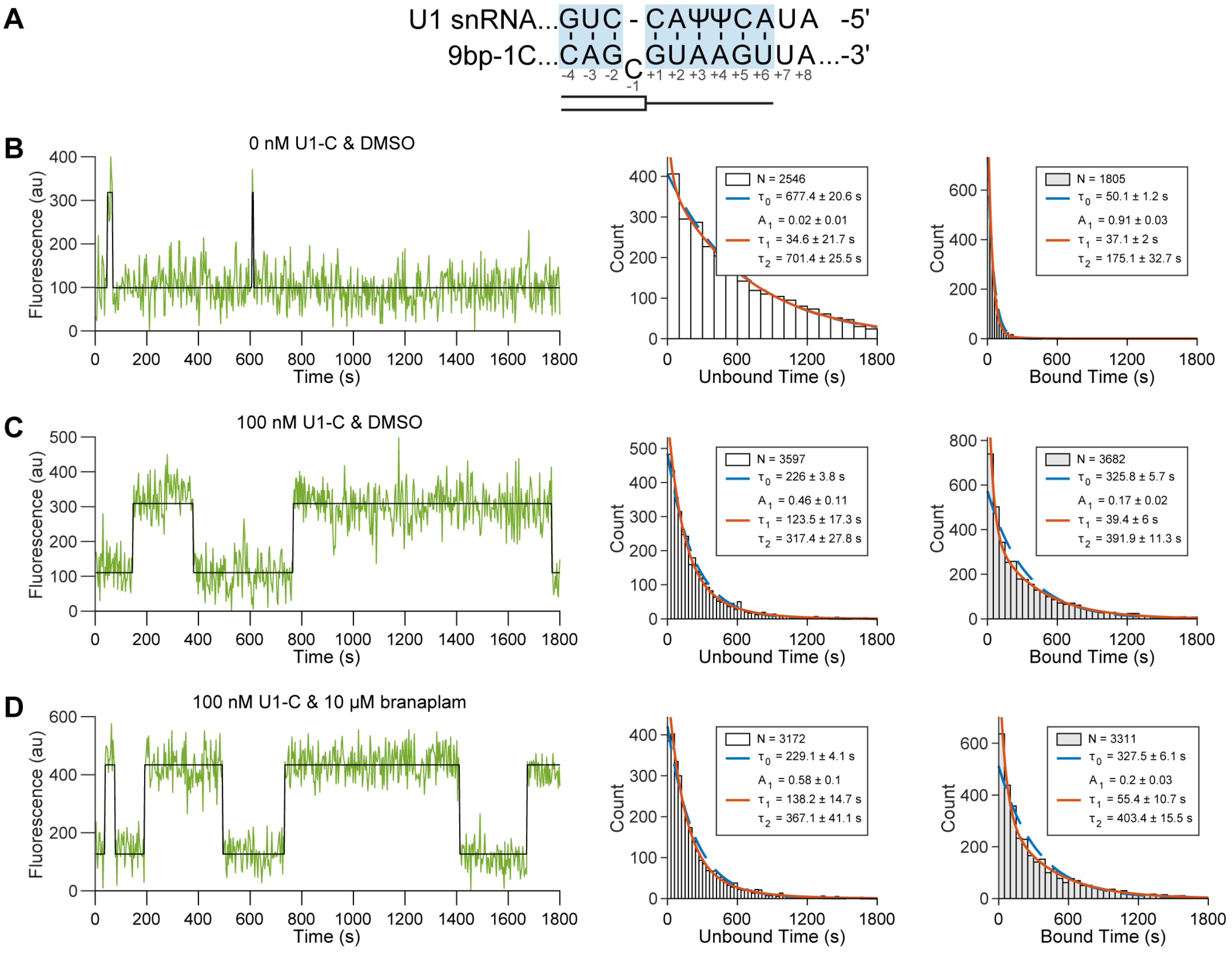

**Figure S21. Impact of U1-C and branaplam on 9bp-1C RNA oligo binding.** (**A**) Predicted duplex between U1 snRNA SSRS and 9bp-1C RNA oligo. (**B-D**) (Left) Representative time series with idealization (black line), (middle) unbound dwell time distributions, and (right) bound dwell time distributions with (100 nM) or without U1-C and with (10 µM) or without (DMSO) branaplam and 1 nM 9bp-1C RNA. Dwell times are overlaid with expectations from MLE (blue and red lines).

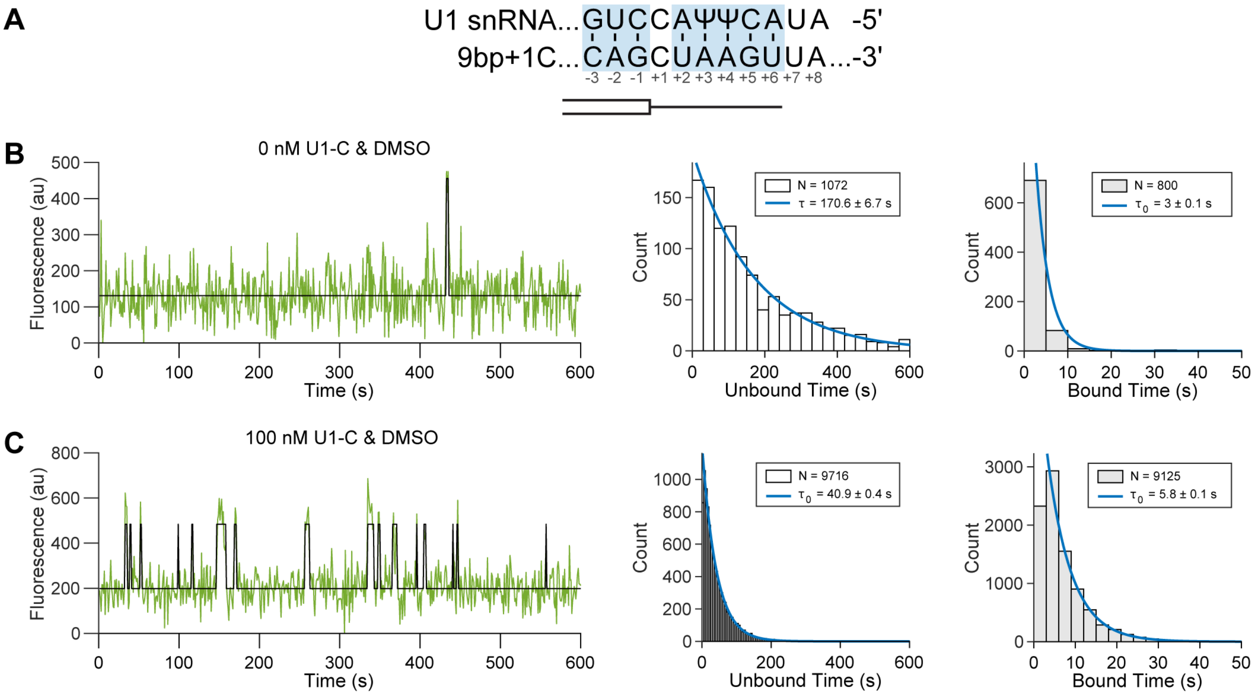


**Figure S22. Effect of U1-C on 9bp+1C RNA oligo binding.** (**A**) Predicted duplex between U1 snRNA and 9bp+1C RNA oligo. (**B-D**) (Left) Representative time series with idealization (black line), (middle) unbound dwell time distributions, and (right) bound dwell time distributions with (100 nM) or without U1-C at 10nM 9bp+1C RNA oligo. Dwell times are overlaid with expectations from MLE (blue lines).

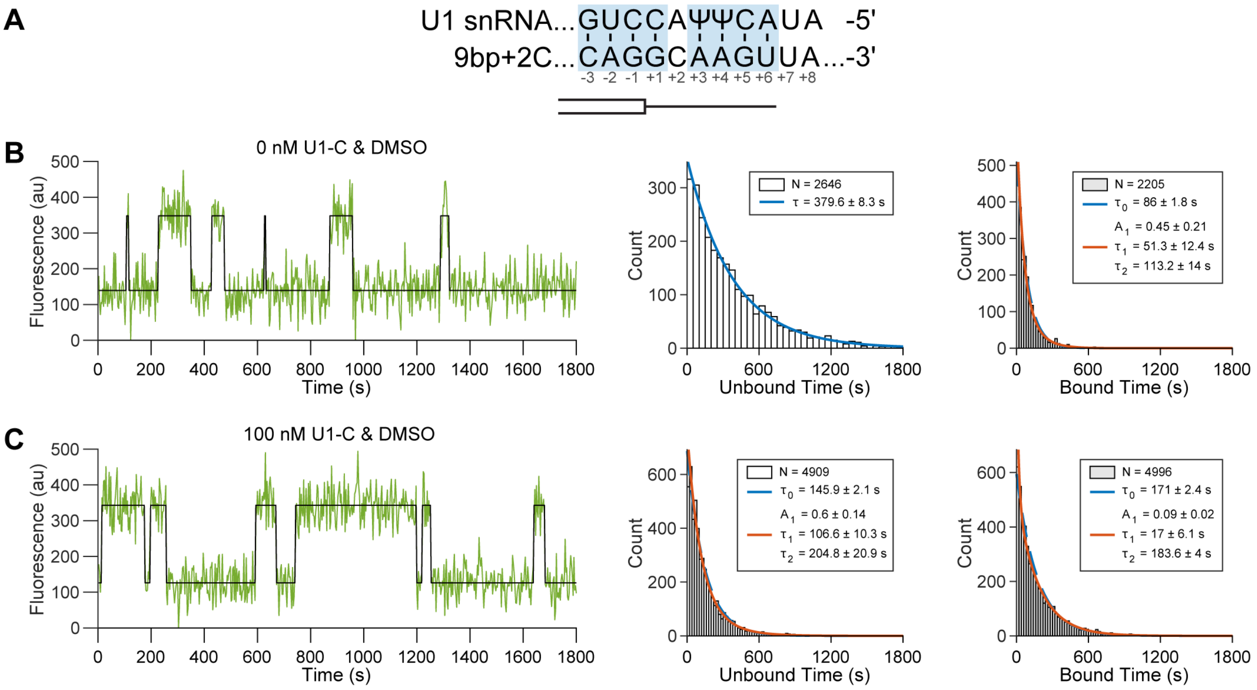


**Figure S23. Effect of U1-C on 9bp+2C RNA oligo binding.** (**A**) Predicted duplex between U1 snRNA and 9bp+2C RNA oligo. (**B-D**) (Left) Representative time series with idealization (black line), (middle) unbound dwell time distributions, and (right) bound dwell time distributions with (100 nM) or without U1-C at 1 nM 9bp+2C RNA oligo. Dwell times are overlaid with expectations from MLE (blue and red lines).

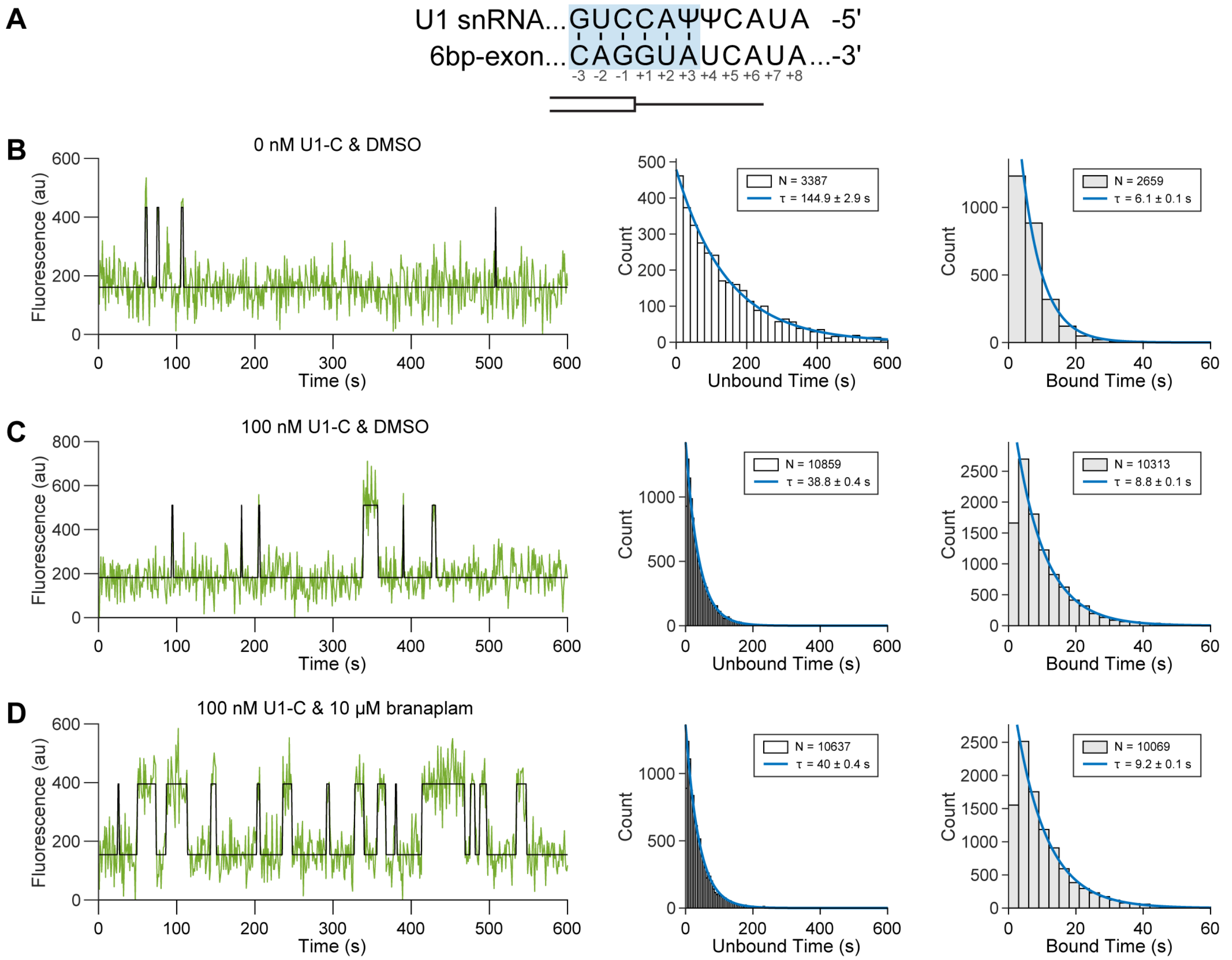


**Figure S24. Effect of U1-C on 6bp-exon RNA oligo binding.** (**A**) Predicted duplex between U1 snRNA SSRS and 6bp-exon RNA oligo. (**B-D**) (Left) Representative time series with idealization (black line), (middle) unbound dwell time distributions, and (right) bound dwell time distributions with (100 nM) or without U1-C at 3 nM 6bp-exon RNA oligo. Dwell times are overlaid with expectations from MLE (blue and red lines).

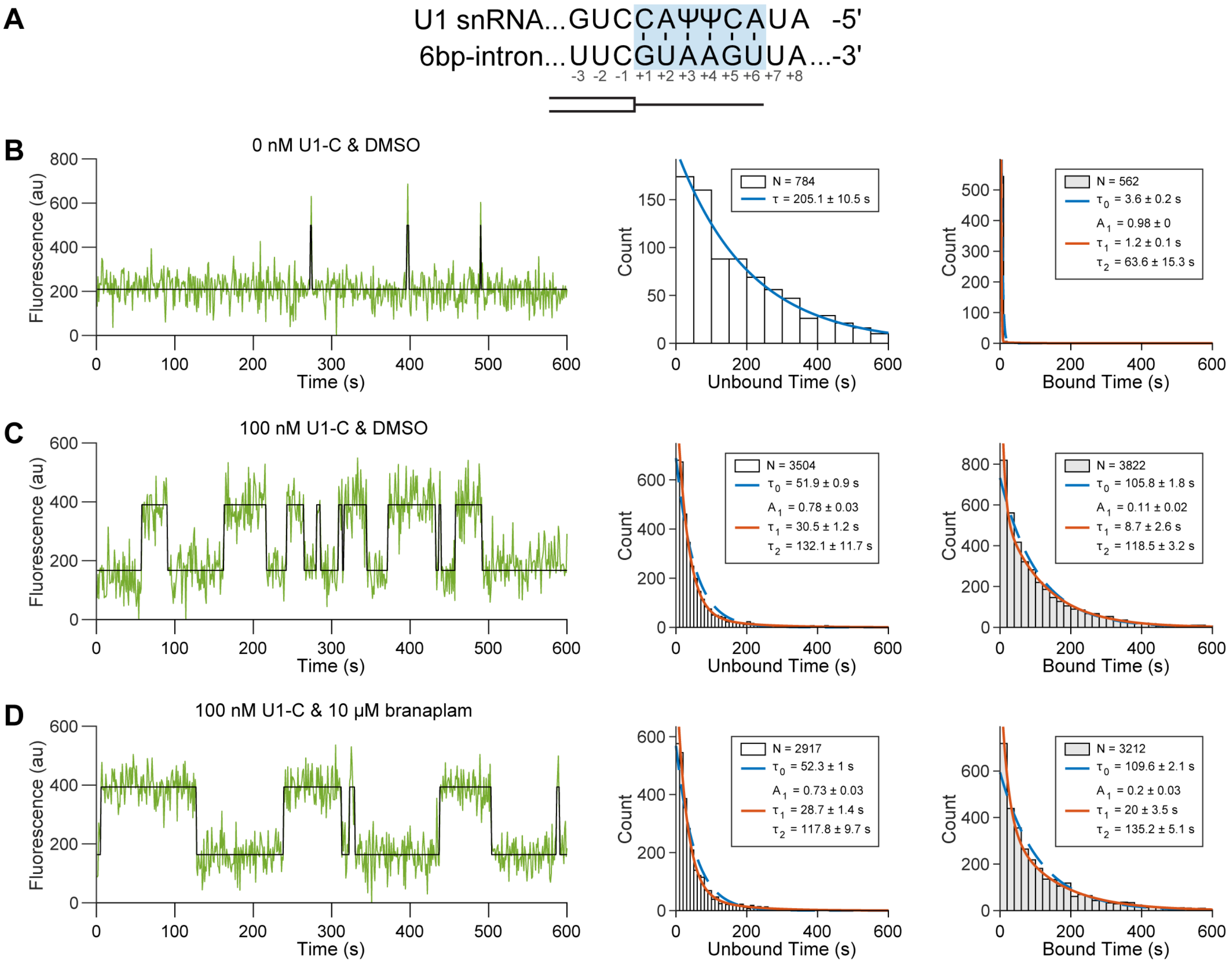


**Figure S25. Effect of U1-C on 6bp-intron RNA binding.** (**A**) Predicted duplex between U1 snRNA SSRS and 6bp-intron RNA oligo. (**B-D**) (Left) Representative time series with idealization (black line), (middle) unbound dwell time distributions, and (right) bound dwell time distributions with (100 nM) or without U1-C at 3 nM 6bp-intron RNA oligo. Dwell times are overlaid with expectations from MLE (blue and red lines).

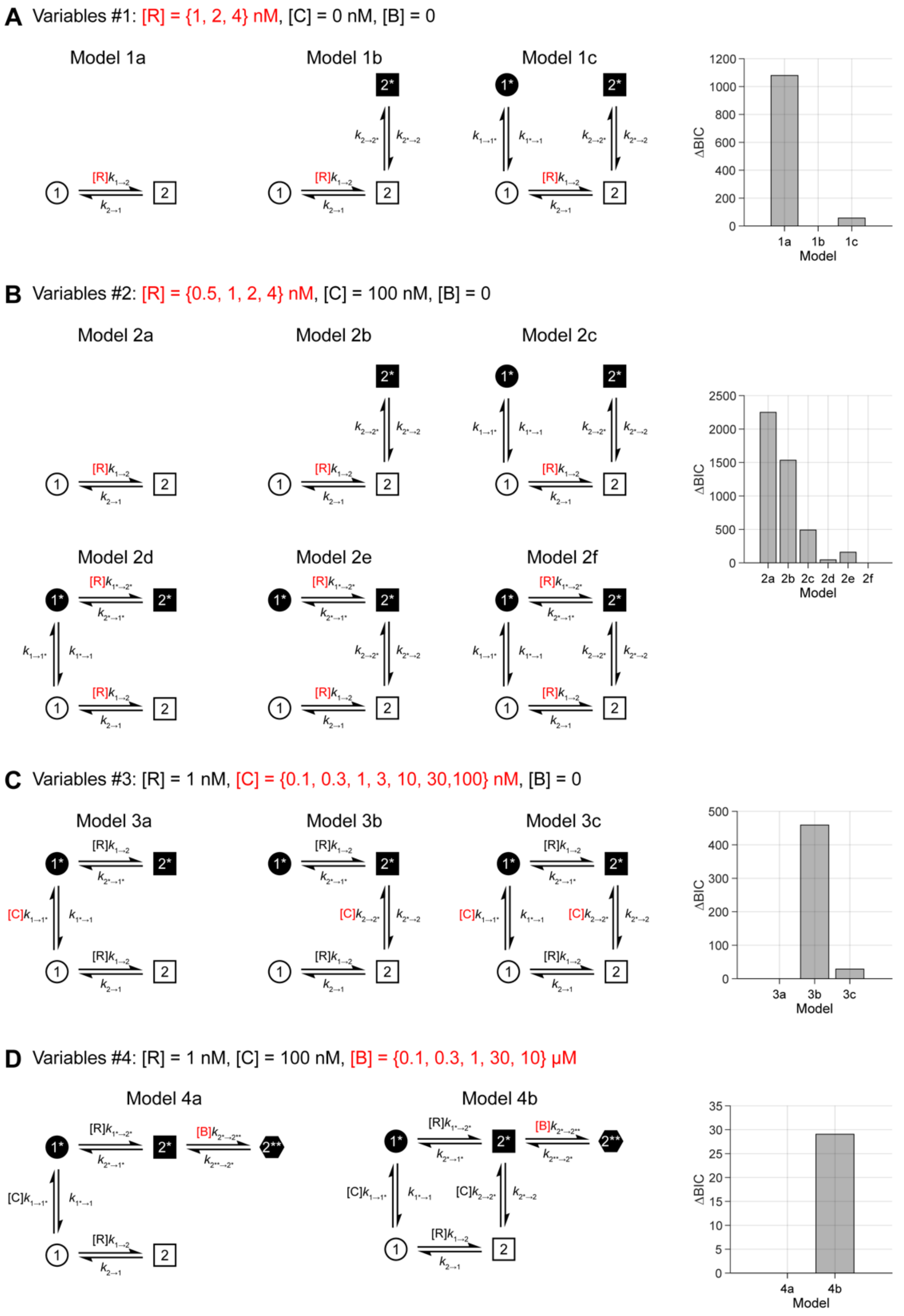


**Figure S26: Kinetic models. (A-D)** Kinetic models evaluated for each single-molecule dataset. R denotes the concentration of 9bp-1A RNA oligo, C denotes the concentration of U1-C, and B denotes the concentration of branaplam. States denoted as 1 indicate 9bp-1A RNA is not bound and as 2 indicate 9bp-1A RNA is bound. Models were ranked according to their BIC score (ΔBIC) where the most probable of the tested models has a ΔBIC score of 0. Optimized rate constants are located in **Table S3.**

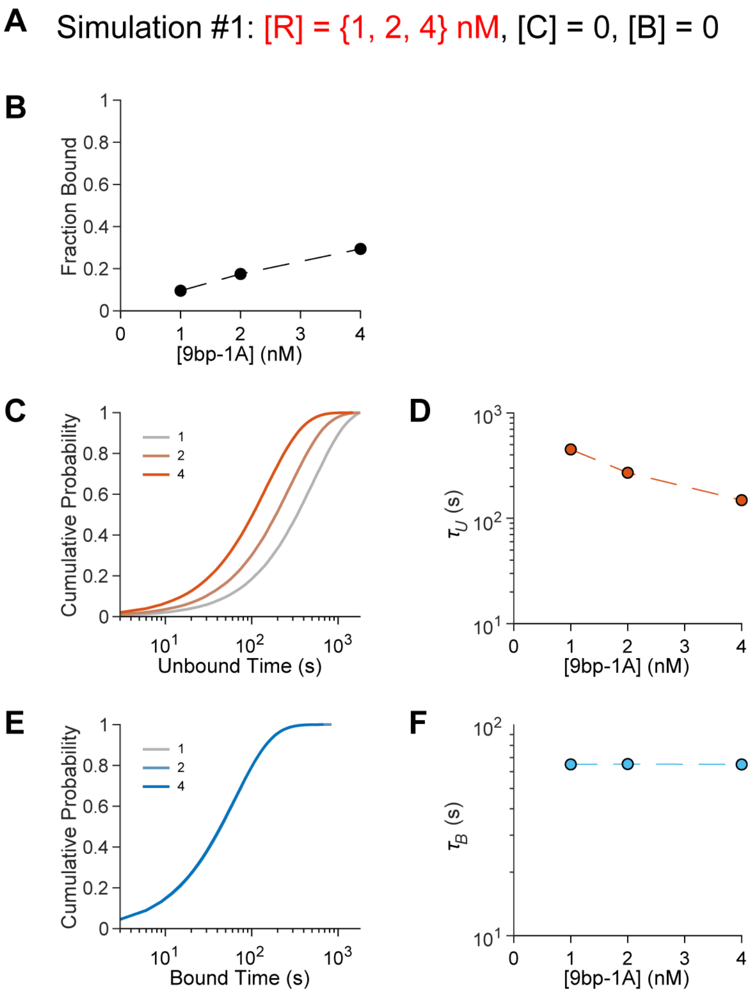


**Figure S27. Simulation of varying 9bp-1A without U1-C and branaplam.** (**A**) Simulation parameters. R denotes the concentration of 9bp-1A RNA oligo, C denotes the concentration of U1-C, and B denotes the concentration of branaplam. For each 9bp-1A concentration, 10,000 molecules were simulated. (**B**) Fraction bound vs 9bp-1A RNA oligo concentration. (**C**). Unbound dwell times across 9bp-1A concentrations. (**D**) Unbound time constants from MLE of single exponential distributions across 9bp-1A RNA oligo concentrations. (**E**) Bound dwell times across 9bp-1A RNA oligo concentrations. (**F**) Bound time constants from MLE of single exponential distributions across 9bp-1A RNA oligo concentrations.

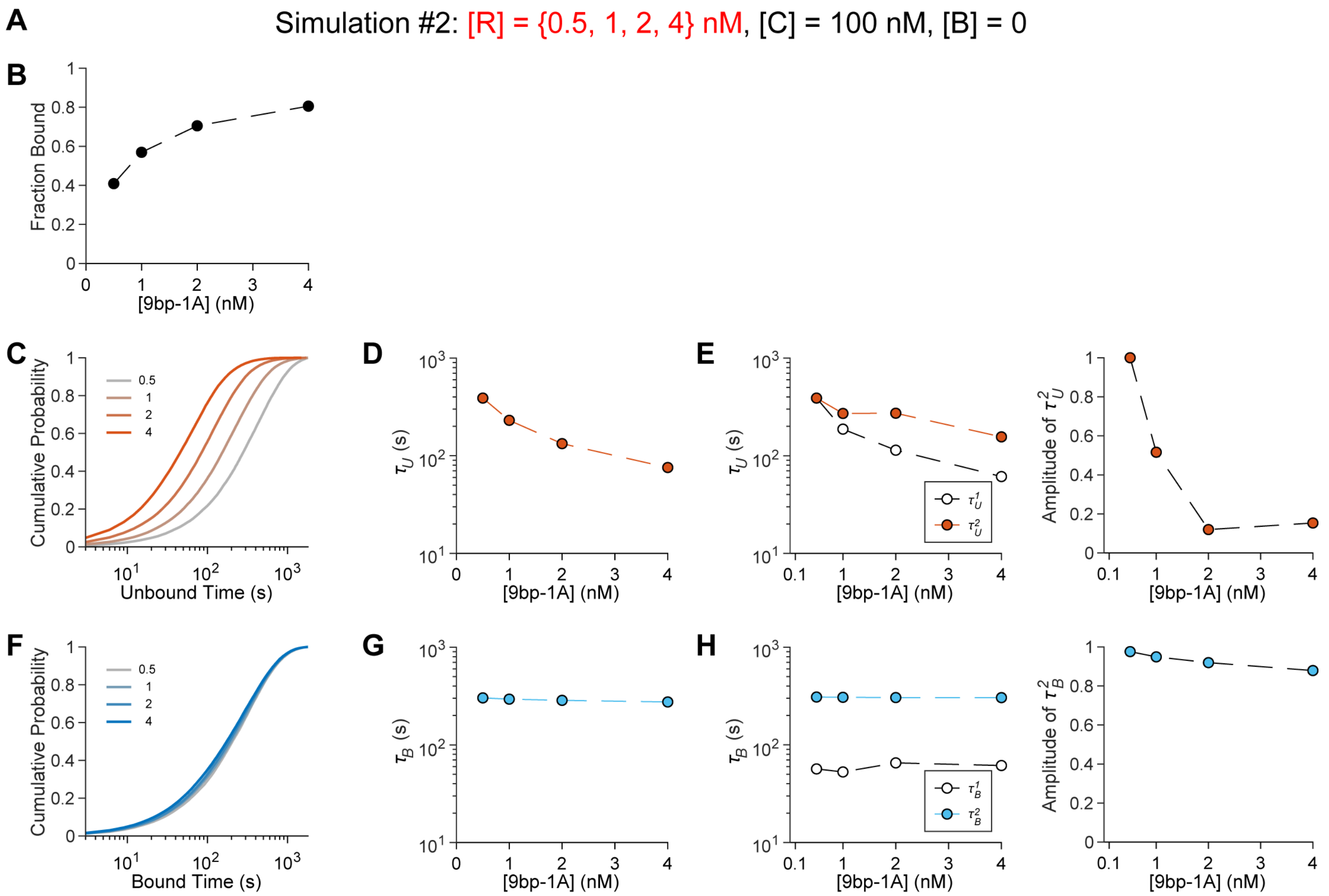


**Figure S28. Simulation of varying 9bp-1A with 100 nM U1-C and no branaplam.** (**A**) Simulation parameters. R denotes the concentration of 9bp-1A RNA oligo, C denotes the concentration of U1-C, and B denotes the concentration of branaplam. For each 9bp-1A concentration, 10,000 molecules were simulated. (**B**) Fraction bound vs 9bp-1A concentration. (**C**). Unbound dwell times across 9bp-1A concentrations. (**D**) Unbound time constants from MLE of single exponential distributions across 9bp-1A concentrations. (**E**) (left) Unbound time constants (right) amplitude of $\tau_U^2$ from MLE of biexponential distributions across 9bp-1A concentrations. (**F**) Bound time constants times across 9bp-1A concentrations. (**G**) Bound time constants from MLE of single exponential distributions across 9bp-1A concentrations. (**H**) (left) Bound time constants (right) amplitude of $\tau_B^2$ from MLE of biexponential distributions across 9bp-1A concentrations.

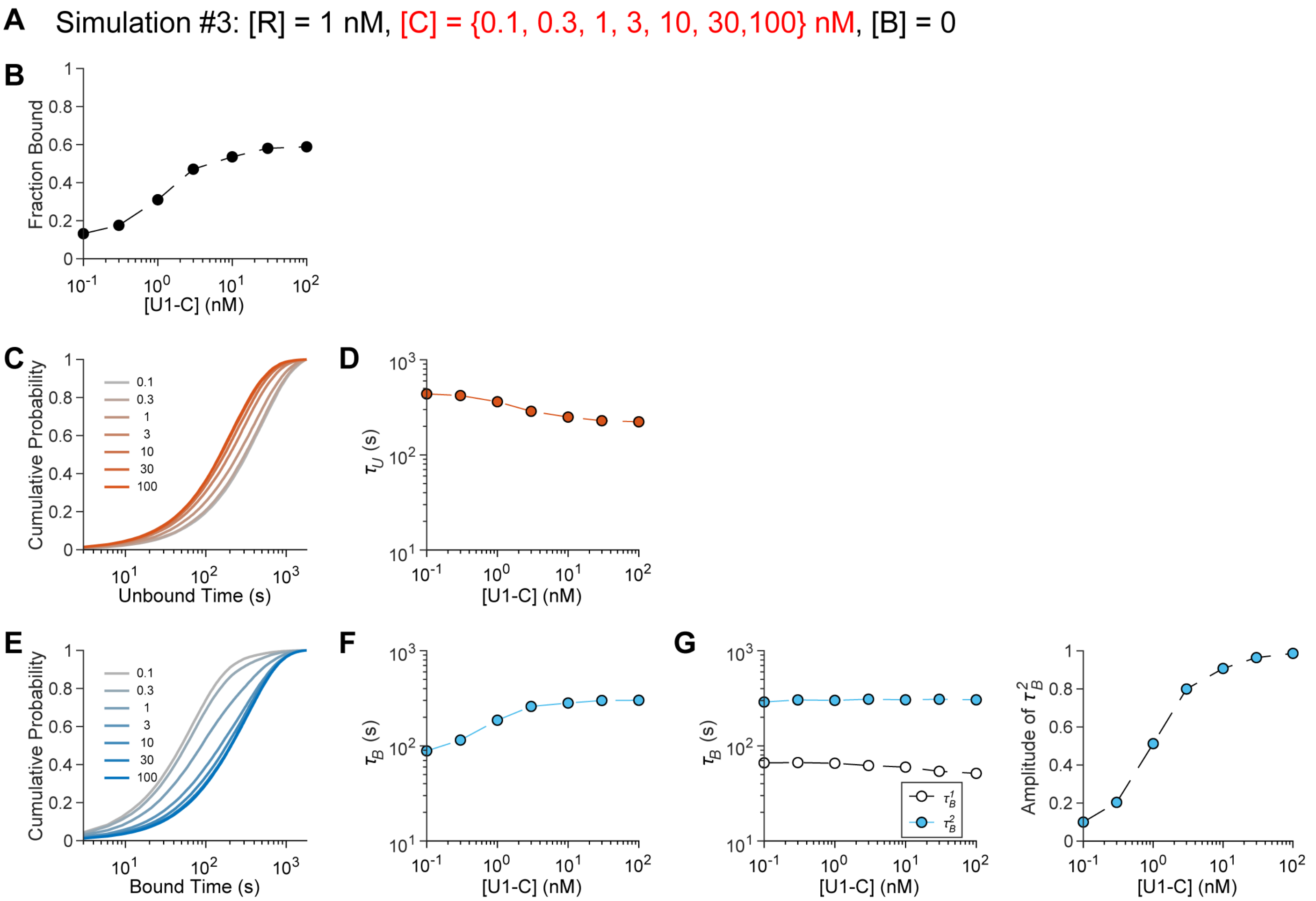


**Figure S29. Simulation of varying U1-C at 1 nM 9bp-1A RNA oligo and no branaplam.** (**A**) Simulation parameters. R denotes the concentration of 9bp-1A RNA oligo, C denotes the concentration of U1-C, and B denotes the concentration of branaplam. For each U1-C concentration, 10,000 molecules were simulated. (**B**) Fraction bound vs 9bp-1A RNA oligo concentration. (**C**). Unbound dwell times across U1-C concentrations. (**D**) Unbound time constants rate from MLE of single exponential distributions across U1-C concentrations. (**E**) Bound dwell times across U1-C concentrations. (**F**) Bound time constants from MLE of single exponential distributions across U1-C concentrations (**G**) (left) Bound time constants (right) amplitude of $\tau_B^2$ from MLE of biexponential distributions across U1-C concentrations.

**A** Simulation #4: [R] = 1 nM, [C] = 100 nM, [B] = {0.1, 0.3, 1, 30, 10} μM

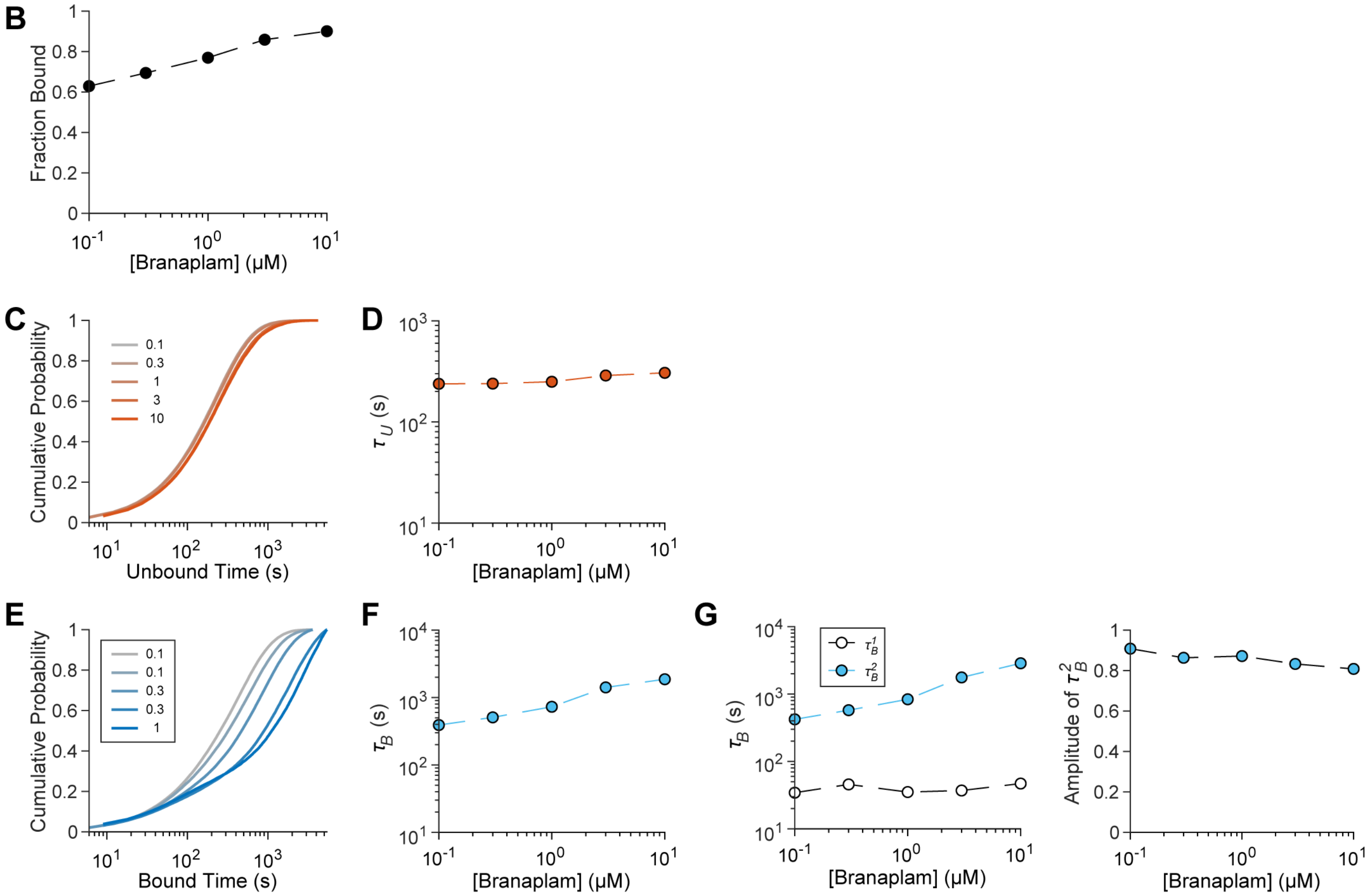


**Figure S30. Simulation of varying branaplam at 1 nM 9bp-1A RNA oligo and 100 nM U1-C.** (**A**) Simulation parameters. R denotes the concentration of 9bp-1A RNA oligo, C denotes the concentration of U1-C, and B denotes the concentration of branaplam. For each branaplam concentration, 10,000 molecules were simulated. (**B**). Fraction bound vs 9bp-1A RNA oligo concentration. (**C**). Unbound dwell times across branaplam concentrations. (**D**) Unbound time constants rate from MLE of single exponential distributions across branaplam concentrations. (**E**) Bound dwell times across branaplam concentrations. (**F**) Bound time constants from MLE of single exponential distributions across branaplam concentrations (**G**) (left) Bound time constants (right) amplitude of $\tau_B^2$ from MLE of biexponential distributions across branaplam concentrations.

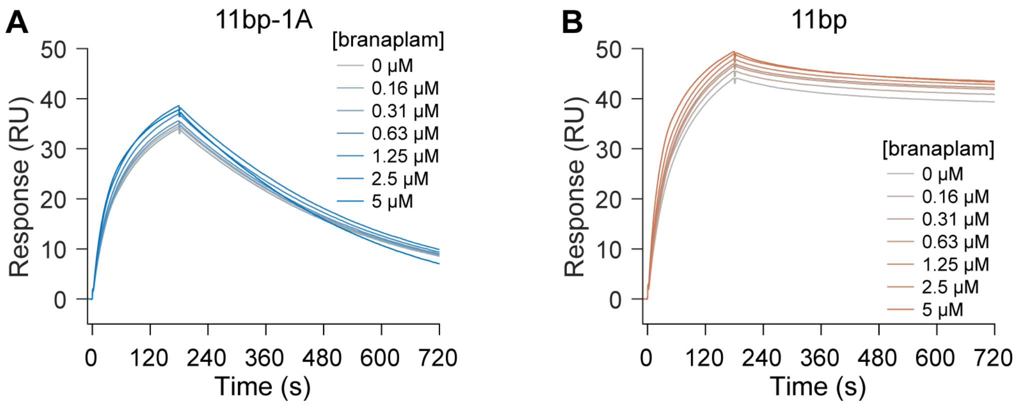


**Figure S31. SPR analysis of the effect of branaplam wash-out on 11bp-1A stability.** Sensorgrams of (**A**) 11bp-1A and (**B**) 11bp at 10nM U1 snRNP across branaplam concentrations. These data were collected under a normal injection mode (as opposed to the co-injection mode of **Figure 2C**) whereby branaplam was washed out of solution prior to recording.

**Table S1. RNA Oligos**

| Name | Assay | RNA Oligo Sequence |
|---|---|---|
| 11bp | SPR | 5'-rCrArGrGrUrArArGrUrArU-LCBi-3' |
| 11bp-1A | SPR | 5'-rCrArGrArGrUrArArGrUrArU-LCBi-3' |
| SMN2 | SPR | 5'-rArGrGrArGrUrArArGrUrCrU-LCBi-3' |
| 11bp | MST | 5'-Cy5-rCrArGrGrUrArArGrUrArU-3' |
| 11bp-1A | MST | 5'-Cy5-rCrArGrArGrUrArArGrUrArU-3' |
| 9bp | CoSMoS | 5'-rArCrArCrArGrGrUrArArGrUrUrArArCrA/3Cy3Sp/-3' |
| 9bp-1A | CoSMoS | 5'-rArCrArCrArGrArGrUrArArGrUrUrArArCrA/3Cy3Sp/-3' |
| 9bp-1C | CoSMoS | 5'-rArCrArCrArGrCrGrUrArArGrUrUrArArCrA/3Cy3Sp/-3' |
| 9bp+1C | CoSMoS | 5'-rArCrArCrArGrCrUrArArGrUrUrArArCrA/3Cy3Sp/-3' |
| 9bp+2C | CoSMoS | 5'-rArCrArCrArGrGrCrArArGrUrUrArArCrA/3Cy3Sp/-3' |
| 6bp_exon | CoSMoS | 5'-rArCrArCrCrArGrGrUrArUrCrArUrArArCrA/3Cy3Sp/-3' |
| 6bp_intron | CoSMoS | 5'-rArCrArCrUrUrCrGrUrArArGrUrUrArArCrA/3Cy3Sp/-3' |
| Control | CoSMoS | 5'-rArCrArCrGrUrCrCrArUrUrCrArUrArArCrA/3Cy3Sp/-3' |
| SMN2 | CoSMoS | 5'-rArCrArArGrGrArGrUrArArGrUrCrUrArCrA/3Cy3Sp/-3' |
| HTT* | CoSMoS | 5'-rArCrArCrArGrArGrUrArArGrGrUrArArCrA/3Cy3Sp/-3' |
| FOXM1 | CoSMoS | 5'-rArCrArArUrGrArGrUrArArGrUrUrCrArCrA/3Cy3Sp/-3' |
| SF3B3 | CoSMoS | 5'-rArCrArUrArGrArGrUrArArGrArCrArArCrA/3Cy3Sp/-3' |
| U1 mimic | CoSMoS | 5'-rArCrArCrGrUrCrCrArUrUrCrArUrArArCrA/3Cy3Sp/-3' |

### Table S2. Summary of CoSMoS Data

| RNA | [RNA] | [U1-C] (nM) | [Branaplam] (μM) | Videos | Total U1 Molecules | Total Frames | Frame Rate (Hz) | Total Time (s) | Total Unbound Events | Total Bound Events |
|---|---|---|---|---|---|---|---|---|---|---|
| 9bp | 0.25 | 100 | 0 | 2 | 1204 | 722400 | 0.33 | 2167200 | 933 | 914 |
| | 0.5 | 0 | 0 | 2 | 999 | 599400 | 0.33 | 1798200 | 826 | 767 |
| | 0.5 | 100 | 0 | 2 | 1283 | 769800 | 0.33 | 2309400 | 1144 | 1118 |
| | 1 | 0 | 0 | 2 | 725 | 435000 | 0.33 | 1305000 | 711 | 658 |
| | 1 | 100 | 0 | 2 | 1271 | 762600 | 0.33 | 2287800 | 1148 | 1101 |
| | 2 | 0 | 0 | 2 | 867 | 520200 | 0.33 | 1560600 | 909 | 863 |
| | 2 | 100 | 0 | 2 | 1156 | 693600 | 0.33 | 2080800 | 987 | 958 |
| | 4 | 0 | 0 | 2 | 774 | 464400 | 0.33 | 1393200 | 783 | 741 |
| 9bp+1C | 10 | 0 | 0 | 2 | 279 | 167400 | 1.00 | 167400 | 1072 | 800 |
| | 10 | 100 | 0 | 2 | 782 | 469200 | 1.00 | 469200 | 9716 | 9125 |
| 9bp+2C | 1 | 0 | 0 | 2 | 650 | 390000 | 0.33 | 1170000 | 2646 | 2205 |
| | 1 | 100 | 0 | 2 | 892 | 535200 | 0.33 | 1605600 | 4909 | 4996 |
| 9bp-1A | 0.5 | 100 | 0 | 2 | 1032 | 619200 | 0.33 | 1857600 | 2670 | 2380 |
| | 1 | 0 | 0 | 2 | 886 | 531600 | 0.33 | 1594800 | 3106 | 2381 |
| | 1 | 0 | 10 | 3 | 1570 | 942000 | 0.33 | 2826000 | 4900 | 3806 |
| | 1 | 0.1 | 0 | 2 | 821 | 492600 | 0.33 | 1477800 | 2920 | 2247 |
| | 1 | 0.3 | 0 | 2 | 958 | 574800 | 0.33 | 1724400 | 3302 | 2590 |
| | 1 | 1 | 0 | 3 | 1577 | 946200 | 0.33 | 2838600 | 5543 | 4764 |
| | 1 | 3 | 0 | 2 | 932 | 559200 | 0.33 | 1677600 | 3035 | 2793 |
| | 1 | 10 | 0 | 2 | 986 | 591600 | 0.33 | 1774800 | 3169 | 3105 |
| | 1 | 30 | 0 | 2 | 872 | 523200 | 0.33 | 1569600 | 2719 | 2749 |
| | 1 | 100 | 0 | 3 | 1682 | 1009200 | 0.33 | 3027600 | 5154 | 5216 |
| | 1 | 100 | 0.1 | 2 | 1194 | 716400 | 0.17 | 4298400 | 5959 | 6234 |
| | 1 | 100 | 0.3 | 2 | 1152 | 691200 | 0.17 | 4147200 | 4463 | 4811 |
| | 1 | 100 | 1 | 2 | 1268 | 760800 | 0.17 | 4564800 | 3103 | 3640 |
| | 1 | 100 | 3 | 2 | 978 | 586800 | 0.11 | 5281200 | 1765 | 2211 |
| | 1 | 100 | 10 | 2 | 1110 | 666000 | 0.11 | 5994000 | 1156 | 1503 |
| | 2 | 0 | 0 | 2 | 959 | 575400 | 0.33 | 1726200 | 5389 | 4789 |
| | 2 | 100 | 0 | 2 | 1000 | 600000 | 0.33 | 1800000 | 3788 | 4093 |
| | 4 | 0 | 0 | 2 | 901 | 540600 | 0.33 | 1621800 | 7344 | 6911 |
| | 4 | 100 | 0 | 2 | 966 | 579600 | 0.33 | 1738800 | 4373 | 4895 |
| 9bp-1C | 1 | 0 | 0 | 2 | 828 | 496800 | 0.33 | 1490400 | 2546 | 1805 |
| | 1 | 100 | 0 | 2 | 1144 | 686400 | 0.33 | 2059200 | 3597 | 3682 |
| | 1 | 100 | 10 | 2 | 1003 | 601800 | 0.33 | 1805400 | 3172 | 3311 |
| FOXM1 | 10 | 0 | 0 | 2 | 241 | 144600 | 1.00 | 144600 | 936 | 703 |
| | 10 | 0 | 10 | 2 | 197 | 118200 | 1.00 | 118200 | 700 | 514 |
| | 10 | 100 | 0 | 2 | 620 | 372000 | 1.00 | 372000 | 7418 | 7136 |
| | 10 | 100 | 10 | 2 | 649 | 389400 | 1.00 | 389400 | 5099 | 5097 |
| HTT* | 3 | 0 | 0 | 2 | 863 | 517800 | 1.00 | 517800 | 4869 | 4047 |
| | 3 | 0 | 10 | 2 | 943 | 565800 | 1.00 | 565800 | 5394 | 4512 |
| | 3 | 100 | 0 | 2 | 917 | 550200 | 1.00 | 550200 | 13428 | 13192 |
| | 3 | 100 | 10 | 2 | 785 | 471000 | 0.33 | 1413000 | 7766 | 8201 |
| SF3B3 | 10 | 0 | 0 | 2 | 22 | 13200 | 1.00 | 13200 | 61 | 43 |
| | 10 | 0 | 10 | 2 | 18 | 10800 | 1.00 | 10800 | 40 | 23 |
| | 10 | 100 | 0 | 2 | 627 | 376200 | 1.00 | 376200 | 11301 | 10844 |
| | 10 | 100 | 10 | 2 | 576 | 345600 | 1.00 | 345600 | 5074 | 5160 |
| SMN2 | 10 | 0 | 0 | 2 | 289 | 173400 | 1.00 | 173400 | 877 | 597 |
| | 10 | 0 | 10 | 2 | 83 | 49800 | 1.00 | 49800 | 254 | 173 |
| | 10 | 100 | 0 | 2 | 901 | 540600 | 1.00 | 540600 | 8646 | 8381 |
| | 10 | 100 | 10 | 2 | 978 | 586800 | 1.00 | 586800 | 7056 | 7005 |
| 6bp-exon | 3 | 0 | 0 | 2 | 230 | 138000 | 1.00 | 138000 | 784 | 562 |
| | 3 | 100 | 0 | 2 | 990 | 594000 | 1.00 | 594000 | 3504 | 3822 |
| | 3 | 100 | 10 | 2 | 887 | 532200 | 1.00 | 532200 | 2917 | 3212 |
| 6bp-intron | 3 | 0 | 0 | 2 | 800 | 480000 | 1.00 | 480000 | 3387 | 2659 |
| | 3 | 100 | 0 | 2 | 893 | 535800 | 1.00 | 535800 | 10859 | 10313 |
| | 3 | 100 | 10 | 2 | 899 | 539400 | 1.00 | 539400 | 10637 | 10069 |
| U1 Mimic | 10 | 100 | 0 | 2 | 736 | 441600 | 0.33 | 1324800 | 14 | 8 |

### Table S3. Optimized rate constants of kinetic models

| Model* | | | | | | | | | | |
|---|---|---|---|---|---|---|---|---|---|---|
| **1a** | 1.56 ± 0.01 x $10^{6}$ | 1.53 ± 0.01 x $10^{-2}$ | - | - | - | - | - | - | - | - |
| **1b** | 1.57 ± 0.01 x $10^{6}$ | 1.79 ± 0.02 x $10^{-2}$ | - | - | - | - | 3.97 ± 0.32 x $10^{-3}$ | 6.62 ± 0.78 x $10^{-4}$ | - | - |
| **1c** | 1.20 ± 0.57 x $10^{6}$ | 1.79 ± 0.02 x $10^{-2}$ | 2.26 ± 2.35 | 8.96 ± 9.03 | - | - | 3.97 ± 0.35 x $10^{-3}$ | 6.62 ± 0.83 x $10^{-4}$ | - | - |
| **2a** | 3.09 ± 0.03 x $10^{6}$ | 2.93 ± 0.02 x $10^{-3}$ | - | - | - | - | - | - | - | - |
| **2b** | 3.10 ± 0.02 x $10^{6}$ | 5.67 ± 0.19 x $10^{-3}$ | - | - | - | - | 9.13 ± 1.08 x $10^{-3}$ | 9.85 ± 0.65 x $10^{-3}$ | - | - |
| **2c** | 4.14 ± 0.06 x $10^{6}$ | 5.70 ± 0.19 x $10^{-3}$ | 1.29 ± 0.11 x $10^{-3}$ | 4.19 ± 0.23 x $10^{-3}$ | - | - | 9.12 ± 1.07 x $10^{-3}$ | 9.85 ± 0.66 x $10^{-3}$ | - | - |
| **2d** | 1.18 ± 0.04 x $10^{6}$ | 1.63 ± 0.08 x $10^{-2}$ | 1.41 ± 0.10 x $10^{-3}$ | 7.87 ± 0.74 x $10^{-4}$ | 4.29 ± 0.06 x $10^{6}$ | 2.59 ± 0.03 x $10^{-3}$ | - | - | - | - |
| **2e** | 1.67 ± 0.04 x $10^{6}$ | 1.09 ± 0.05 x $10^{-2}$ | - | - | 4.35 ± 0.07 x $10^{6}$ | 2.38 ± 0.03 x $10^{-3}$ | 4.56 ± 0.46 x $10^{-3}$ | 3.08 ± 0.42 x $10^{-4}$ | - | - |
| **2f** | 1.22 ± 0.04 x $10^{6}$ | 1.56 ± 0.10 x $10^{-2}$ | 9.93 ± 1.75 x $10^{-4}$ | 3.08 ± 0.97 x $10^{-4}$ | 4.44 ± 0.07 x $10^{6}$ | 2.44 ± 0.04 x $10^{-3}$ | 1.08 ± 0.75 x $10^{-3}$ | 1.74 ± 0.41 x $10^{-4}$ | - | - |
| **3a** | 1.68 ± 0.02 x $10^{6}$ | 1.87 ± 0.03 x $10^{-2}$ | 2.43 ± 0.30 x $10^{3}$ | 7.69 ± 0.91 x $10^{-3}$ | 3.37 ± 0.04 x $10^{6}$ | 2.71 ± 0.03 x $10^{-3}$ | - | - | - | - |
| **3b** | 1.81 ± 0.02 x $10^{6}$ | 1.63 ± 0.03 x $10^{-2}$ | - | - | 3.31 ± 0.04 x $10^{6}$ | 2.52 ± 0.03 x $10^{-3}$ | 2.96 ± 00.21 x $10^{6}$ | 1.05 ± 0.07 x $10^{-3}$ | - | - |
| **3c** | 1.68 ± 0.02 x $10^{6}$ | 1.87 ± 0.03 x $10^{-2}$ | 2.43 ± 0.28 x 103 | 7.69 ± 0.83 x $10^{-3}$ | 3.37 ± 0.04 x $10^{6}$ | 2.71 ± 0.03 x $10^{-3}$ | 0.01 ± 1.19 | 0.02 ± 1.99 x $10^{-8}$ | - | - |
| **4a** | 1.05 ± 0.03 x $10^{6}$ | 1.16 ± 0.05 x $10^{-2}$ | 4.66 ± 0.25 x $10^{3}$ | 2.44 ± 0.19 x $10^{-3}$ | 4.24 ± 0.05 x $10^{6}$ | 2.21 ± 0.02 x $10^{-3}$ | - | - | 1.207 ± 0.01 x $10^{4}$ | 8.40 ± 0.24 x $10^{-3}$ |
| **4b** | 1.05 ± 0.02 x $10^{6}$ | 1.16 ± 0.04 x $10^{-2}$ | 4.66 ± 0.25 x $10^{3}$ | 2.44 ± 0.16 x $10^{-4}$ | 4.24 ± 0.05 x $10^{6}$ | 2.21 ± 0.02 x $10^{-3}$ | 1.76 ± 1.76 x $10^{-5}$ | 1.07 ± 1.08 x $10^{-12}$ | 1.27 ± 0.03 x $10^{4}$ | 8.40 ± 0.11 x $10^{-3}$ |

*Units are $M^{-1}s^{-1}$ or $s^{-1}$ depending on the model. See **Figure S26.**
